# Discovery, Synthesis, and Optimization of 1,2,4-Triazolyl Pyridines Targeting *Mycobacterium tuberculosis*

**DOI:** 10.1101/2022.11.14.516356

**Authors:** Tomayo Berida, Samuel R. McKee, Shamba Chatterjee, Wei Li, Pankaj Pandey, Siddharth Kaushal Tripathi, Robert J. Doerksen, Mary Jackson, Christian Ducho, Christina L. Stallings, Sudeshna Roy

## Abstract

Tuberculosis (TB) results in 1.5 million deaths every year. The rise in multi-drug resistant TB underscores the urgent need to develop new antibacterials, particularly those with new chemical entities and/or novel mechanisms of action that can be used in combination therapy with existing drugs to prevent the rapid emergence of resistance. Herein, we report the discovery and synthesis of a new series of compounds containing a 3-thio-1,2,4-triazole moiety that show inhibition of *Mycobacterium tuberculosis* (*Mtb*) growth and survival. Structure-activity relationship studies led us to identify potent analogs displaying nanomolar inhibitor activity, specifically against *Mtb*. These potent analogs exhibit a promising ADME/pharmacokinetic profile and no cytotoxicity in mammalian cells at over 100 times the effective dose in *Mtb*. Our preliminary investigations into the mechanism of action suggest this series is not engaging promiscuous targets and, thereby, could be acting on a novel target.

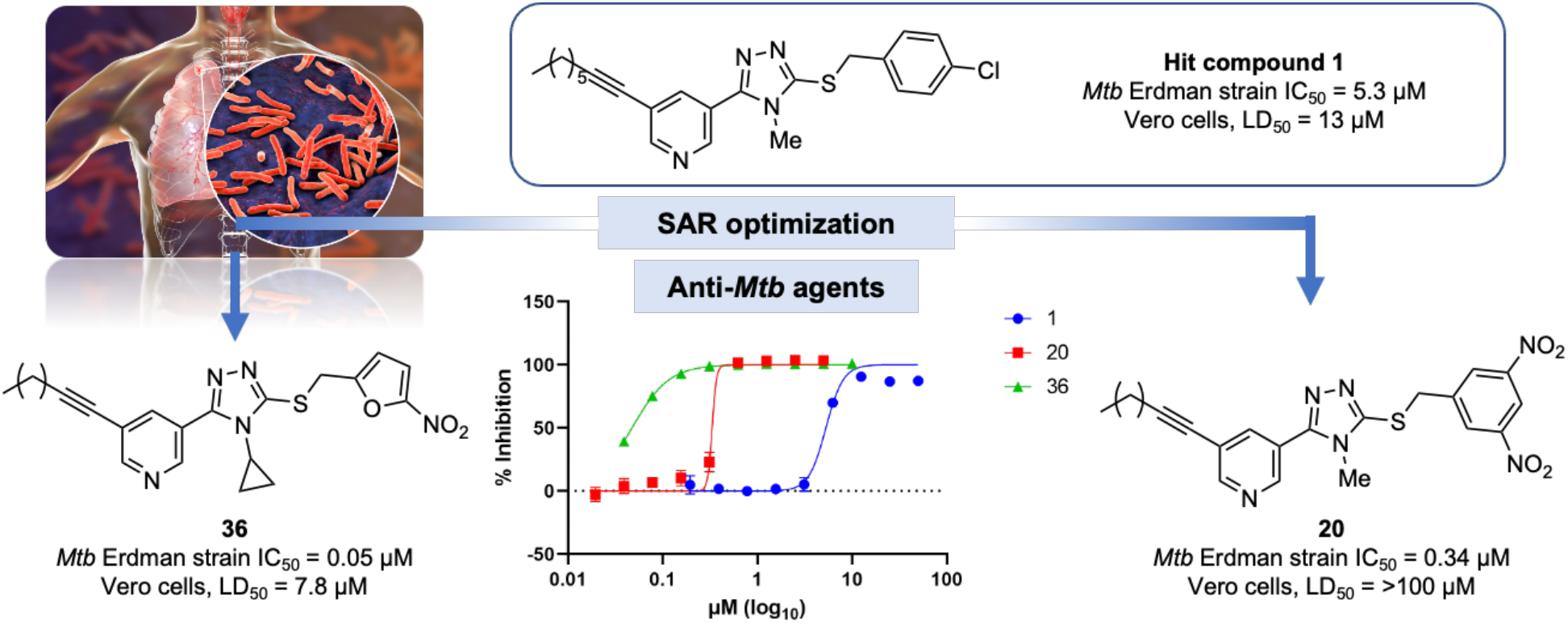

## 1. INTRODUCTION

Tuberculosis (TB) is an airborne infectious disease caused by the pathogen *Mycobacterium tuberculosis* (*Mtb*). It is responsible for ~1.5 million deaths yearly, making it a major global health threat.^1^ TB is curable using a combination therapy of four first-line agents: rifampicin (RIF), isoniazid (INH), ethambutol (EMB), and pyrazinamide (PZA). However, a successful cure of TB requires a long duration of treatment, which can result in poor patient adherence and allow for the selection of drug-resistant mutants.^2^ The emergence of multidrug-resistant TB (MDR-TB) and extensively drug-resistant TB (XDR-TB) pose a significant threat to the control of the disease.^3^ MDR-TB is caused by strains of *Mtb* that are not responsive to INH and RIF (Figure 1).^3, 4^ In 2019, >200,000 MDR-TB or rifampicin-resistant (MDR/RR-TB) infections were detected worldwide, amounting to a 10% increase from 2018.^5^ Treatment of MDR/RR-TB requires the addition of second-line drugs, including a fluoroquinolone along with one of the injectable aminoglycosides, such as amikacin, streptomycin, or kanamycin. Regrettably, this treatment regimen is ineffective against XDR-TB, which is caused by *Mtb* strains resistant to RIF and INH plus second-line agents.

The latest TB drug, pretomanid, was approved by the Food and Drug Administration (FDA) as part of a combination regimen with bedaquiline and linezolid. However, the approval is only for limited cases of adults suffering from XDR-TB and treatment-intolerant or non-responsive MDR pulmonary TB (Figure 1).^6^ Moreover, adverse effects such as hepatotoxicity, myelosuppression, and peripheral neuropathy associated with this combination or other pretomanid-containing regimens are a source of concern.^6, 7^ Another challenge with this combination is the reported resistance to bedaquiline and linezolid.^8^ In fact, the World Health Organization (WHO) recently redefined XDR-TB as being caused by *Mtb* strains that meet the criteria for MDR-TB and are resistant to a fluoroquinolone and either bedaquiline or linezolid or both.^9, 10^ The challenges with drug-resistant TB infections present an urgent need for new anti-*Mtb* drugs, particularly those with new chemical entities and novel mechanisms of action to combat the rise in MDR-TB and XDR-TB. To address this need, we report herein the discovery and development of a novel series of compounds with potent anti-*Mtb* activity and limited cytotoxicity in mammalian cells, which provide the foundation for a new therapeutic strategy for TB.

**Figure 1.**
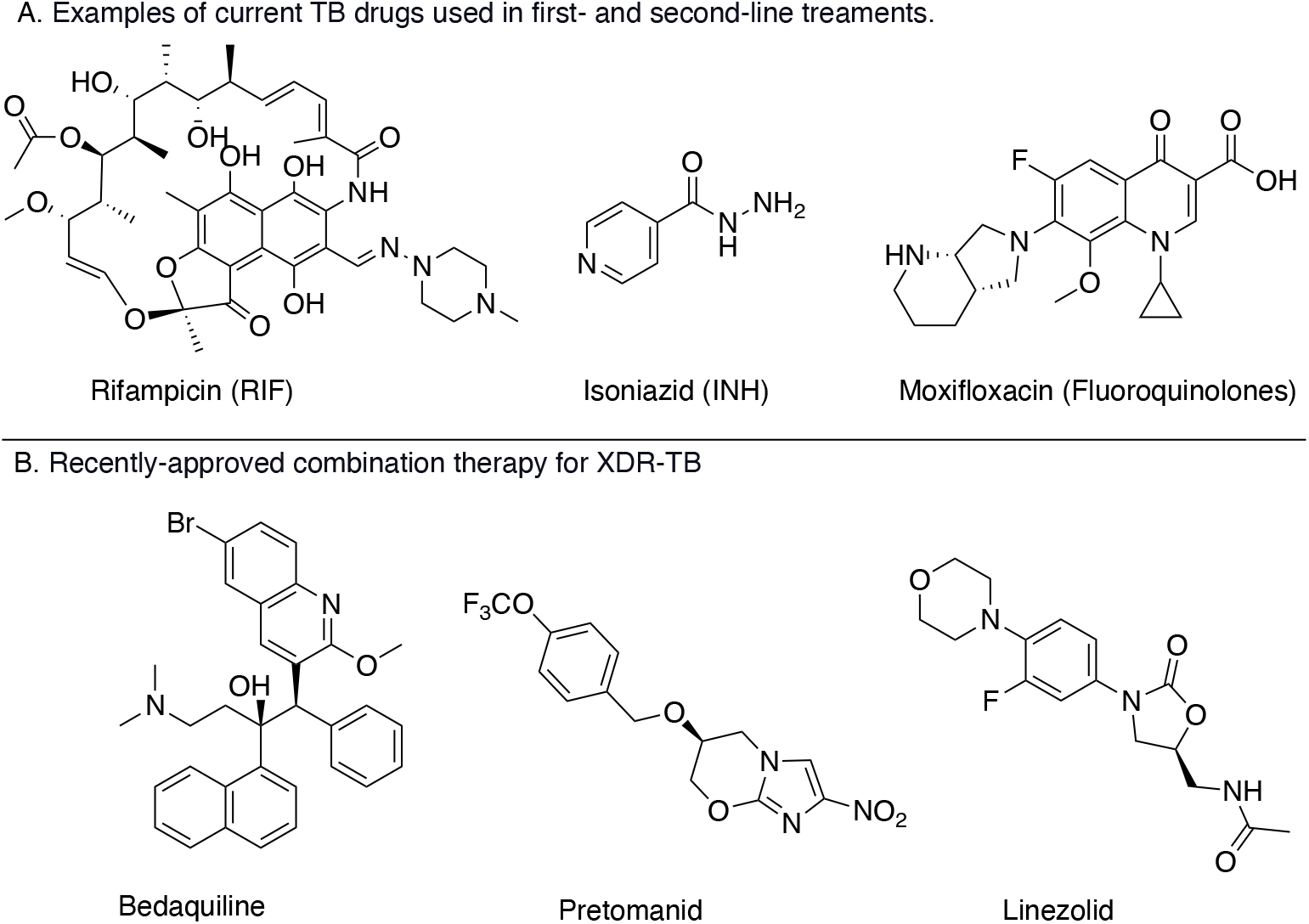
Drugs used for TB treatment.

## 2. RESULTS AND DISCUSSION

### 2.1. Hit discovery

As a part of our ongoing efforts to identify novel anti-*Mtb* agents, we carried out a phenotypic screen for compounds that exhibited *in vitro* anti-*Mtb* properties against the wild-type (WT) *Mtb* Erdman strain using the microplate Alamar Blue assay (MABA). The MABA is commonly used to evaluate the efficacy of compounds for restraining *Mtb* growth.^11^ The assay utilizes the dye resazurin, which is dark blue and non-fluorescent in its oxidized form but becomes pink and fluorescent when reduced to resorufin as a result of cellular metabolism.^11, 12^, The degree of this color change is quantified, and compounds that inhibit *Mtb* growth or survival will decrease or block this color change. By performing this assay with a range of concentrations of each compound, we calculated IC_50_ values for active compounds, which is defined as the concentration of antibiotic that inhibits mycobacterial survival by 50%. These efforts led to the discovery of the hit compound **1**, 3-(5-((4-chlorobenzyl)thio)-4-methyl-4*H*-1,2,4-triazol-3-yl)-5-(oct-1-yn-1-yl)pyridine. The hit **1** exhibits growth inhibition against *Mtb* with an IC_50_ = 5.3 μM in the MABA (Figure 2) and mammalian cytotoxicity against VERO cells *in vitro* with an LD_50_ = 13 μM, where LD_50_ is defined as the lethal dose (or concentration) of antibiotic that inhibits Vero cells viability by 50%. Hence, we deemed this series worthwhile to pursue a hit-to-lead optimization campaign using a traditional medicinal chemistry approach. Here, we outline our structure-activity-relationship (SAR) efforts and the synthesis of the analogs that resulted in the identification of novel anti-*Mtb* agents with nanomolar potency. We also outline our preliminary experiments on the mechanism of action studies, which suggest that this series does not inhibit common promiscuous targets in *Mtb*, such as Mycobacterial membrane protein Large 3 (MmpL3) and QcrB a component of the cytochrome bc1 complex in the electron transport chain, and thereby could be acting on a novel target.^13–15^

**Figure 2.**
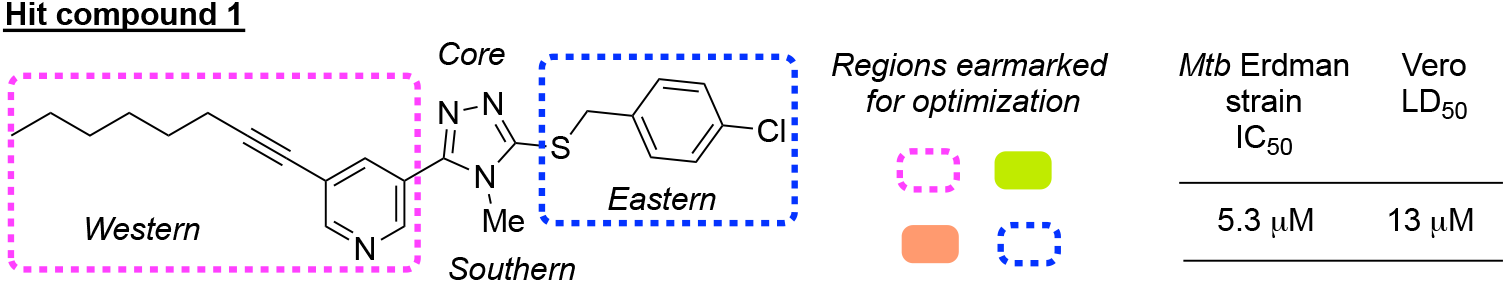
Regions for optimization in hit compound **1**.

### 2.2. Chemistry

#### Synthesis of hit compound 1 and analogs with variations in the eastern and southern regions (Scheme 1)

The synthesis of hit compound **1** and its analogs (**2**–**36**) were assembled via the corresponding 1,2,4-triazole-3-thiones (**55**, **56**–**60**) intermediates that were derived from the hydrazide **54**. The hydrazide **54** was synthesized from commercially available methyl-5-bromonicotinate in two steps (SI Scheme S1) in 86% overall yield. The 1,4-disubstituted thiosemicarbazide intermediates (not shown) obtained from heating the hydrazide **54** under reflux conditions in ethanol with various substituted isothiocyanates were used for the next step without further purification. The 1,4-disubstituted thiosemicarbazides were then treated with 10% aqueous NaOH at 60 °C for 4 h to generate the 1,2,4-triazole-3-thiones through an intramolecular cyclization reaction (**55**, **56**–**60**) in 70–99% yield. Subsequent alkylation of the 1,2,4-triazole-3-thiones with various benzyl (or alkyl) halides was performed to obtain the target compounds (**1**– **22**, **24**–**36**).

**Scheme 1.**
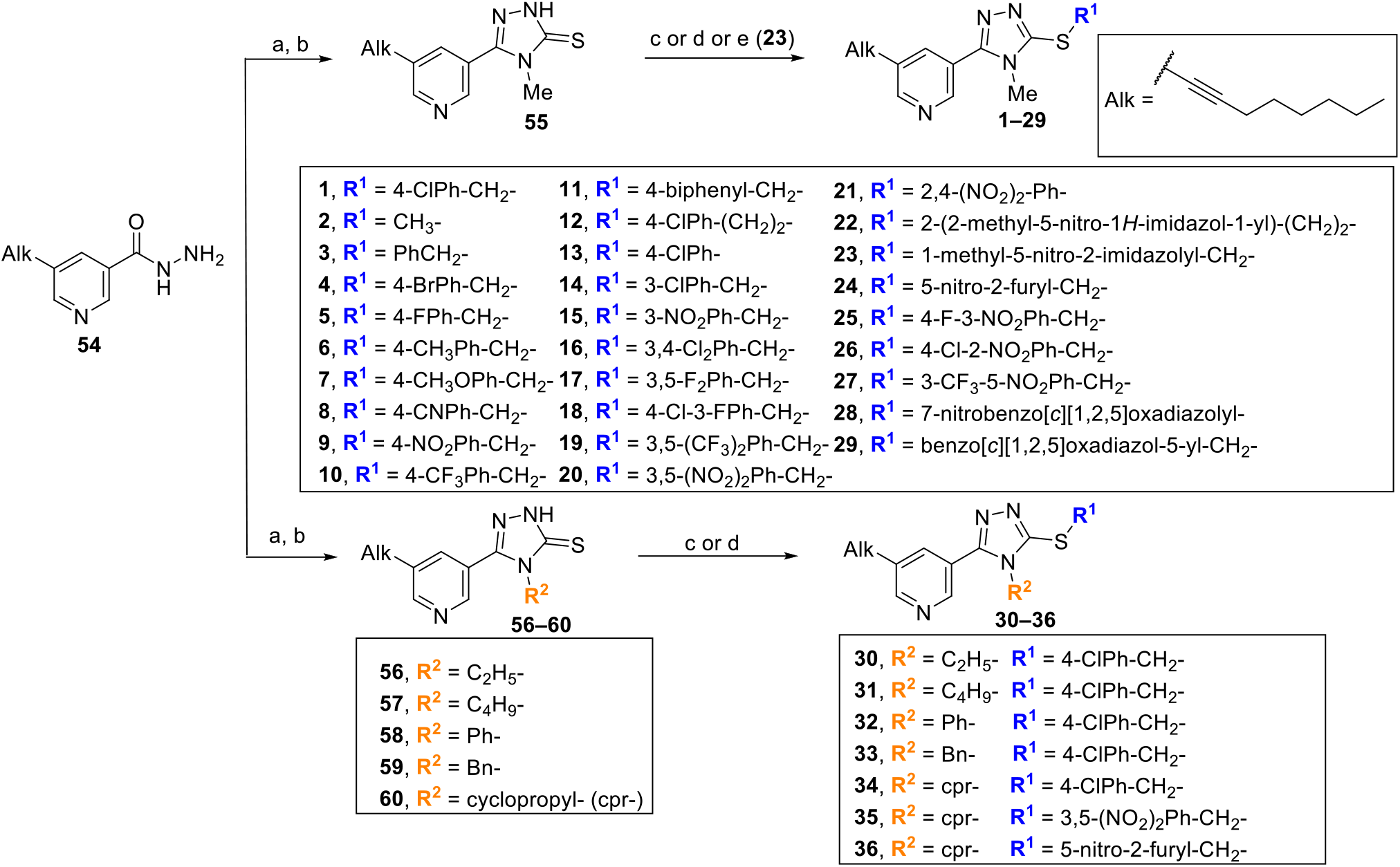
Synthesis of analogs with variations in the eastern and southern regions on hit **1**.^[a]^ ^[a]^ Reagents and conditions: (a) alkyl or aryl isothiocyanates, EtOH, reflux, 4 h; (b) 10% aq. NaOH, 60 °C, 3 h, 70–99% (over two steps); (c) corresponding aryl or alkyl halide, K_2_CO_3_, acetone/MeOH, rt, overnight, 43–87%; (d) aryl methyl halide, Et_3_N, CH_3_CN, 1–24 h, 25–87%; (e) for the synthesis of **23**, 1-methyl-5-nitroimidazole-2-methanol, DIAD, PPh_3_, THF, 0 °C, 30 min, then rt, 3 h, 62%.

Alternatively, the Mitsunobu reaction was used for the alkylation of **55** to generate compound **23** (Scheme 1, condition (e)), taking advantage of the acidity of 5-membered heteroaromatic thiol.^16^ The analog **23** was therefore synthesized under Mitsunobu conditions using 1-methyl-5-nitroimidazole-2-methanol. Next, the analogs **12**, **13**, **21**, and **28** were generated by varying linker lengths between the *S*-atom and eastern aromatic ring. For the two-carbon linker analog **12**, compound **55** was reacted with 4-chlorophenethyl bromide via an S_N_2 reaction. For the analogs (**13**, **21**, and **28**) with the *S*-atom directly connected to the eastern aromatic ring, a palladium-catalyzed coupling reaction between 1-bromo-4-chlorobenzene and **55** was utilized to produce the analog **13**, whereas a nucleophilic aromatic substitution (S_N_Ar) reaction was employed for the analogs **21** and **28**. This chemistry was feasible due to the electron-withdrawing nitro (NO_2_) groups at the ortho or para positions relative to the halide leaving groups on the aromatic scaffold.

#### Synthesis of analogs with variations in the western region (Scheme 2)

The syntheses of compounds **37–38** and **40–43** were achieved in three steps starting from the hydrazides (**61i**–**vi**), which were prepared from commercially available esters in two steps (SI Scheme S2–S3). The hydrazides (**61i**–**vi**) were heated in ethanol with methyl isothiocyanates to obtain the 1,4-disubstituted thiosemicarbazides, which were subsequently treated with 10% aqueous NaOH at 60 °C for 4 h to furnish the 1,2,4-triazole-3-thiones (**62 i**–**vi**) in 42–97% yield (over two steps). Finally, compounds **37–38** and **40–43** were obtained in moderate to excellent yields (61–92%) by reacting 4-chlorobenzyl bromide with the corresponding thiones (**62 i**–**vi**).

Initially, we encountered a challenge in the synthesis of compound **39** when we attempted the same synthetic route used for the series **37–38** and **40–43**. We could not form the 1,2,4-triazole-3-thione in the presence of the terminal acetylene group. This might be due to the interference of the acetylene functionality with the cyclization of the 1,4-disubstituted thiosemicarbazide precursor. Moreover, trimethylsilyl (TMS) protection of the acetylene could not solve this problem since the TMS group was labile under cyclization conditions (Scheme 2). Analog **39** was then synthesized in 91% yield by coupling compound **38** with TMS acetylene, followed by deprotection with K_2_CO_3_ in MeOH. The analog **45** was synthesized from the commercially available carboxylic acid **64**, which was coupled with 4-methyl-3-thiosemicarbazide using EDC and HOBt and subsequently heated in 10% aq. NaOH at 60 °C to generate the intermediate **65**, which was then subjected to alkylation with 4-chlorobenzyl bromide.

**Scheme 2.**
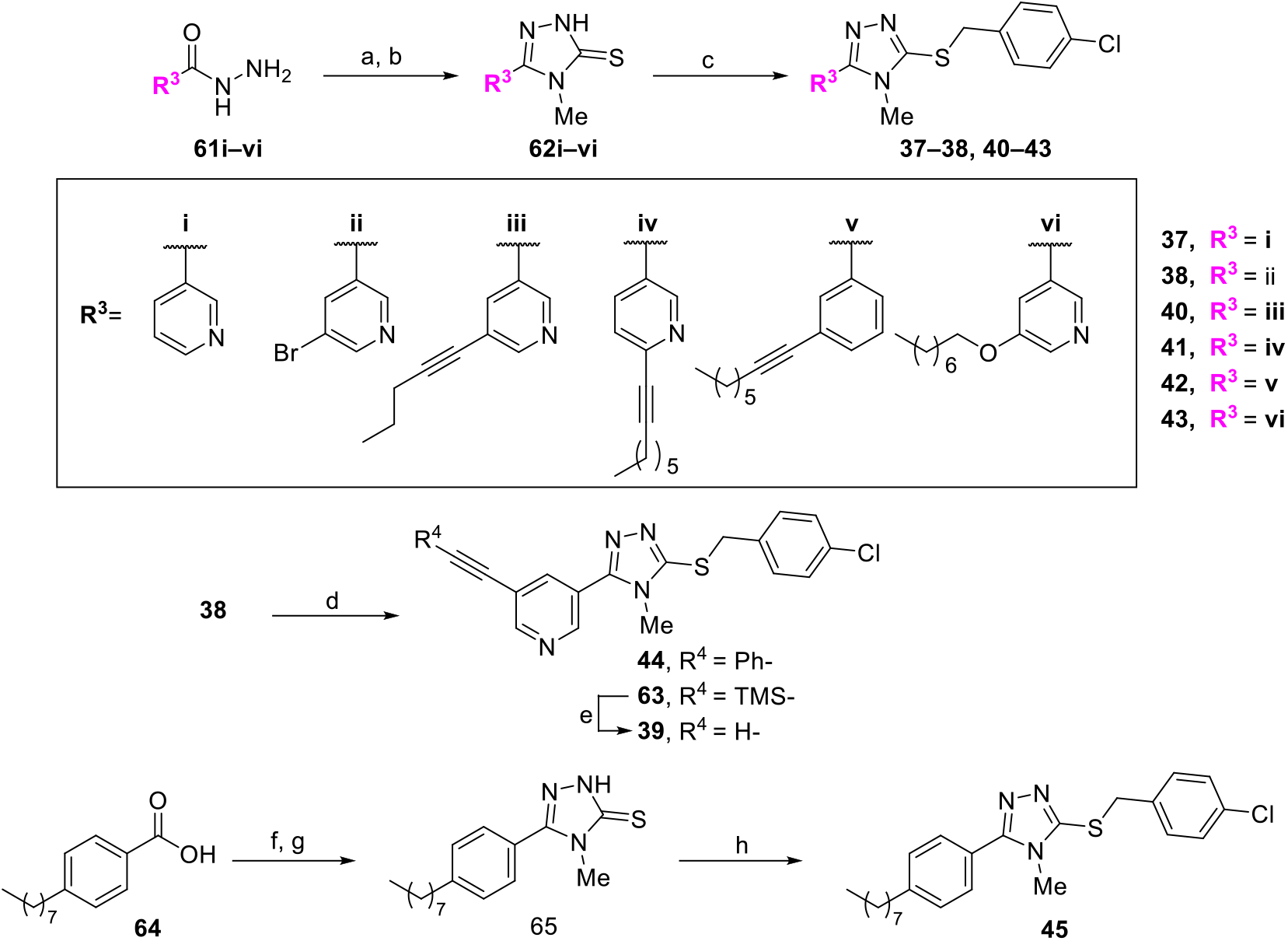
Synthesis of analogs with variations in the western region. ^[a]^ ^[a]^ Reagents and conditions: (a) methyl isothiocyanate, EtOH, reflux, 4 h; (b) 10% aq. NaOH, 60 °C, 3 h, 27–99% (over two steps); (c) 4-ClPhCH_2_Br, K_2_CO_3_, acetone/MeOH, rt, 4–14 h, 61–92%; for **43**, 4-ClPhCH_2_Br, Et_3_N, CH_3_CN, 4 h, 61%; (d) TMS-acetylene or phenylacetylene, Pd(PPh_3_)_2_Cl_2_, CuI, Et_3_N, rt, 18 h, 52–53%; (e) K_2_CO_3_, MeOH, rt, 4 h, 91%. (f) 4-methyl-3-thiosemicarbazide, HOBt, EDC, DIPEA, DMF, rt, 2 h; (g) 10% aq. NaOH 60 °C, 3 h, 89% (over two steps); (h) 4-ClPhCH_2_Br, K_2_CO_3_, acetone/MeOH, rt, 12 h, 92%.

#### Synthesis of analogs with replacement of the 1,2,4-triazole core (Scheme 3)

Core scaffold replacements were evaluated in the SAR studies by switching the 1,2,4-triazole with pyrrole (**46**), oxadiazole (**47**), thiadiazole (**48**), 1,2,3-triazole (**49**), pyridine (**50**), thiocarbamate (**51**), and thiourea (**52**), each of which followed distinct synthetic schemes. We also synthesized **53**, in which we flipped the western and eastern moieties of compound **1** around the 1,2,4-triazole-3-thiol core. The pyrrole core-containing analog **46** was synthesized by coupling the commercially available boronate **66** with the intermediate **67** and subsequent bromination with NBS, followed by coupling with 4-chlorobenzyl mercaptan. Both the oxadiazole (**48**) and thiadiazole (**49**) analogs were synthesized from hydrazide **54** (used in Scheme 1) following literature methods with slight modifications.^17^ The 1,2,3-triazole core (**49**) was obtained from the intermediate **67** via coupling with TMS acetylene, TMS deprotection, and an ensuing copper-catalyzed azide-alkyne cycloaddition (CuAAC) reaction with the azide (**72**). The pyridine-core analog **50** was synthesized by coupling the boronate ester (**74**) with the intermediate **67** via a Suzuki coupling reaction. The boronate **74** was synthesized via a substitution reaction of 3,5-dibromopyridine (**67**) with 4-chlorobenzyl mercaptan followed by a borylation reaction. The synthesis of thiocarbamate (**51**) and thiourea (**52**) analogs were assembled from the intermediate **76**. The intermediate **76** was subjected to a one-pot synthesis with triphosgene followed by concomitant addition of 4-chlorobenzyl mercaptan to construct the thiocarbamate core (**51**), whereas treatment of the intermediate **76** with thiophosgene to obtain **77** followed by the addition of 4-chlorobenylamine resulted in the thiourea core (**52**). Analog **53** was synthesized by alkylating the thione **78** (SI Scheme S9)^18^ with the alcohol **79** under Mitsunobu conditions.

**Scheme 3.**
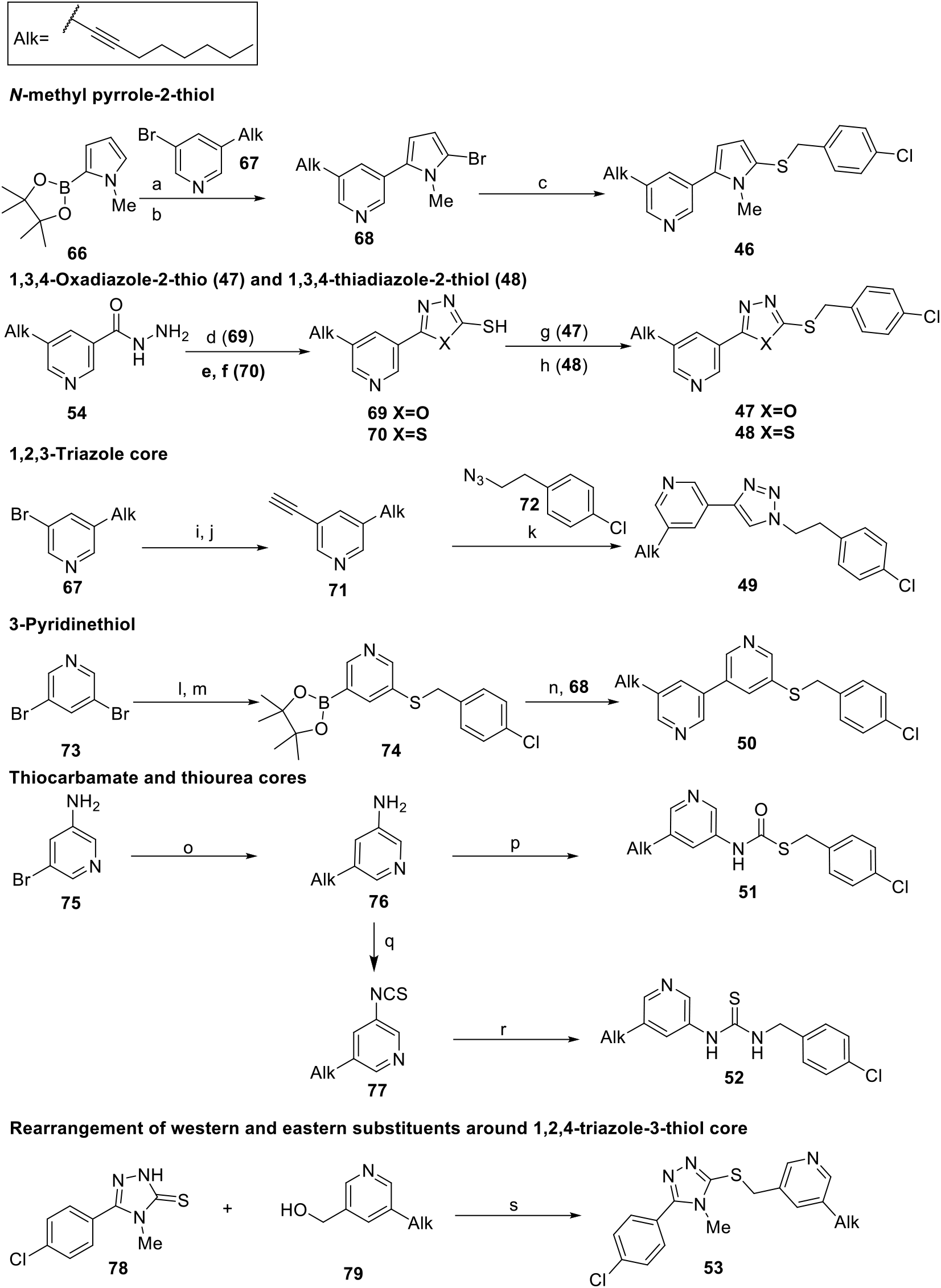
Synthesis of analogs with replacement of the 1,2,4-triazole core. ^[a]^ ^[a]^ Reagents and conditions: (a) Pd(PPh_3_)_4_, K_2_CO_3_, dioxane/H_2_O (4:1), 80 °C, 12 h, 33 %; (b) NBS, CH_2_Cl_2_, rt, 12 h, 85%; (c) 4-ClPhCH_2_SH, Xantphos, Pd_2_(dba)_3_, DIPEA, 1,4-dioxane, 15 h, 59%; (d) CS_2_, KOH, EtOH, reflux, 12 h, 80%; (e) CS_2_, KOH, EtOH, rt, 12 h; (f) H_2_SO_4,_ 0 °C, then rt, 4 h, 47%; (g) 4-ClPhCH_2_Br, K_2_CO_3_, acetone/MeOH, rt, 11 h, 76%; (h) 4-ClPhCH_2_Br, NaOH, tetrabutyl ammonium iodide, H_2_O/CH_2_Cl_2_, rt, overnight, 53%; (i) TMS acetylene, Pd(PPh_3_)_2_Cl_2_, CuI, Et_3_N, rt, 6 h; (j) K_2_CO_3_, MeOH, rt, 12 h, 47% over two steps; (k) CuSO_4_•5H_2_O, sodium ascorbate, CH_3_CN, 80 °C, 91%; (l) 4-ClPhCH_2_SH, NaH, DMF, 0 °C, 2 h then 80 °C, 12 h, 76%; (m) Pd(dbbf)Cl_2_, bis(pinacolato)diboron, KOAc,, toluene, 100 °C, 18 h; (n) Pd(PPh_3_)_4_, K_2_CO_3_, 1,4-dioxane/H_2_O (5:1), 85 °C, 63%; (o) oct-1-yne, Pd(PPh_3_)_2_Cl_2_, CuI, Et_3_N, rt, 12 h, 60 %; (p) triphosgene, 0 °C to rt, 4 h, then 4-ClPhCH_2_SH, CH_2_Cl_2_, overnight, 36%; (q) thiophosgene, NaOH, CH_3_CN, 0 °C to rt, CH_2_Cl_2_, 78%; (r) 4-ClPhCH_2_NH_2_, Et_3_N, CH_2_Cl_2_, 68%; (s) DIAD, PPh_3_, THF, 0 °C, 30 min, then rt, 3 h, 29%.

### 2.3. Structure-activity relationship (SAR) optimization

Our optimization campaign started with the modifications of the parent “hit” compound **1** in the eastern, southern, and western regions and modifications around the core, as highlighted in Figure 2. All synthesized compounds were tested for their *in vitro* anti-*Mtb* properties against the WT *Mtb* Erdman strain using the MABA.

#### SAR of the eastern and southern regions (Table 1)

We started by evaluating the role of substituents in the aromatic ring in the eastern region (Table 1). The 5-fold decrease in the activity of the methyl thioether analog lacking an aromatic ring **(2)** and a 35-fold decrease for the unsubstituted benzyl thioether (**3**) revealed that the substituted aromatic ring is important for maintaining anti-*Mtb* activity. Overall, we observed that *para*-substitution is essential for preserving activity and is tolerant to a wide range of groups, including halogens (**4** and **5**), electron-donating methoxy group (**7**), and electron-withdrawing groups (**8**,**9**). However, the introduction of trifluoromethyl (**10**) at this position resulted in an eight-fold decrease in activity, while a more hydrophobic *para*-phenyl (**11**) led to a complete loss in activity. Mono-substitution in the *meta*-position led to a three-fold reduction in activity for *meta*-chloro (**14)** and six-fold in the case of *meta*-nitrobenzyl analog (**15**).

**Table 1.**
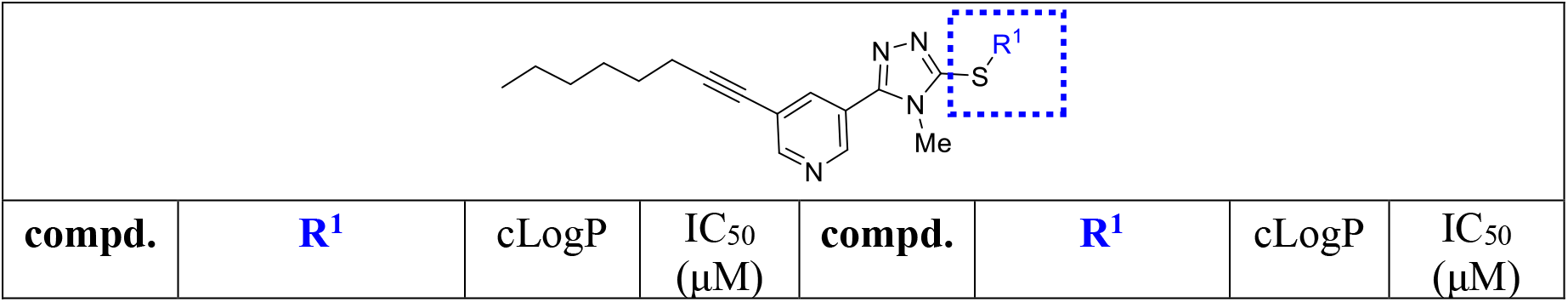
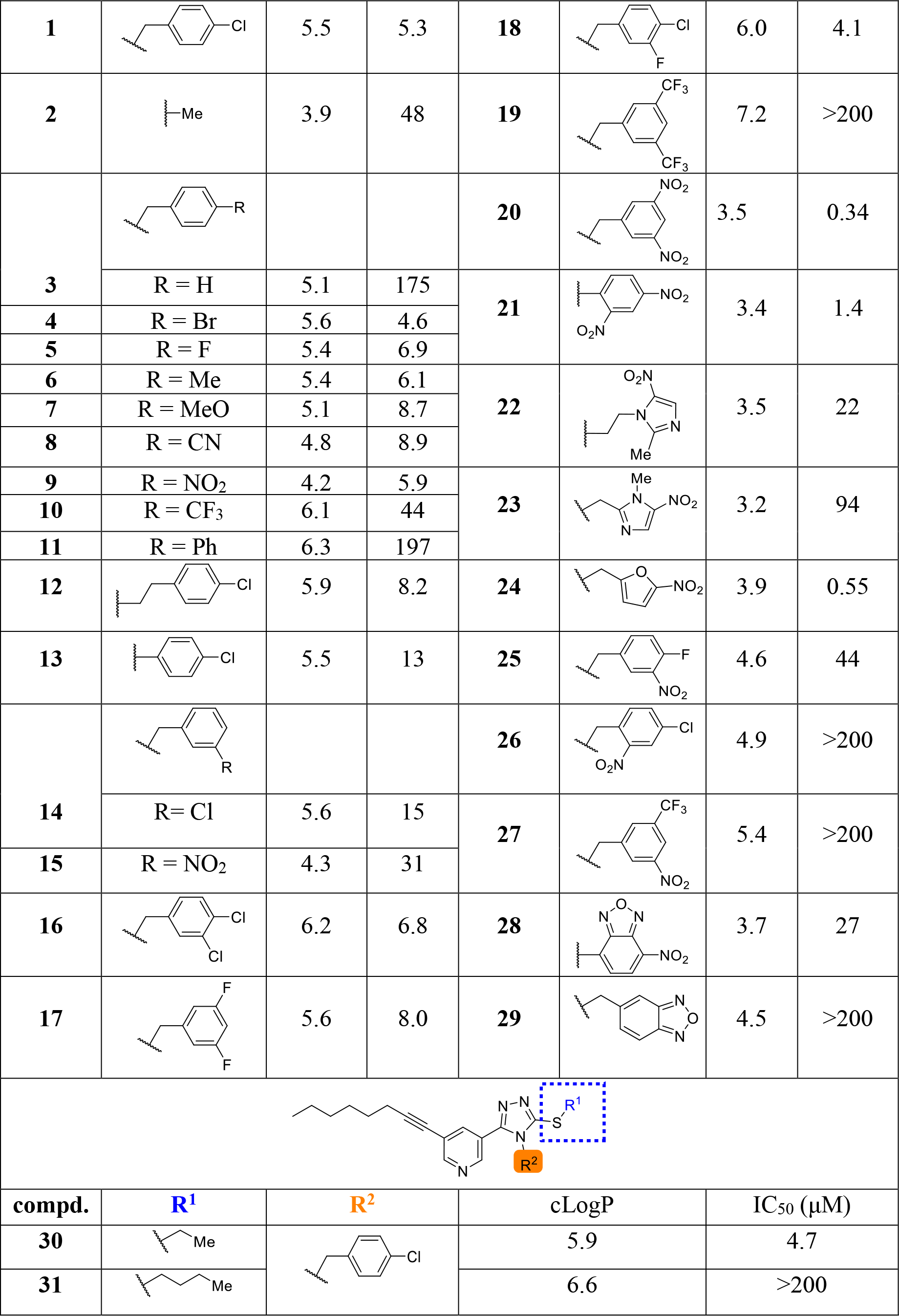
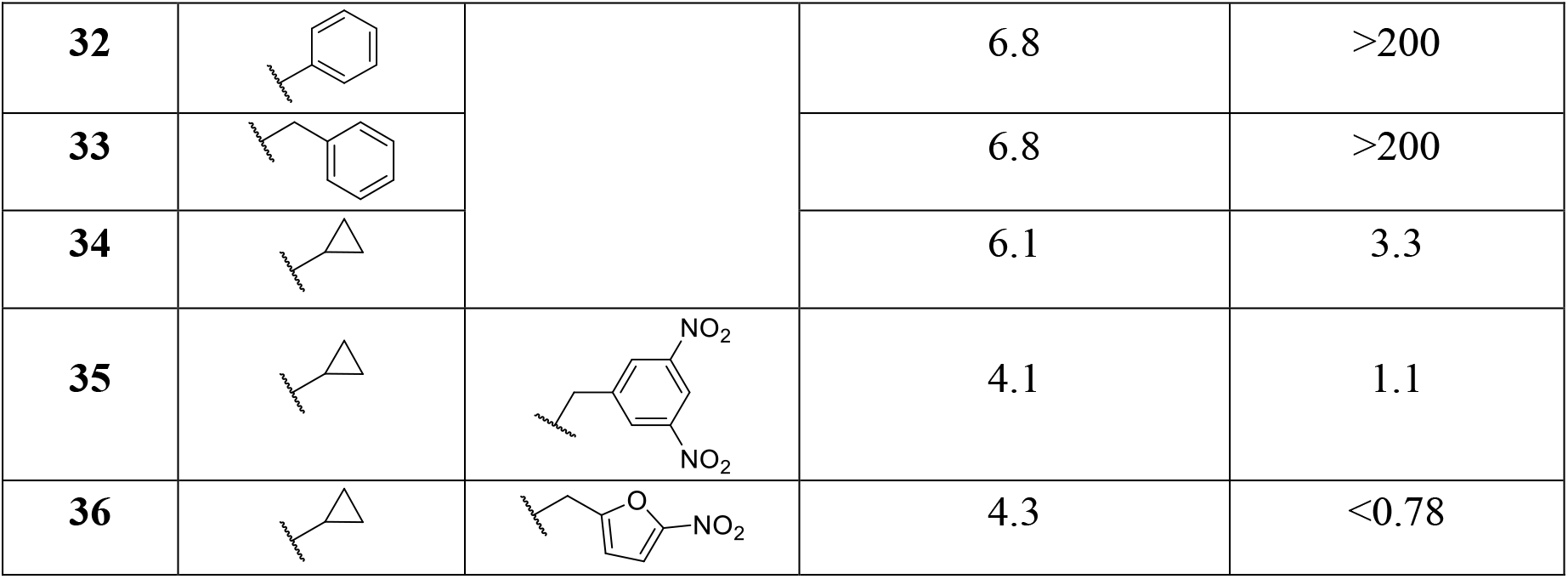
SAR for the eastern and southern regions of the 3-thio-1,2,4-triazole series. ^[a]^ ^[a]^ The calculated LogP values were determined using SwissADME web service. Anti-*Mtb* activity in wild-type (WT) *Mtb* strain determined from the MABA (n = 1).

Upon examining the disubstituted benzyl analogs (**16**–**20**), we observed that the di-halogenated analogs (**16**–**18**) maintained similar activity as the parent compound (**1**), while the bis-trifluoromethyl analog (**19**) lost its anti-*Mtb* activity. We observed that the 3,5-dinitrobenzyl analog (**20**) was highly potent with a low IC_50_ value of 0.34 μM. This observation led us to evaluate different nitro-substituted analogs in the eastern aromatic region (**21**–**25**). The SAR of the analogs where one of the nitro groups is in conjunction with halogen-substituents at the 3,4-(**25**) and 2,4-positions (**26)** suggests the *meta*-orientation of the di-nitro group is critical. Nitro-containing classes of antibiotics, such as nitroimidazoles and nitrofurans, have been in clinical use for several decades.^19, 20^ Nitro groups are also prevalent in TB drugs; examples include pretomanid and delamanide (Figure 1).^21^ We sought inspiration from these nitro-containing antibiotics and chose to evaluate analogs **22**–**24**, as they were amenable to our synthetic route and subsequent SAR evaluation. While the nitroimidazole analogs **22** and **23** possessed IC_50_ values of 22 μM and 94 μM, respectively, nitrofuran analog **24** exhibited improved potency with an IC_50_ value of 0.55 μM. A potential concern for nitro-containing drugs is their associated toxicity, which often prevents their further development in medicinal chemistry.^19^ Therefore, we evaluated a few bioisosteres of the nitro group, such as CF_3_ (**27**), nitrobenzoxadiazole (**28**), and benzoxadiazole (**29**).^22^ However, we observed either attenuated or complete loss in activity with bioisosteric replacements, indicating the importance of the 3,5-dinitrobenzyl motif.

We next inspected the role of *N*4-methyl substituent of the 1,2,4-triazole moiety by installing different lipophilic groups. The ethyl (**30**) and cyclopropyl (**34**) analogs were equipotent as the hit compound **1** with IC_50_ values of 4.7 μM and 3.3 μM, respectively. To determine if lipophilicity plays a role, we substituted with an *n*-butyl group on the *N*4-atom (**31)**, which turned out to furnish an inactive analog. Similarly, this position did not tolerate phenyl (**32**) or benzyl groups (**33**), perhaps due to steric hindrances or an unfavorable increase in lipophilicity. These observations reflected that the *N*4-methyl group could be modified with a cyclopropyl group (**34**–**36**), which in fact retained the activity.

#### SAR of the western region (Table 2)

The pyridyl ring and the position of its substituents (**37**–**45)** were evaluated in the western region, which revealed a clear correlation between the length of the lipophilic chain on C5 of the pyridyl ring and the resultant anti-*Mtb* activity. Compound **37** with no substitution at C5 was completely inactive, while compounds **39, 40**, and the parent compound **1** with two-, five-, and eight-carbon chain lengths, respectively, at the C5 position of the pyridyl ring exhibited an *Mtb* growth inhibition with IC_50_ values of 86 μM, 15 μM, and 5.3 μM, respectively (Table 2). This trend could be due to the level of lipophilicity imparted by the alkyne chain, as reflected by the higher cLogP values (compound **1** versus **37**, **39**, and **40**) (Table 2). The replacement of the linear eight-carbon oct-1-ynyl group in compound **1** with an eight-carbon phenylacetylene moiety in **44** resulted in similar activity (IC_50_ = 4 μM). Our results also suggest that the position of substituents on the pyridyl ring was crucial for the anti-*Mtb* activity. This was demonstrated by the loss of activity in **41**, in which the oct-1-ynyl chain was moved to the C6 position of the pyridyl ring. Similarly, analog **46** bearing 4-octylphenyl in the western region was inactive. The nitrogen atom of the pyridyl ring appears to play little role in the activity of this new subset of compounds, as its absence in **42** only led to a two-fold decrease in anti-*Mtb* activity. The replacement of the oct-1-ynyl group of **1** with the octyloxy- chain was tolerated resulting in the equipotent analog **43**.

**Table 2.**
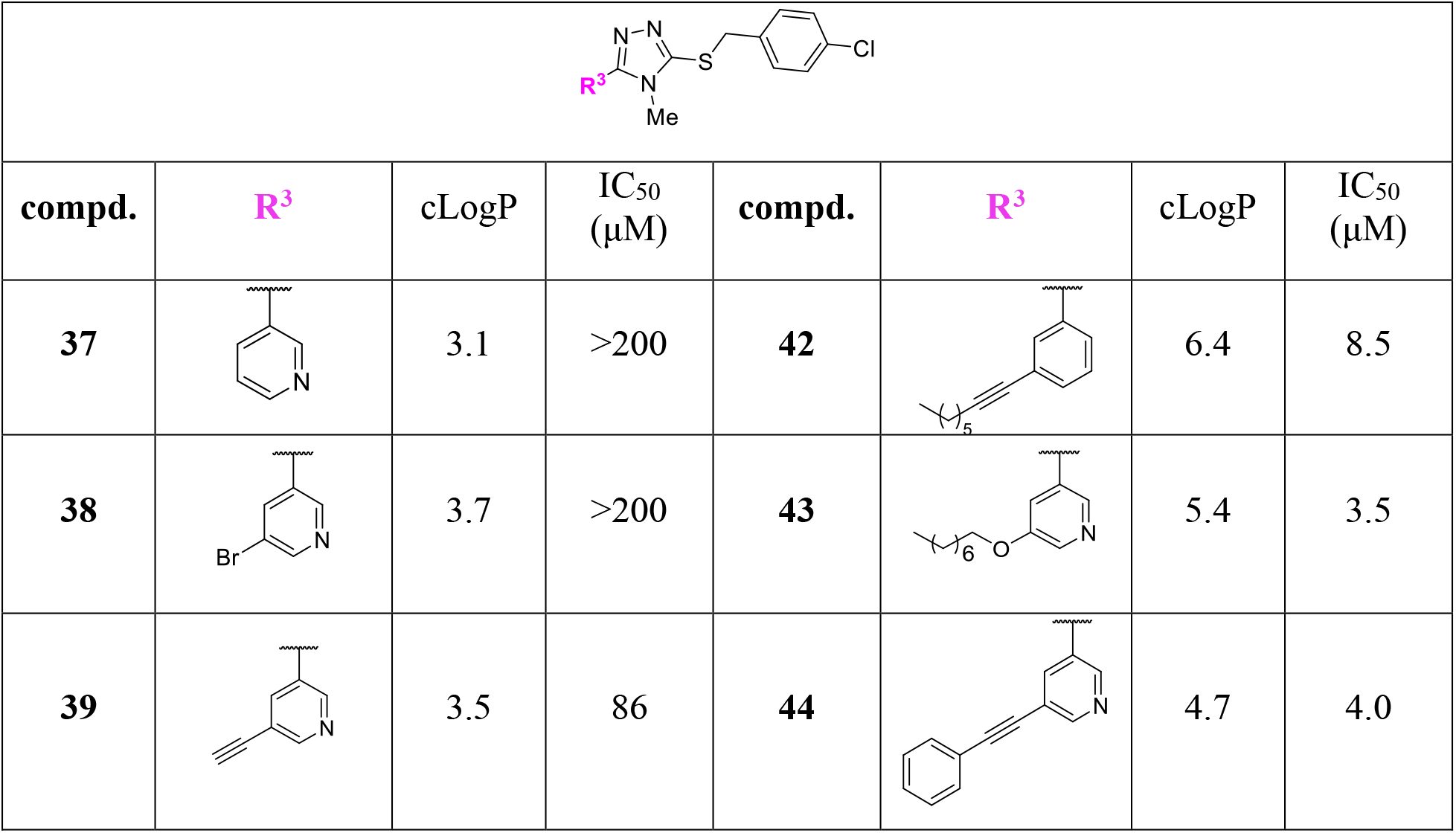
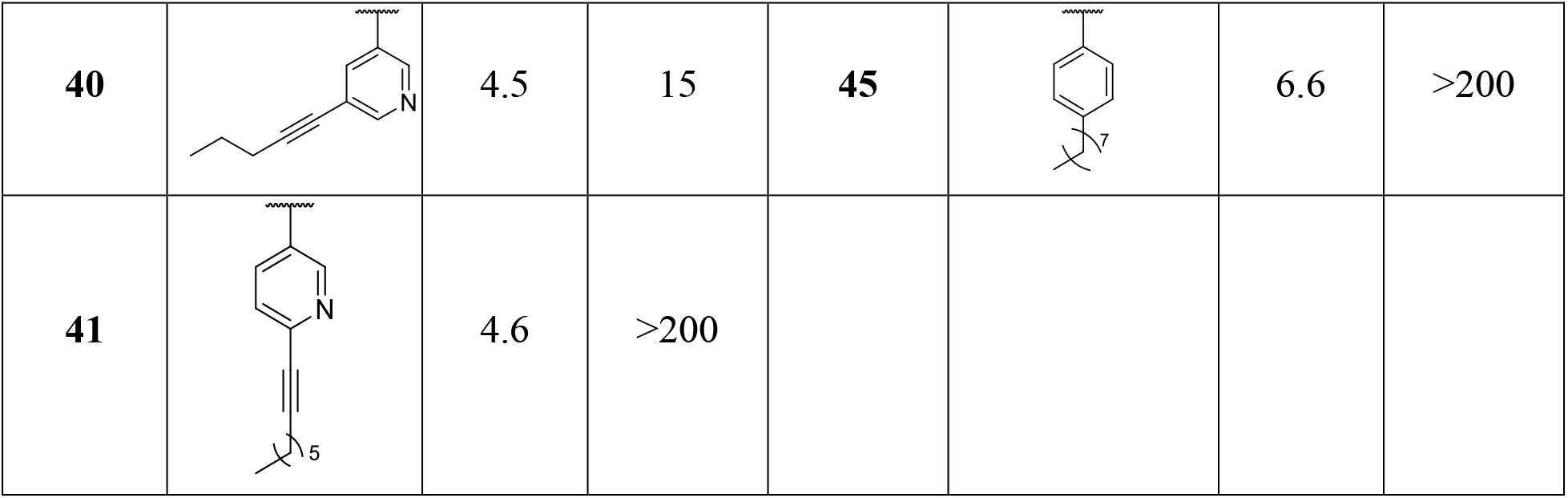
SAR for the western region of the 3-thio-1,2,4-triazole series. ^[a]^ ^[a]^ The calculated LogP values were determined using SwissADME web service. Anti-*Mtb* activity in wild-type (WT) *Mtb* strain determined from the MABA (n = 1).

|  |  |  |  |  |  |  |  |
| --- | --- | --- | --- | --- | --- | --- | --- |
| compd. | R<sup>3</sup> | cLogP | IC<sub>50</sub><br>($\mu M$) | compd. | R<sup>3</sup> | cLogP | IC<sub>50</sub><br>($\mu M$) |
| **37** |  | 3.1 | >200 | **42** |  | 6.4 | 8.5 |
| **38** |  | 3.7 | >200 | **43** |  | 5.4 | 3.5 |
| **39** |  | 3.5 | 86 | **44** |  | 4.7 | 4.0 |
| **40** |  | 4.5 | 15 | **45** |  | 6.6 | >200 |
| **41** |  | 4.6 | >200 |  |  |  |  |

#### SAR of the 1,2,4-triazole core (Table 3)

The replacement of the 1,2,4-triazole core with a pyrrole moiety (**46**) led to a three-fold decrease in activity, suggesting that the *N*1- and *N*2-of the triazole might have a role in the activity. Oxadiazole and thiadiazole are known bioisosteres of triazoles which led us to use these motifs as a core replacement to the design of analogs of 1,2,4-triazoles.^17, 23^ However, both oxadiazole (**47**) and thiadiazole (**48**) analogs showed a complete loss of anti-*Mtb* activity. We hypothesized this might be due to the absence of the *N*4-methyl group found in the hit compound (**1**). This assumption was further supported by the loss of activity in the 1,2,3-triazole (**49**) analog. To our delight, compound **50** bearing a 6-membered pyridine ring without a methyl substitution retained anti-*Mtb* activity (IC_50_ = 6.5 μM), which we hypothesize might be due to the similarity in the steric volumes of the pyridine and *N*4-methyl-1,2,4-triazole rings. We also tested **51** with thiocarbamate and **52** with thiourea core units, respectively. However, both the analogs were inactive. Swapping of the western region and eastern 4-chlorobenzene around the 1,2,4-triazole core (**53**) led to a loss of activity.

**Table 3.**
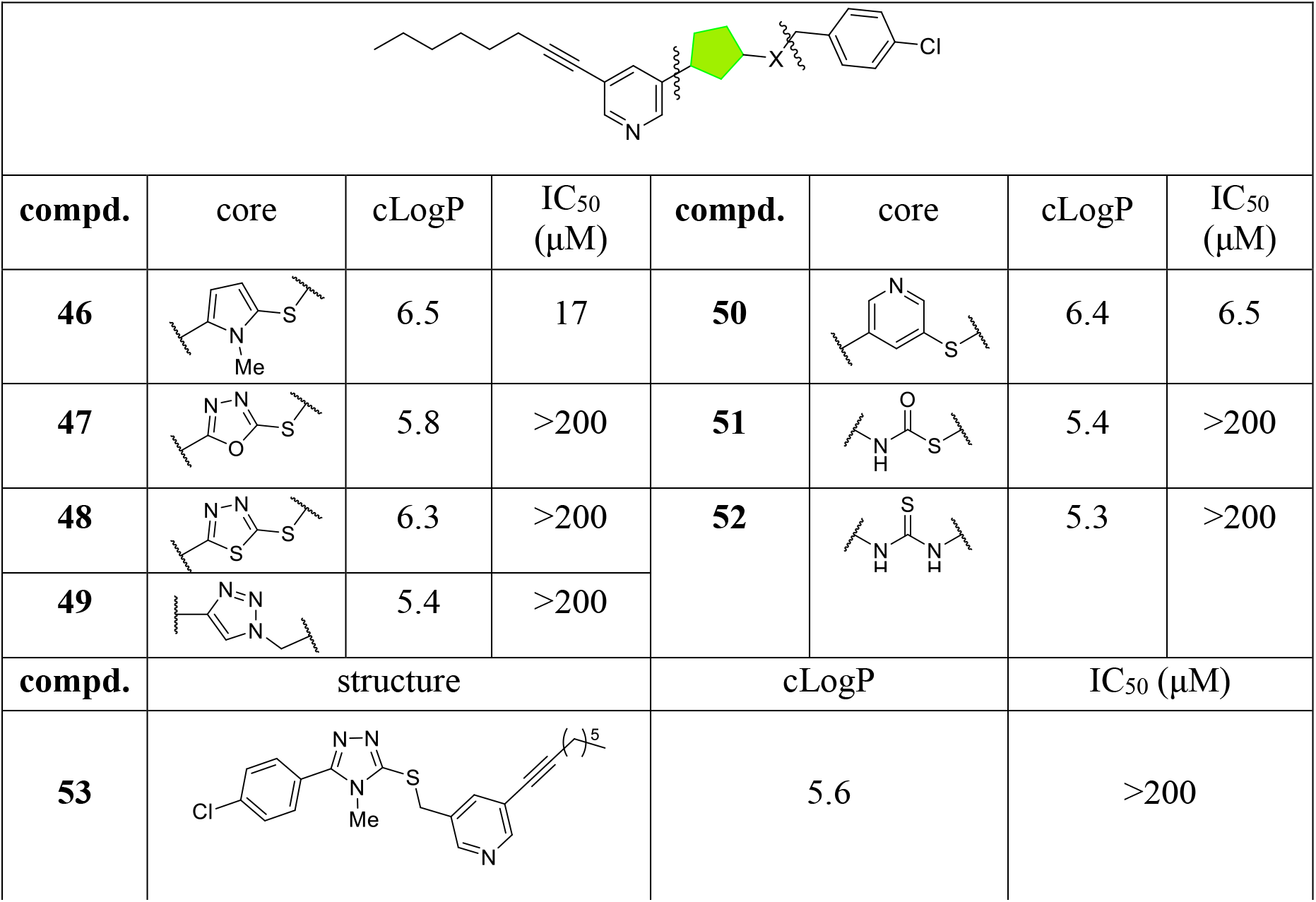
SAR for the 1,2,4-triazole core. ^[a]^ ^[a]^ The calculated LogP values were determined using SwissADME web service. Anti-*Mtb* activity in wild-type (WT) *Mtb* strain determined from the MABA (n = 1).

|  |  |  |  |  |  |  |  |
| --- | --- | --- | --- | --- | --- | --- | --- |
| compd. | core | cLogP | IC<sub>50</sub> (μM) | compd. | core | cLogP | IC<sub>50</sub> (μM) |
| 46 |  | 6.5 | 17 | 50 |  | 6.4 | 6.5 |
| 47 |  | 5.8 | >200 | 51 |  | 5.4 | >200 |
| 48 |  | 6.3 | >200 | 52 |  | 5.3 | >200 |
| 49 |  | 5.4 | >200 |  |  |  |  |
| compd. | structure |  |  | cLogP |  | IC<sub>50</sub> (μM) |  |
| 53 |  |  |  | 5.6 |  | >200 |  |

### 2.4 Lipophilicity trends within the 3-thio-1,2,4-triazole series

Antitubercular drugs tend to be more lipophilic compared to other antimicrobials;^24^ thus we were curious whether the trends in potency observed within the *3-thio-1,2,4-triazole* series correlated to lipophilicity. We calculated the LogP values of all the analogs using the SwissADME online data^25, 26, 27^ and plotted these values against IC_50_ in *Mtb*. We observed that all compounds that were active against *Mtb* have cLogP values clustered between 3.1 and 6.5 (Figure 3). The western region, particularly, had a stronger correlation between higher cLogP values of 4.3 to 6.3 being associated with more potent compounds. This trend indicates that the lipophilic tail is essential for activity. From the SAR studies of the analogs, we observed that overall lipophilicity is impacted by the lipophilicity of the western region.

**Figure 3.**
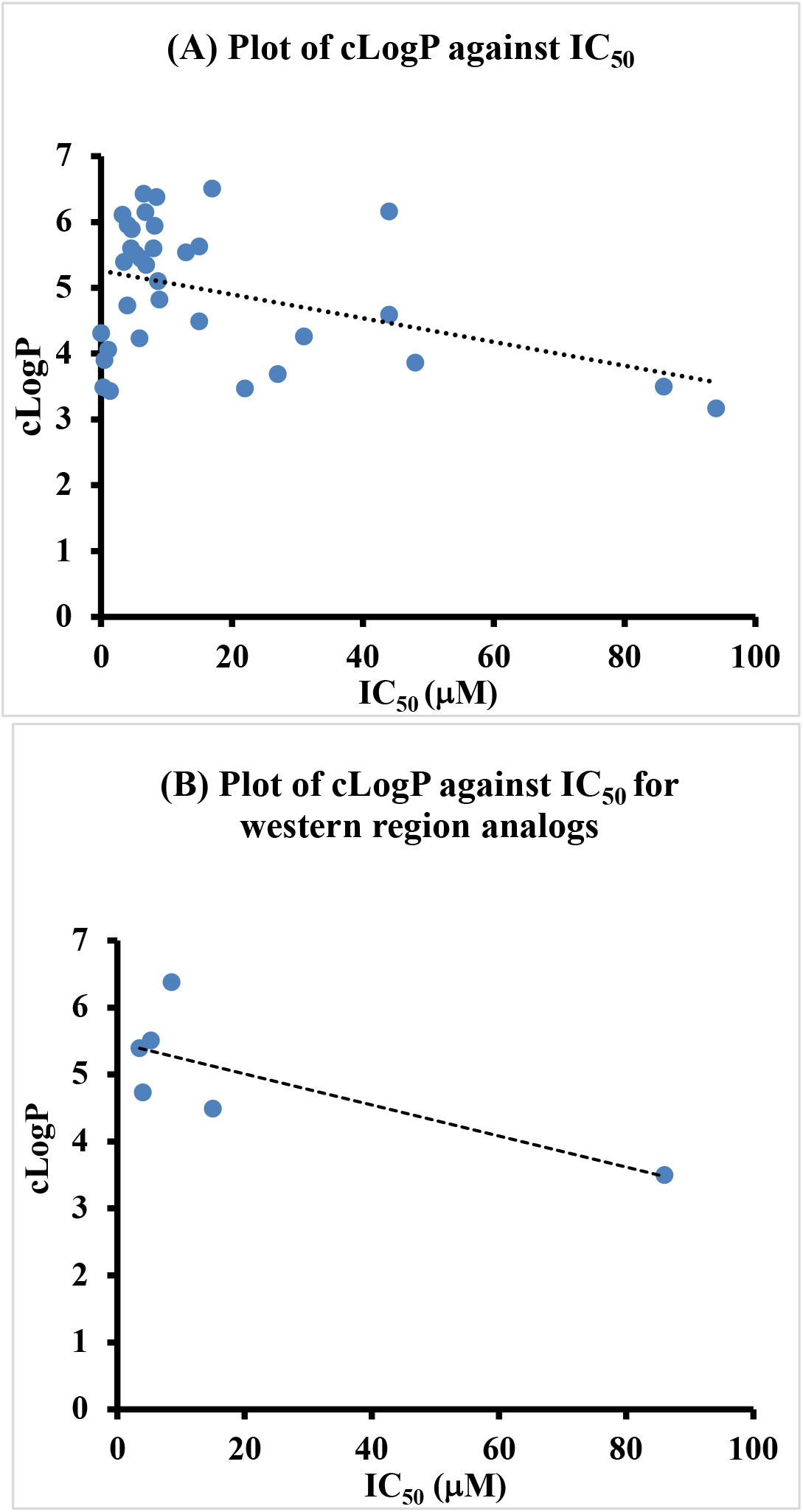
Plots of cLogP against IC_50_ (μM) of (A) all analogs with IC_50_ values <100 μM and (B) all active analogs in the western region (analogs **39**, **40**, **42**, **43**, **44**) and the hit compound **1**.

### 2.5. SAR summary and toxicity profiles of the most potent compounds

Following our initial screen to identify active compounds and based on our SAR evaluation, we selected compounds **1**, **8**, **9**, **18**, **20**, **21**, **25**, **34**, **36**, and **44** to be assayed in triplicate (Table 4) in the MABA for anti-*Mtb* activity to confirm their activity against WT *Mtb* Erdman (Figure 4, Table 4). Given the toxicity of our hit compound **1** in mammalian cells (Figure 2) and the potential concern for toxicity more generally with nitro-containing drugs^19^, we also examined the selected compounds for toxicity in a mammalian cell line using Promega’s Cell Titer-Glo assay to assess cellular ATP levels as a read-out of viability. We determined the concentration of the selected compound that resulted in a 50% decrease in cellular ATP (LD_50_) in Vero cells,^28^ an African green monkey kidney epithelial cell line, over a 72-hour incubation. We then calculated a selectivity index (SI) (Table 4) for each compound by determining the ratio of the LD_50_ in Vero cells to the MABA IC_50_ in *Mtb*, where the larger the SI, the safer the compound is in eukaryotic cells relative to its effective dose. From this analysis, we identified multiple highly potent anti-*Mtb* compounds with favorable selectivity indexes, some with SI > 100. Therefore, not only were our SAR efforts successful in identifying compounds with increased potency against *Mtb*, but also, we were able to overcome the toxicity concerns of the original hit compound.

**Figure 4.**
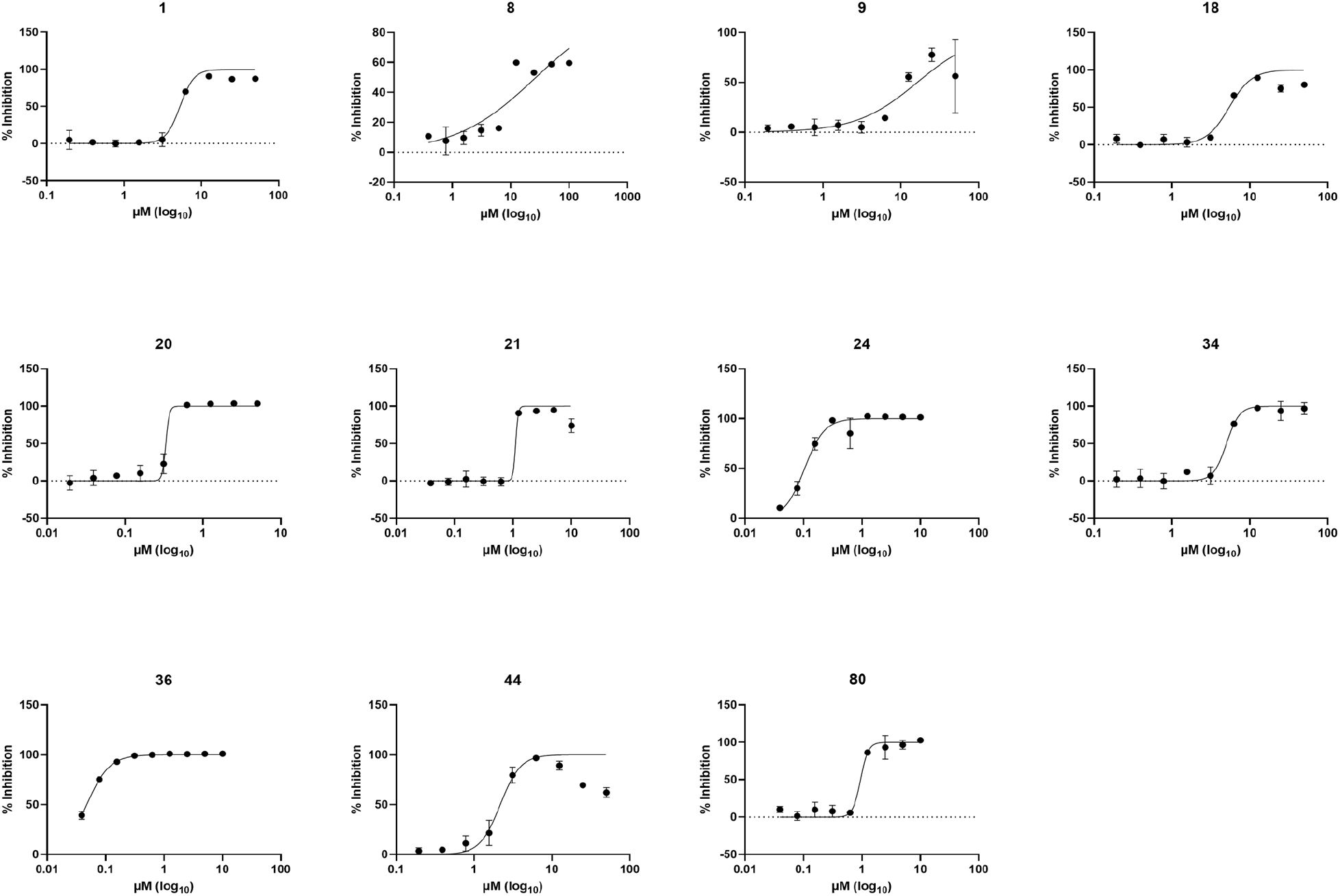
Concentration-response in the MABA for *in vitro* anti-*Mtb* activity of selected potent compounds listed in Table 4. WT *Mtb Erdman* strain was used in the MABA (n = 3).

**Table 4.**
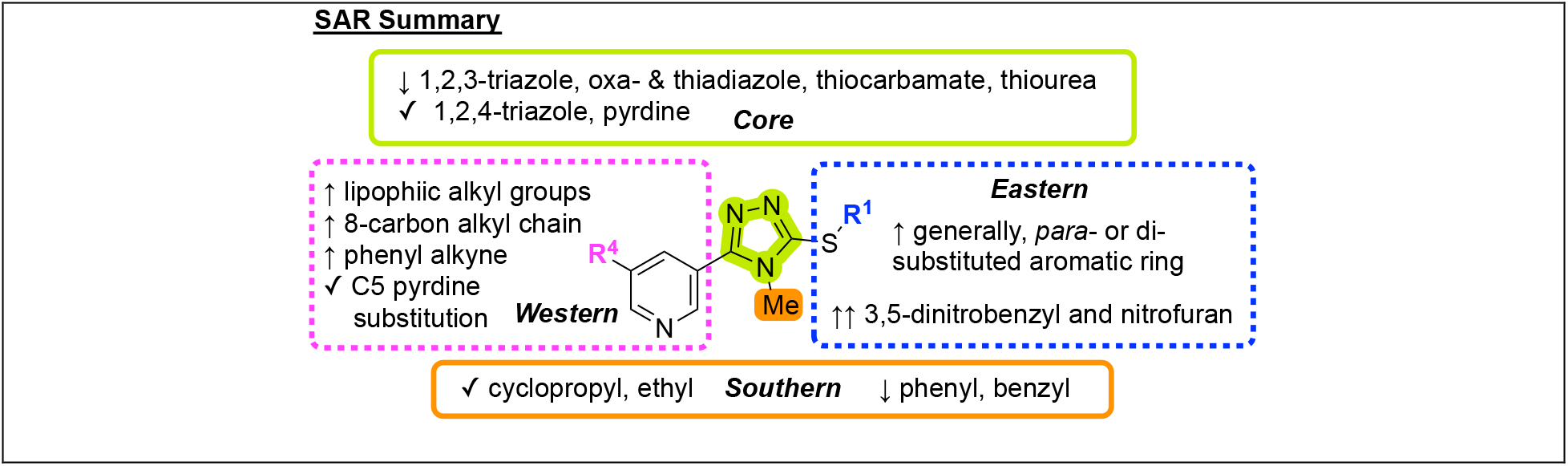
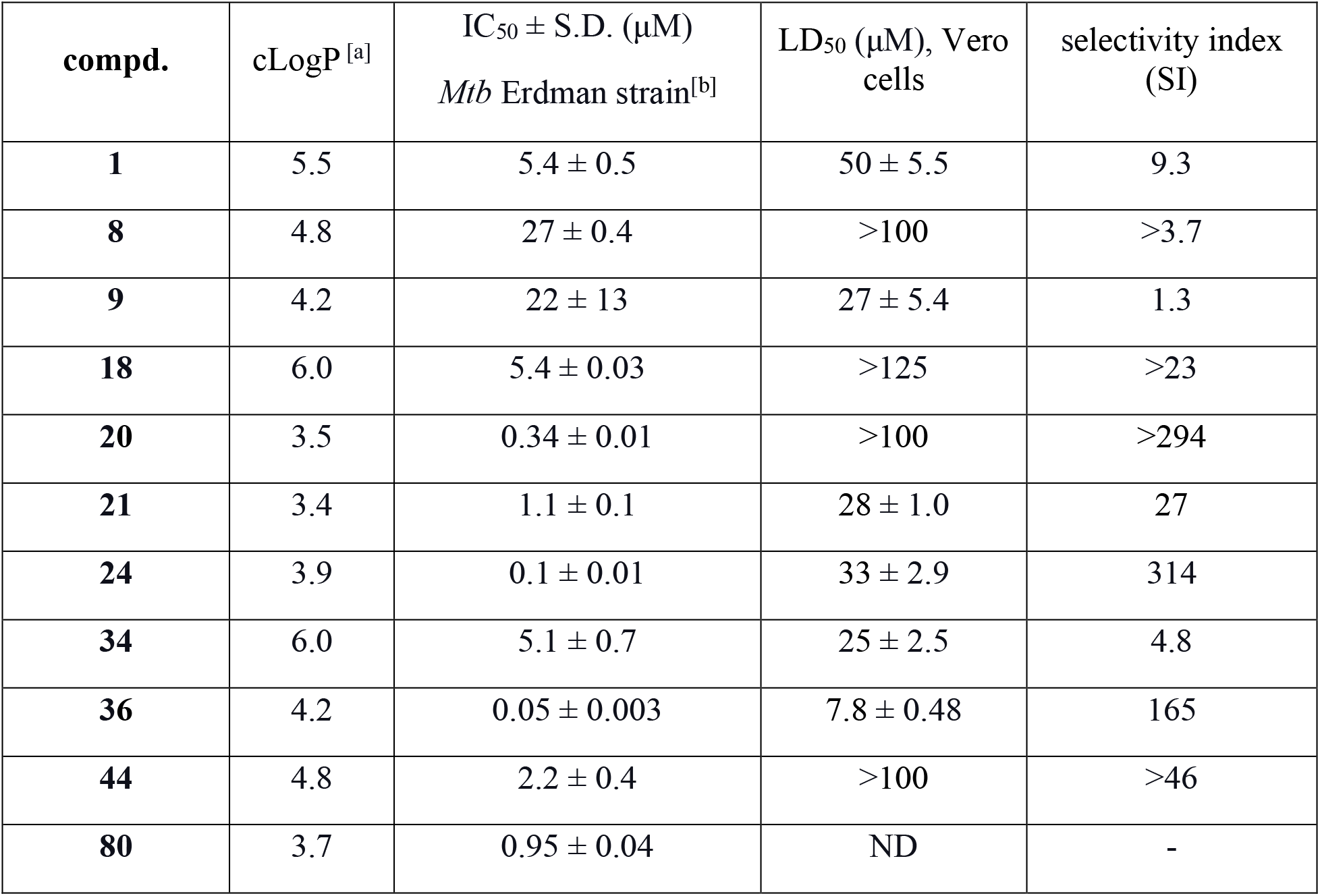
SAR trends and *in vitro* anti-*Mtb* activity and cytotoxicity of selected potent compounds with selectivity index (SI) values.^[a]^ ^[a]^ The calculated LogP values were determined using SwissADME web service. ^[b]^Anti-*Mtb* activity in wild-type (WT) *Mtb* Erdman strain was determined using the MABA (n = 3). LD_50_ in Vero cells is based on the Promega CellTiter-Glo Luminescent Cell Viability Assay (n = 3).

### 2.6. Activity against other mycobacteria and non-mycobacteria

The experiments described above were performed in a Lineage 4 strain of *Mtb* (Erdman). To examine if the compounds were active against another clinically relevant *Mtb* lineage, we determined the IC_50_ values of selected compounds **1**, **8**, **9**, and **20** in the hypervirulent HN878 *Mtb* strain of the W/Beijing Lineage (Lineage 2), which is increasing in incidence in active TB cases and is frequently associated with the occurrence of drug resistance.^29^ We observed that **1**, **8**, **9**, and **20** were twice as potent against HN878 *Mtb* compared to the Erdman strain (Table 5). We also investigated whether the 3-thio-1,2,4-triazole series was specific for activity against *Mtb* or if the compounds also exhibited activity against other mycobacteria and found that compounds **1**, **4**, **5**, **8**, and **9** did not show activity (MIC > 64 μg/mL or >120 μM) against *Mycobacterium smegmatis*, *Mycobacterium avium*, or *Mycobacterium abscessus* (SI Table S1–2). Furthermore, we screened **1**, **8**, **9**, **18**, **20**, **21**, **24**, **34**, **36**, **and 44** for activity against a panel of nonmycobacterial species, including Methicillin-resistant *Staphylococcus aureus* (MRSA), *Escherichia coli*, *Pseudomonas aeruginosa*, *Klebsiella pneumoniae*, and Vancomycin-resistant *Enterococcus faecium* (VRE). None of the tested compounds exhibited antimicrobial activity against the nonmycobacterial species at concentrations ≤16 μg/mL (< 38 μM) (SI Table S3). Therefore, the 3-thio-1,2,4-triazole series is specifically inhibitory for the growth of *Mtb*. This indicates that the 3-thio-1,2,4-triazole series may target a process that is particularly important in *Mtb*, that *Mtb* metabolism may more efficiently convert them to an active form, or that perhaps the *Mtb* cell envelope is more permeable to these compounds.

**Table 5.** Anti-*Mtb* activities (MIC values) of selected analogs against HN878 *Mtb* strains. ^[a]^Anti-*Mtb* activity in wild-type (WT) *Mtb* Erdman and HN878 strains were determined using the MABA (n = 3).

| entry | IC<sub>50</sub> ± S.D. (µM)<sup>[a]</sup> |  |
| --- | --- | --- |
|  | Erdman | HN878 |
| **1** | 1.5 ± 0.19 | 0.85 ± 0.043 |
| **8** | 5.3 ± 1.17 | 2.3 ± 0.2 |
| **9** | 2.9 ± 0.45 | 1.3 ± 0.15 |
| **20** | 0.17 ± 0.016 | 0.061 ± 0.012 |
| **INH** | 0.18 ± 0.014 | 0.12 ± 0.00051 |

### 2.7 Activity against mono-resistant strains of *Mtb* (Table 6)

An ideal new antibiotic for the treatment of *Mtb* will have activity in both drug-sensitive and drug-resistant *Mtb*. Therefore, we tested the activity of the 3-thio-1,2,4-triazoles **1**, **8**, and **9** against three mono-resistant *Mtb* strains: a rifampin-resistant (RIF: rpoBS450L) strain, an isoniazid-resistant (INH: katGdel) strain, and a moxifloxacin-resistant (MOX: gyrAD94K) strain. All three compounds maintained their activity in the INH-resistant, RIF-resistant, and MOX-resistant strains, demonstrating their utility against drug resistant *Mtb* isolates and suggesting that they act by a unique mode of action (Table 6).

**Table 6.** Anti-*Mtb* activities (MIC values) of selected analogs against clinically relevant mono-resistant *Mtb* strains. ^[a]^ ^[a]^ Activity was tested against rifampin-resistant (RIF: rpoBS450L) strain (ATCC #35838), isoniazid-resistant (INH: katGdel) strain (MS015), and a moxifloxacin-resistant (MOX: gyrAD94K) strains. Studies supported by NIAID Atomic. More information is provided in the SI.

| cpd. | MIC ( μg/mL) <sup>[a]</sup> | | | |
| --- | --- | --- | --- | --- |
|  | H37Rv | H37Rv<br>(rpoBS450L) | H37Rv<br>(katGdel) | H37Rv<br>(gyrAD94K) |
| **1** | 8 | 4 | 8 | 8 |
| **9** | 16 | 8 | 4 | 8 |
| **Isoniazid** | 0.06 | 0.06 | >2 | 0.06 |
| **Moxifloxacin** | 0.125 | 0.06 | 0.125 | >2 |

### 2.8. Preliminary target identification studies

Phenotypic screens for whole-cell anti-*Mtb* activity has been successful in identifying compounds that progress to the clinical stages of development. One example is bedaquiline which was approved in 2012 by the FDA for the treatment of MDR-TB.^30^ However, it has been noted that a majority of hits identified by phenotypic screens of diverse libraries act on a few well-known targets referred to as promiscuous targets. Examples of such *Mtb* targets include MmpL3, QcrB, and DprE1.^14^ In particular, inhibitors of MmpL3 and QcrB follow a similar profile to our compounds of having narrow spectrum activity against *Mtb* and but not against other mycobacterial and non-mycobacterial species.^31^ Since our hit compound was identified through a phenotypic screen, we investigated whether the 3-thio-1,2,4-triazole series of compounds inhibited the promiscuous targets MmpL3 and QcrB.

#### Activity against MmpL3 mutants (Table 7)

The mycobacterial membrane protein large (MmpL) protein family are members of a superfamily of transporters that utilize the proton motive force (PMF) to transport substrates across the mycobacterial cell membrane.^38^ Specifically, MmpL3 transports mycolic acid (MA) in the form of trehalose monomycolates (TMM) from the inner membrane to the outer membrane for incorporation into trehalose dimycolate (TDM) or transfer to arabinogalactan. Inhibition of MmpL3 results in the accumulation of TMM in the cytoplasm and the inhibition of TDM and mycolylated arabinogalactan formation, thereby leading to loss of outer membrane formation and eventual death of *Mtb*.^32^ MmpL3 has been identified as a target for multiple classes of compounds identified in phenotypic screens with *Mtb*.^14, 15, 33^ To determine whether the 3-thio-1,2,4-triazole compounds were targeting MmpL3, we compared the ability of compound **20** to inhibit *Mtb* H37Rv mc26206 expressing the WT MmpL3 protein from *Mtb* or mutated variants that have frequently been associated with resistance to MmpL3 inhibitors (MmpL3 G253E and MmpL3 L567P) (Table 7).^34^ The *Mtb* H37Rv mc26206 strains used in this analysis are auxotrophic for leucine and pantothenate, so they can be cultured under BSL2 conditions with amino acid supplements. Minimum inhibitory concentrations (MICs) that inhibit mycobacterial growth and metabolic activity by at least 80% (MIC^80^) were determined by performing the MABA with a range of concentrations for each compound and further monitoring bacterial growth spectrophotometrically at OD600. Similar to the controls RIF and amikacin, compound **20** exhibited comparable activity in all three strains expressing the MmpL3 variants, suggesting that it did not target this mycolic acid transporter.^38^ We observed that compound **20** exhibited decreased activity in the *Mtb* H37Rv mc26206 strain (MIC_80_ = 2 μM or 1 μg/mL, Table 7) relative to the IC_50_ value of 0.34 μM in *Mtb* Erdman, which could be due to the modifications of the *Mtb* H37Rv mc26206 strain that contribute to its auxotrophy and attenuation.

**Table 7.** Anti-*Mtb* activity against strains expressing variants of MmpL3. ^[a]^ ^[a]^ MICs are in μg/mL. MIC was read by resazurin color change.

| compd. | *Mtb* H37Rv mc2-6206 WT (MIC<sub>80</sub>) | *Mtb* H37Rv mc2-6206 [MmpL3 G253E] (MIC<sub>80</sub>) | *Mtb* H37Rv mc2-6206 [MmpL3 L567P] (MIC<sub>80</sub>) |
| --- | --- | --- | --- |
| **20** | 1 | 1 | 2 |
| **AMK** | 0.5 | 0.5 | 0.5 |
| **RIF** | 0.0625 | 0.0625 | 0.0625 |

<sup>[a]</sup> MICs are in µg/mL. MIC was read by resazurin color change.

#### Activity against ΔcydA mutants (Table 8)

Another promiscuous target from phenotypic screens in *Mtb* is QcrB, which is a subunit in the cytochrome bc_1_ complex.^15, 35^ This complex is localized at the plasma membrane and forms a super complex with the aa_3_-type cytochrome oxidase to catalyze the terminal electron transfer reaction in the mycobacterial electron transport chain. In addition to the cytochrome bc_1_:aa_3_ oxidase complex, *Mtb* encodes a second terminal oxidase, cytochrome bd. Cytochrome bc_1_:aa_3_ oxidase and cytochrome bd oxidase have somewhat overlapping roles, where the cytochrome bd oxidase can partially compensate for the loss of cytochrome bc1:aa3 oxidase activity. As a result, deletion of the genes encoding for cytochrome bd oxidase, *cydA* and *cydB*, results in hypersensitivity to QcrB inhibitors.^36, 37^ Therefore, we tested whether an *Mtb* Erdman *ΔcydA* mutant was more sensitive than WT *Mtb* Erdman to the original hit compound **1** and the more potent analog **20**. Unlike as observed with Q203, a QcrB inhibitor currently in clinical trial,^17^ the IC_50_ values for hit compound **1** and analog **20** were similar in WT and *ΔcydA Mtb* (Table 8), indicating that *the* 3-thio-1,2,4-triazole series of compounds does not target QcrB.

**Table 8.**
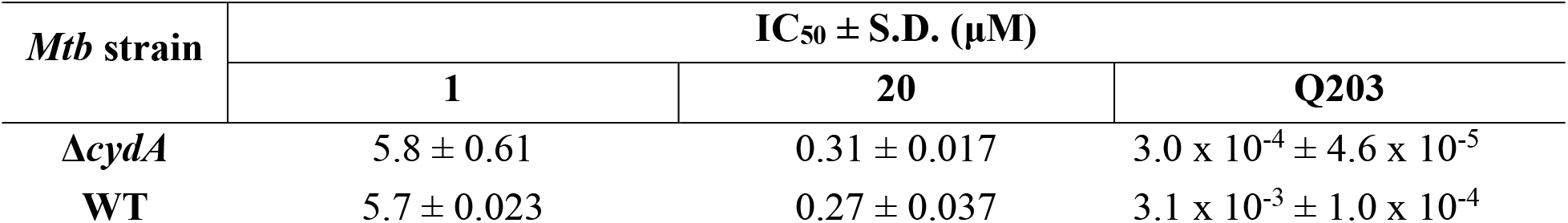
Anti-*Mtb* activity against Δ*cydA* mutants of *Mtb*^[a]^. ^[a]^ Anti-*Mtb* activity in WT *Mtb* Erdman and *ΔcydA Mtb* strain were determined using the MABA (n = 4).

| *Mtb* strain | IC<sub>50</sub> ± S.D. (μM) |  |  |
| --- | --- | --- | --- |
|  | **1** | **20** | **Q203** |
| $\Delta cydA$ | 5.8 ± 0.61 | 0.31 ± 0.017 | 3.0 x 10<sup>-4</sup> ± 4.6 x 10<sup>-5</sup> |
| WT | 5.7 ± 0.023 | 0.27 ± 0.037 | 3.1 x 10<sup>-3</sup> ± 1.0 x 10<sup>-4</sup> |

<sup>[a]</sup>Anti-*Mtb* activity in WT *Mtb* Erdman and $\Delta cydA$ *Mtb* strain were determined using the MABA (n = 4).

### 2.9 *In vitro* pharmacokinetic properties

Having identified nanomolar growth inhibitors of *Mtb* with good selectivity indices, we determined the aqueous solubility, human plasma protein binding, plasma stability, and liver microsomal stability of selected compounds (analogs **20**, **36**, and **80**) to establish a baseline for future analogs and provide insight into the potential for future *in vivo* studies. Compounds **20** and **36** showed a poor solubility profile comparable to that of reserpine, which is a reference for low solubility (Table 9). We next evaluated compound **80** to see if pyridinium salts improved solubility; however, the improvement was only marginal. The pyridinium salt **80** (synthesized from compound **35**, Scheme 4) retained its anti-*Mtb* potency with an IC_50_ of 0.95 μM compared to the free base **35 (**IC_50 =_ 1.2 μM) in MABA. We hypothesize the poor solubility might be due to the lipophilic side chain in the western region. Lipophilicity in TB drugs is often associated with low solubility that can be addressed by formulation or mode of administration, such as inhalation.^38^ We also observed poor Caco-2 cell permeability that might be due to the low aqueous solubility and high lipophilicity of these compounds (SI Table S4).^27^ Solubilizers, such as propylene glycol (PG), hydroxypropyl-beta-cyclodextrin (HP-beta-CD), and nanoparticles, are often used to enhance the bioavailability of drugs with poor aqueous solubility.^26^ In terms of metabolic stability, compound **20** exhibited an *in vitro* intrinsic clearance (CL_int_) of 287 and 357 μL/min/mg and degradation half-live (t_1⁄2_) of 4.8 and 3.9 min in mouse and human liver microsomes, respectively. We identified the presence of the thioether and *N*-demethylation at position *N*4-of the 1,2,4-triazole core as potential sources of metabolic instability. However, the introduction of the *N*4-cyclopropyl group (compounds **36** and **80**) did not circumvent the metabolic instability. Compounds **20** and **80** displayed excellent stability in both mouse and human plasma with t_1⁄2_ > 480 min (Table 9) when compared to propantheline (t_1⁄2 human_ = 21 min; t_1⁄2 mouse_ 24 min) and warfarin (t_1⁄2 human and mouse_ > 480 min). Compared to **20** and **80**, analog **36** was less stable in mouse and human plasma with a t_1⁄2_ of 238 min and 259 min, respectively. Compounds **20**, **36**, and **80** showed high levels human plasma protein binding (PPB >99%), similar to that of warfarin (PPB >99%). Although it was commonly thought that only the unbound drug is available to engage its targets, recent findings have suggested that compounds with >99% PPB remain valid drug candidates without requiring further property optimization.^39^

**Table 9.** Physicochemical parameters and *in vitro* ADME properties of compounds **20**, **36**, and **80**. ^[a]^ ^[a]^ Data collected at Cyprotex. ^[b]^Phosphate-buffered saline (PBS) solubility limit is highest concentration with no detectable precipitate. ^[c]^High solubility control and ^[d]^low solubility control for PBS solubility. Plasma stability data for compounds **20**, **36**, and **81**, in human and mouse plasma (n = 1). Liver microsomal stability data for **20**, **36**, and **80**, in human and mouse plasma (n = 1).

| cpd. | PBS solubility<sup>[b]</sup> (μM) | liver microsomal stability (human; mouse) |  | plasma stability (human; mouse) | human plasma protein binding |
| --- | --- | --- | --- | --- | --- |
| | | CL<sub>int</sub> (μL/min/mg) | $t_{1/2}$ (min) | $t_{1/2}$ (min) | |
| **20** | <1.6 | 357; 287 | 3.9; 4.8 | >480; >480 | >99.9% |
| **36** | 3.1 | 470; 546 | 1.9; 2.5 | 259; 238 | >99.9% |
| **80** | 3.1 | 725; 235 | 3.0; 5.9 | >480; >480 | >99.9% |
| **verapamil** | >500<sup>[c]</sup> | 62; 257 | 23; 5.4 | - | - |
| **reserpine** | 7.8<sup>[d]</sup> | - | - | - | - |
| **propantheline** | - | - | - | 21; 24 | - |
| **warfarin** | - | - | - | >480; >480 | >99.9% |

**Scheme 4.**
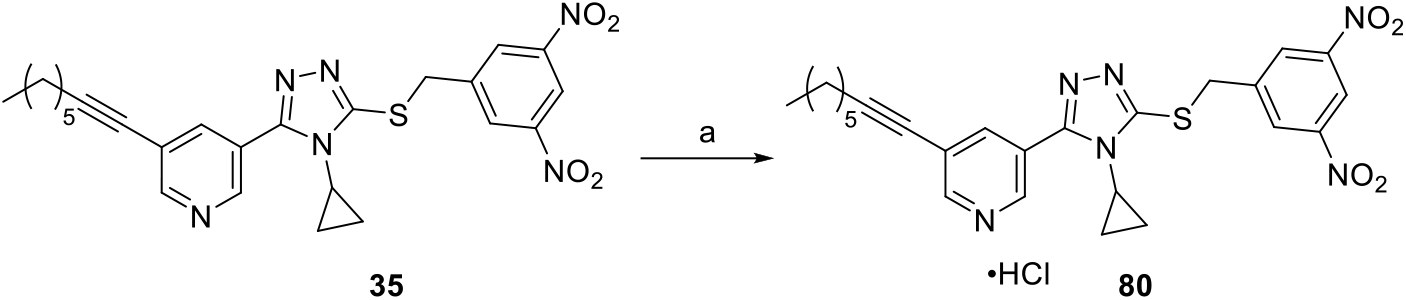
Synthesis of compound **80**. ^[a]^ ^[a]^ Reagents and conditions: (a) HCl, CH_2_Cl_2_, 1 h, rt, 84%.

## 3. CONCLUSION

In summary, a phenotypic screen and subsequent SAR optimization efforts to identify anti-*Mtb* agents have resulted in 3-thio-1,2,4-triazole compounds whose structure, activity, and properties were optimized to generate highly valuable candidates for the treatment of tuberculosis with nanomolar activity and potential for further development. The 3-thio-1,2,4-triazole series displays a selectivity towards *Mtb* and no activity against non-tuberculosis mycobacteria, MRSA, *E. coli, P. aeruginosa, K. pneumoniae*, and VRE. Moreover, the most potent analogs (**20** and **36**) exhibiting nanomolar activity, show high selectivity indices (>294 and 156, respectively), which alleviates the toxicity concern often associated with nitro-groups. Detailed studies are underway to identify the biological mechanism of action for these compounds. Our preliminary studies show that these compounds do not act on MmpL3 or QcrB, two promiscuous targets in *Mtb*, which provides evidence that they may display a unique mechanism of action. The SAR insight gained herein will be used for lead optimization studies focusing on improving metabolic stability and pharmacokinetic properties and for further development as potential drugs to combat MDR-TB.

## 4. METHODS

### General Methods and Instruments

All solvents and reagents were purchased from standard commercial vendors and used without further purification. Synthetic reactions were monitored using thin-layer chromatography (TLC) (Sorbtech silica XG TLC plates) and visualized under UV at 254 nm or with appropriate staining. Purification was performed using medium pressure liquid chromatography (MPLC) on a Biotage Isolera One using Biotage SNAP 10 g–50 g cartridges. NMR spectra were recorded on a Bruker Avance-500 or Bruker Avance-400 spectrometer at 298.15 K. Chemical shifts are reported in ppm using deuterated solvents (CDCl_3_, CD_3_OD, or DMSO-d_6_) for ^1^H and ^13^C NMR. CDCl_3_ (δ = 77.16 ppm), CD_3_OD (δ = 49.00 ppm) or DMSO-*d*_6_ (δ = 39.52 ppm) were used as internal standards for ^13^C NMR. For ^1^H NMR, CDCl_3_ (δ = 7.26 ppm), CD_3_OD (δ = 3.31 ppm) or DMSO-*d*_6_ (δ = 2.50 ppm) or TMS (δ = 0 ppm) were used as internal standards. Data were reported as: s = singlet, br = broad singlet, d = doublet, t = triplet, q = quartet, p = pentet, m = multiplet, b = broad, ap = apparent; coupling constants, *J*, in Hz. For high-resolution mass spectrometry (HRMS), a quadruple-TOF was used to obtain the data both in positive or negative modes. Attenuated total reflectance infrared (ATR-IR) spectroscopy was performed using an Agilent Technologies Cary 600 series FTIR Spectrometer. Purity (>95%) was determined using a Dionex Ultimate 3000 UPLC system (Thermo Fisher Scientific). Melting points were determined using an OptiMelt automated melting point system.

#### Chemistry

The general procedures of the synthesis of the final compounds listed in Table 4 are reported below. The synthesis and characterization of the remaining compounds and intermediates are found in the supporting information.

*General procedure A for alkylation of 4,5-disubstituted-1,2,4-triazole-2-thiones (compounds **1, 8**– **9**, **18**, **20**–**21**, **24, 34**, **36**, **44**)*

##### Method A1

The appropriate alkylating agent (bromide or chloride) (1.2 equiv) and the corresponding 1,2,4-triazole-3-thione (1.0 equiv) were dissolved in 1:2 parts of MeOH/acetone (0.1 M) at room temperature. Upon addition of K_2_CO_3_ (1.5 equiv), the mixture was stirred overnight at room temperature. The completion of the reaction was monitored by TLC, after which the solvent was evaporated under reduced pressure. The residue was dissolved with EtOAc, washed with water (x2), brine. The extracted organic layer was dried over anhydrous Na_2_SO_4_, and then concentrated under low pressure. The resultant residue was then purified by MPLC.

##### Method A2

^17^ The appropriate halide alkylating agent (1.2 equiv) was added to a solution of the corresponding 1,2,4-triazole-3-thiol (1.0 equiv) and Et_3_N (2.0 equiv) in CH_3_CN (0.1 M) and stirred for 4 h at room temperature. Upon completion of the reaction, the solvent was removed under reduced pressure. The residue was then dissolved in EtOAc and washed with water and brine. The extracted organic layer was dried over anhydrous Na_2_SO_4_, and concentrated under low pressure. The resultant residue was purified by MPLC.

#### 3-(5-((4-Chlorobenzyl)thio)-4-methyl-4H-1,2,4-triazol-3-yl)-5-(oct-1-yn-1-yl)pyridine (1)

1-(Bromomethyl)-4-chlorobenzene (150 mg, 1.2 equiv, 731 μmol) was reacted with 4-methyl-5-(5-(oct-1-yn-1-yl)pyridin-3-yl)-2,4-dihydro-3H-1,2,4-triazole-3-thione (**55**) (183 mg, 1.0 equiv, 609 μmol) according to general procedure A1. Final product was purified by MPLC (mobile phase: 40–80% EtOAc/Hex). Yield 79% (205 mg) as a white solid; mp 93–95 °C. ^1^H NMR (400 MHz, CDCl_3_) δ 8.71 (dd, *J* = 5.2, 2.1 Hz, 2H), 7.92 (t, *J* = 2.1 Hz, ^1^H), 7.34–7.23 (m, 4H), 4.42 (s, 2H), 3.43 (s, 3H), 2.44 (t, *J* = 7.1 Hz, 2H), 1.68–1.56 (m, 2H), 1.52–1.40 (m, 2H), 1.40–1.26 (m, 4H), 0.91 (t, *J* = 6.8 Hz, 3H). ^13^C NMR (101 MHz, CDCl_3_) δ 153.5, 153.0, 151.9, 146.8, 138.2, 135.4, 133.9, 130.6, 129.0, 123.0, 121.8, 96.1, 37.5, 31.8, 31.4, 28.7, 28.5, 22.6, 19.6, 14.2. HRMS (ESI): m/z [M+H]^+^ Calcd for [C_23_H_25_ClN_4_S + H]^+^ 425.1567, found 425.1565. HPLC Purity: 97.4%.

#### 4-(((4-Methyl-5-(5-(oct-1-yn-1-yl)pyridin-3-yl)-4H-1,2,4-triazol-3-yl)thio)methyl)benzonitrile (8)

4-(Chloromethyl)benzonitrile (24 mg, 1.2 equiv, 0.16 mmol) was reacted with 4-methyl-5-(5-(oct-1-yn-1-yl)pyridin-3-yl)-2,4-dihydro-3H-1,2,4-triazole-3-thione (**55**) (40 mg, 1.0 equiv, 0.13 mmol) according to general procedure A1. Final product was purified by MPLC (mobile phase: 40–80% EtOAc/Hex). Yield 87% (48 mg) as an off-white solid; mp 101– 103 °C ^1^H NMR (400 MHz, CDCl_3_) δ 8.73–8.67 (m, 2H), 7.92 (t, *J* = 2.1 Hz, 1H), 7.67–7.44 (m, 4H), 4.52 (s, 2H), 3.49 (s, 3H), 2.43 (t, *J* = 7.1 Hz, 2H), 1.61 (p, *J* = 7.1 Hz, 2H), 1.48–1.38 (m, 2H), 1.38–1.25 (m, 4H), 0.89 (t, *J* = 7.0 Hz, 3H) ^13^C NMR (101 MHz, CDCl_3_) δ 153.5, 153.1, 151.6, 146.7, 142.5, 138.2, 132.2, 130.0, 122.8, 121.8, 118.6, 111.9, 96.2, 76.5, 36.8, 31.8, 31.4, 28.7, 28.5, 22.6, 19.6, 14.2. (ESI): m/z [M+H] ^+^ Calcd for [C_24_H_25_N_5_S + H] ^+^ 416.1909, found 416.1920. HPLC Purity: 97.9%.

#### 3-(4-Methyl-5-((4-nitrobenzyl)thio)-4H-1,2,4-triazol-3-yl)-5-(oct-1-yn-1-yl)pyridine (9)

1-(Bromomethyl)-4-nitrobenzene (198 mg, 1.1 equiv, 915 μmol) was reacted with 4-methyl-5-(5-(oct-1-yn-1-yl)pyridin-3-yl)-2,4-dihydro-3H-1,2,4-triazole-3-thione (**55**) (250 mg, 1.0 equiv, 832 μmol) according to general procedure A1. Final product was purified by MPLC (mobile phase: 40–80% EtOAc/Hex). Yield 53% (192 mg) as a yellowish white solid; mp 96–98 °C. ^1^H NMR (500 MHz, CDCl_3_) δ 8.76–8.62 (m, 2H), 8.20–8.08 (m, 2H), 7.91 (t, *J* = 2.1 Hz, ^1^H), 7.66–7.53 (m, 2H), 4.57 (s, 2H), 3.51 (s, 3H), 2.42 (t, *J* = 7.1 Hz, 2H), 1.65–1.54 (m, 2H), 1.50–1.38 (m, 2H), 1.37–1.24 (m, 4H), 0.89 (t, *J* = 7.0 Hz, 3H). ^13^C NMR (126 MHz, CDCl_3_) δ 153.5, 153.2, 151.5, 147.5, 146.7, 144.5, 138.1, 130.2, 123.9, 123.8, 122.8, 121.77, 96.2, 76.5, 36.4, 31.8, 31.4, 28.7, 28.5, 22.5, 19.6, 14.2. HRMS (ESI): m/z [M+H] ^+^ Calcd for [C_23_H_25_N_5_O_2_S + H] ^+^ 416.1807, found 416.1831. HPLC Purity: 99.3%.

#### 3-(5-((4-Chloro-3-fluorobenzyl)thio)-4-methyl-4H-1,2,4-triazol-3-yl)-5-(oct-1-yn-1-yl)pyridine (18)

4-(Bromomethyl)-1-chloro-2-fluorobenzene (54 mg, 33 μL, 1.2 equiv, 0.24 mmol) was reacted with 4-methyl-5-(5-(oct-1-yn-1-yl)pyridin-3-yl)-2,4-dihydro-3H-1,2,4-triazole-3-thione (**55**) (60 mg, 1.0 equiv, 0.20 mmol) according to general procedure A1. Final product was purified by MPLC (mobile phase: 40–100% EtOAc/Hex). Yield 74% (65 mg) as light brown solid; mp 92–95 °C. ^1^H NMR (400 MHz, CDCl_3_) δ 8.72 (s, 2H), 7.99 (t, *J* = 2.0 Hz, 1H), 7.35–7.29 (m, 1H), 7.22 (dd, *J* = 9.6, 2.1 Hz, 1H), 7.15–7.11 (m, 1H), 4.46 (s, 2H), 3.51 (s, 3H), 2.44 (t, *J* = 7.1 Hz, 2H), 1.68–1.56 (m, 2H), 1.51–1.39 (m, 2H), 1.38–1.26 (m, 4H), 0.90 (t, *J* = 6.9 Hz, 3H). ^13^C NMR (101 MHz, CDCl_3_) δ 159.3, 156.9, 152.9, 152.0, 146.0, 138.8, 137.7 (d, *J* = 6.7 Hz), 130.9, 125.7 (d, *J* = 3.7 Hz), 123.2, 122.2, 120.7 (d, *J* = 17.6 Hz), 117.4 (d, *J* = 21.6 Hz), 96.7, 36.7, 31.9, 31.4, 28.7, 28.5, 22.7, 19.6, 14.2. HRMS (ESI): m/z [M+H] + Calcd for [C_23_H_24_ClFN_4_S + H] ^+^ 443.1472, found 443.1446. HPLC Purity: 99.0%.

#### 3-(5-((3,5-Dinitrobenzyl)thio)-4-methyl-4H-1,2,4-triazol-3-yl)-5-(oct-1-yn-1-yl)pyridine (20)

1-(Bromomethyl)-3-nitro-5-(trifluoromethyl)benzene (48 mg, 28 μL, 1.2 equiv, 0.17 mmol) was reacted with 4-methyl-5-(5-(oct-1-yn-1-yl)pyridin-3-yl)-2,4-dihydro-3H-1,2,4-triazole-3-thione (**55**) (42 mg, 1.0 equiv, 0.14 mmol) according to general procedure A1. Final product was purified by MPLC (mobile phase: 40–100% EtOAc/Hex). Yield 71% (57 mg) as of white solid; mp 144– 146 °C. ^1^H NMR (400 MHz, CDCl_3_) δ 8.94 (t, *J* = 2.1 Hz, 1H), 8.78–8.63 (m, 4H), 7.92 (t, *J* = 2.1 Hz, 1H), 4.72 (s, 2H), 3.60 (s, 3H), 2.43 (t, *J* = 7.1 Hz, 2H), 1.65 – 1.56 (m, 2H), 1.49–1.39 (m, 2H), 1.37–1.25 (m, 4H), 0.90 (t, *J* = 6.9 Hz, 3H). ^13^C NMR (126 MHz, CDCl_3_) δ 153.7, 153.5, 150.9, 148.7, 146.7, 141.9, 138.2, 129.5, 122.6, 121.9, 118.3, 96.3, 76.5, 35.1, 31.8, 31.4, 28.7, 28.5, 22.7, 19.6, 14.2. HRMS (ESI): m/z [M+H]^+^ Calcd for [C_23_H_24_N_6_O_4_S + H]^+^ 481.1658, found 481.1672. HPLC Purity: 99.4%.

#### 3-(5-((2,4-Dinitrophenyl)thio)-4-methyl-4H-1,2,4-triazol-3-yl)-5-(oct-1-yn-1-yl)pyridine (21)

1-Fluoro-2,4-dinitrobenzene (41 mg, 28 μL, 1.2 equiv, 0.22 mmol) was reacted with 4-
methyl-5-(5-(oct-1-yn-1-yl)pyridin-3-yl)-2,4-dihydro-3H-1,2,4-triazole-3-thione (**55**) (55 mg, 1.0 equiv, 0.18 mmol) according to general procedure A2. Final product was purified by MPLC (mobile phase:10–50% EtOAc/Hex). Yield 39% (33 mg) as a yellow solid; mp 156–157 °C. ^1^H NMR (400 MHz, CDCl_3_) δ 9.14 (d, *J* = 2.4 Hz, 1H), 8.87–8.72 (m, 2H), 8.30 (dd, *J* = 9.0, 2.5 Hz, 1H), 8.10 – 8.01 (m, 1H), 7.19 (d, *J* = 9.0 Hz, 1H), 3.77 (s, 3H), 2.45 (t, *J* = 7.1 Hz, 2H), 1.67– 1.55 (m, 2H), 1.50–1.39 (m, 2H), 1.38–1.27 (m, 4H), 0.90 (t, *J* = 7.0 Hz, 3H). ^13^C NMR (101 MHz, CDCl_3_) δ 155.0, 154.2, 146.8, 146.6, 146.0, 145.0, 141.5, 138.3, 129.2, 128.2, 122.3, 122.1, 121.9, 96.8, 76.3, 32.7, 31.4, 28.7, 28.5, 22.7, 19.6, 14.2. HRMS (ESI): m/z [M+H] ^+^ Calcd for [C_22_H_22_N_6_O_4_S + H] ^+^ 467.1501, found 467.1534. HPLC Purity: 100%.

#### 3-(4-Methyl-5-(((5-nitrofuran-2-yl)methyl)thio)-4H-1,2,4-triazol-3-yl)-5-(oct-1-yn-1-yl)pyridine (24)

2-(Bromomethyl)-5-nitrofuran (53 mg, 1.2 equiv, 0.26 mmol) was reacted with 4-methyl-5-(5-(oct-1-yn-1-yl)pyridin-3-yl)-2,4-dihydro-3H-1,2,4-triazole-3-thione (**55**) (64 mg, 1.0 equiv, 0.21 mmol) following general procedure A2. Final product was purified by MPLC (mobile phase: 5% MeOH/ CH_2_Cl_2_). Yield 25% (23 mg) yellow sticky solid. ^1^H NMR (400 MHz, CDCl_3_) δ 8.77–8.67 (m, 2H), 7.95 (t, *J* = 2.1 Hz, 1H), 7.22 (d, *J* = 3.7 Hz, 1H), 6.68 (d, *J* = 3.7 Hz, 1H), 4.58 (s, 2H), 3.60 (s, 3H), 2.44 (t, *J* = 7.1 Hz, 2H), 1.66–1.57 (m, 2H), 1.51–1.40 (m, 2H), 1.39–1.26 (m, 4H), 0.90 (t, *J* = 7.0 Hz, 3H). ^13^C NMR (101 MHz, CDCl_3_) δ 153.9, 153.7, 153.5, 151.0, 146.8, 138.2, 122.8, 121.9, 113.0, 112.8, 96.3, 76.5, 31.9, 31.4, 29.2, 28.7, 28.5, 22.7, 19.6, 14.2. HRMS (ESI): m/z [M+H] ^+^ Calcd for [C_21_H_23_N_5_O_3_S + H] ^+^ 426.1600, found 436.1601. HPLC Purity: 97.0%.

#### 3-(5-((4-Chlorobenzyl)thio)-4-cyclopropyl-4H-1,2,4-triazol-3-yl)-5-(oct-1-yn-1-yl)pyridine (34)

1-Chloro-4-(chloromethyl)benzene (30 mg, 1.2 equiv, 0.19 mmol) was reacted with 4-cyclopropyl-5-(5-(oct-1-yn-1-yl)pyridin-3-yl)-2,4-dihydro-3H-1,2,4-triazole-3-thione (**60**) (51 mg, 1.0 equiv, 0.16 mmol) according to general procedure A1. Final product was purified by MPLC (mobile phase: 20-60 EtOAc/Hex). Yield 71% (50 mg) yellowish sticky solid. ^1^H NMR (400 MHz, CDCl_3_) δ 8.88 (s, 1H), 8.69 (s, 1H), 8.10 (t, *J* = 1.8 Hz, 1H), 7.41 (d, *J* = 8.4 Hz, 2H), 7.29 (d, *J* = 8.4 Hz, 2H), 4.53 (s, 2H), 3.17–3.07 (m, 1H), 2.44 (t, *J* = 7.1 Hz, 2H), 1.67–1.56 (m, 2H), 1.50–1.39 (m, 2H), 1.39–1.23 (m, 4H), 1.15–1.06 (m, 2H), 0.90 (t, *J* = 7.0 Hz, 3H), 0.74– 0.66 (m, 2H). ^13^C NMR (101 MHz, CDCl_3_) δ 154.7, 153.2, 152.6, 146.5, 146.4, 138.3, 135.4, 133.8, 130.8, 128.9, 123.3, 121.6, 96.0, 76.6, 35.8, 31.4, 28.7, 28.5, 25.8, 22.6, 19.6, 14.2, 9.4. HRMS (ESI): m/z [M+H] ^+^ Calcd for [C_25_H_27_ClN_4_S + Cs] ^+^ 583.0699, found 583.0692. HPLC Purity: 97.0%.

#### 3-(4-Cyclopropyl-5-(((5-nitrofuran-2-yl)methyl)thio)-4H-1,2,4-triazol-3-yl)-5-(oct-1-yn-1-yl)pyridine (36)

2-(Bromomethyl)-5-nitrofuran (39 mg, 1.2 equiv, 0.19 mmol) was reacted with 4-cyclopropyl-5-(5-(oct-1-yn-1-yl)pyridin-3-yl)-2,4-dihydro-3H-1,2,4-triazole-3-thione (**60**) (51 mg, 1.0 equiv, 0.16 mmol) according to general procedure A2. Final product was purified by MPLC (mobile phase: 20–50% EtOAc/Hex). Yield 43% (30 mg) as an off-white solid; mp 111– 113 °C. ^1^H NMR (400 MHz, CDCl_3_) δ 8.87 (d, *J* = 2.1 Hz, 1H), 8.68 (d, *J* = 2.0 Hz, 1H), 8.07 (t, *J* = 2.1 Hz, 1H), 7.22 (d, *J* = 3.7 Hz, 1H), 6.74 (d, *J* = 3.7 Hz, 1H), 3.25–3.15 (m, 1H), 2.44 (t, *J* = 7.1 Hz, 2H), 1.66–1.53 (m, 2H), 1.50–1.38 (m, 2H), 1.38–1.25 (m, 4H), 1.18–1.09 (m, 2H), 0.89 (t, *J* = 7.1 Hz, 3H), 0.77–0.67 (m, 2H). ^13^C NMR (101 MHz, CDCl_3_) δ 154.2, 153.9, 153.3, 153.3, 146.9, 138.0, 122.9, 121.5, 113.2, 112.9, 95.8, 76.7, 31.4, 28.7, 28.5, 27.9, 25.8, 22.7, 19.6, 14.2, 9.3. HRMS (ESI): m/z [M+H] ^+^ Calcd for [C_23_H_25_N_5_O_3_S + H] ^+^ 452.1756, found 452.1752. HPLC Purity: 97.8%.

#### 3-(5-((4-Chlorobenzyl)thio)-4-methyl-4H-1,2,4-triazol-3-yl)-5-(phenylethynyl)pyridine (44)

A mixture of the 3-bromo-5-(5-((4-chlorobenzyl)thio)-4-methyl-4H-1,2,4-triazol-3-yl)pyridine (**38**) (38 mg, 1.0 equiv, 96 μmol), bis(triphenylphosphine)palladium(II) chloride (3.4 mg, 0.05 equiv, 4.8 μmol), and copper(I) iodide (1.8 mg, 0.1 equiv, 9.6 μmol) in DMF (6 mL) was purged thoroughly with argon and treated with ethynylbenzene (12 mg, 0.01 mL, 1.25 equiv, 0.12 mmol) previously stirred with Et_3_N (19 mg, 27 μL, 2.0 equiv, 0.19 mmol) under argon in a separate vial. The reaction mixture was stirred at room temperature for 18 h. The reaction mixture was diluted with NaHCO_3_, filtered with celite pad. The filtrate was extracted with EtOAC, and the organic phase was washed with water and brine, dried over Na_2_SO_4_. and concentrated under reduced pressure. The residue was purified by MPLC (mobile phase: 100% EtOAc). Yield 52% (21 mg) as a white solid; mp 171–173 °C. ^1^H NMR (500 MHz, CDCl_3_) δ 8.89–8.83 (m, 1H), 8.80– 8.74 (m, 1H), 8.07 (t, *J* = 2.1 Hz, 1H), 7.59–7.52 (m, 2H), 7.42 – 7.36 (m, 3H), 7.33–7.25 (m, 4H), 4.44 (s, 2H), 3.47 (s, 3H). ^13^C NMR (126 MHz, CDCl_3_) δ 153.3, 152.9, 152.1, 147.4, 138.1, 135.4, 134.0, 132.3, 132.2, 131.9, 130.6, 129.3, 129.0, 128.7, 123.2, 122.2, 121.1, 94.3, 85.0, 37.5, 31.8. HRMS (ESI): m/z [M+H] ^+^ Calcd for [C_23_H_17_ClN_4_S + H] ^+^ 417.0941, found 417.0934. HPLC Purity: 97.3%.

#### 3-(4-Cyclopropyl-5-((3,5-dinitrobenzyl)thio)-4H-1,2,4-triazol-3-yl)-5-(oct-1-yn-1-yl)pyridine hydrochloride (80)

To 3-(4-cyclopropyl-5-((3,5-dinitrobenzyl)thio)-4H-1,2,4-triazol-3-yl)-5-(oct-1-yn-1-yl)pyridine (30 mg, 1.0 equiv, 59 μmol) dissolved in CH_2_Cl_2_ (0.3 mL) was added HCl in diethyl ether HCl (2.4 mg, 33 μL, 2 molar, 1.1 equiv, 65 μmol) and the mixture was stirred for 1 h. The solvent was evaporated under reduced pressure and the residue triturated with diethyl ether to obtain product as an off-white solid; mp 102–103 °C. Yield was 84% (27 mg). ^1^H NMR (400 MHz, MeOD) δ 8.98–8.93 (m, 1H), 8.92–8.88 (m, 1H), 8.87–8.80 (m, 3H), 8.46 (t, *J* = 2.0 Hz, 1H), 4.81 (s, 2H), 3.57–3.50 (m, 1H), 2.51 (t, *J* = 7.1 Hz, 2H), 1.70–1.59 (m, 2H), 1.53–1.44 (m, 2H), 1.41–1.30 (m, 4H), 1.21–1.14 (m, 2H), 0.95–0.88 (m, 3H), 0.84–0.78 (m, 2H). ^13^C NMR (126 MHz, MeOD) δ 157.6, 152.9, 150.6, 149.9, 144.8, 144.1, 143.2, 130.8, 125.1, 124.3, 118.9, 100.6, 75.6, 35.5, 32.5, 29.7, 29.3, 27.8, 23.6, 20.2, 14.4, 9.7. HRMS (Free base) (ESI): m/z [M+H]^+^ Calcd for [C_25_H_26_N_6_O_4_S + H] ^+^ 507.1815, found 507.1812. HPLC Purity: 96.5 %.

#### Microplate Alamar Blue Assay (MABA)

WT *Mtb* Erdman or Δ*cydA*^36^ *Mtb* was cultured in Middlebrook 7H9 liquid media supplemented with 60 μL/L oleic acid, 5 g/L bovine serum albumin, 2 g/L dextrose, 0.003 g/L catalase (OADC), 0.5% glycerol, and 0.05% Tween 80 at 37°C The bacteria were inoculated at a final OD_600_ of 1.6 x 10^−3^ in 200μL per well in 96-well plates with two-fold titrations of compounds. The concentration of DMSO was maintained at 1% for all wells to avoid toxicity. The 96-well plates were incubated in a humidified incubator at 37°C and 5% CO_2_ for one week. After the week incubation, 32.5 μL of a mixture containing an 8:5 ratio of 0.6 mM resazurin (Sigma) dissolved in 1X PBS to 20% Tween 80 was added and the production of fluorescent resorufin was measured after incubation at 37°C in 5% CO_2_ for 24 hours. Relative fluorescence units (RFU) were measured using a BioTek Synergy H1 Microplate Reader H1M at an excitation λ of 530 nm and an emission λ of 590 nm. Media in the absence of bacteria served as the negative control, and media with bacteria in the absence of compound served as the positive control. Percent inhibition was calculated as follows:

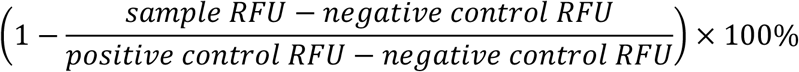

Percent inhibition was input into PRISM GraphPad to calculate IC_50_ values by plotting as a non-linear regression curve.

#### Cytotoxicity Assay

Vero cells (derived from African Green Monkey kidney epithelial tissue) were grown in Dulbecco’s Modified Eagle Medium (DMEM) supplemented with 10% fetal bovine serum (FBS), 10 mM HEPES buffered saline, and 2 mM L-glutamine. 5,000 cells per well were seeded into a 96-well plate containing two-fold titrations of compounds. The concentration of DMSO was maintained at 0.5% for all wells to avoid toxicity. The plates were incubated for 72 hours, at which point they were acclimatized to room temperature for 30 minutes. 25 μL of the substrate/buffer solution from the Promega CellTiter-Glo Luminescent Cell Viability Assay kit was added to each well. The plates were shaken at a low orbital speed for 3 minutes and let rest at room temperature for 10 minutes, after which luminescence was measured using a BioTek Synergy H1 Microplate Reader H1M. The positive control was defined as wells containing cells without compounds. Percent viability was calculated as follows:

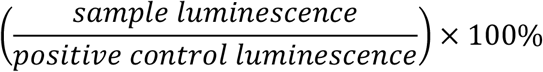

Percent viability was input into PRISM GraphPad to calculate LD_50_ values by plotting as a non-linear regression curve.

## ASSOCIATED CONTENT

The Supporting Information is available free of charge at.

Additional biological protocol, data on the activity against other mycobacteria and non-mycobacteria species, additional chemistry procedures including, ^1^H, ^13^C NMR, HRMS, and HPLC purity analysis, HPLC trace for selected compounds etc. are provided in the Supporting Information.

## AUTHOR INFORMATION

### Notes

The authors declare no competing financial interest.

## ACKNOWLEDGMENT

Support was provided by the University of Mississippi School of Pharmacy. We acknowledge the Department of Chemistry and Biochemistry and the Glycoscience Center of Research Excellence (GLYCORE, funded by National Institutes of Health 1P20GM130460-01A1-7936) Analytical and Biophysical Chemistry Research Core for the acquisition of high-resolution mass spectrometry (HRMS) data. Authors acknowledge the support of NIAID Anti-mycobacterial Target or Mechanism Identification Contract (AToMIc) and Dr. Richard Slayden of the Department of Microbiology, Immunology and Pathology, Colorado State University, Fort Collins, Colorado for the experimental data on the drug-resistant strains. For the preliminary ADME results, we also acknowledge Dr. Suresh P. Sulochana of the University of Mississippi School of Pharmacy’s Center of Biomedical Research Excellence (COBRE) CDMPK.

