## Supporting Information for "Discovery, Synthesis, and Optimization of 1,2,4-Triazolyl Pyridines Targeting *Mycobacterium tuberculosis*"

1. **Biological protocols**

### 1.1 Antimicrobial susceptibility test against NTM strains (data from NIAID AToMIc)

**Table S1. Activities of selected analogs against *M. smegmatis***

| MIC ( μg/mL) | |
|---|---|
| **entry** | *M. smegmatis*[a] |
| **1** | >60 |
| **4** | >60 |
| **5** | >60 |
| **8** | >60 |
| **9** | >60 |
| **Amikacin** | 2 |

[a] MIC was determined by using the MABA.

**Table S2. Activities of selected analogs against *M. smegmatis*, *M. avium,* and *M. abscessus* strains**

| MIC ( μg/mL) | | |
|---|---|---|
| **entry** | *M. avium*[a] | *M. abscessus*[a] |
| **1** | >60 | >60 |
| **4** | >60 | >60 |
| **5** | >60 | >60 |
| **8** | >60 | >60 |
| **9** | >60 | >60 |
| **Amikacin** | ND | 4 |
| **Clarithromycin** | 0.06 | 1 |
| **Rifampicin** | 0.06 | ND |

**[a]** The susceptibility of *M. avium* (ATCC 700891–MAC 101) and *M. abscessus* to **1**, **4**, **5**, **8**, and **9** were determined using the microbroth dilution method. MIC were determined using optical density (OD600nm) and confirmed using a calorimetric growth indicator (i.e., Alamar Blue).

**Materials and media procedures for MIC determination against NTM strains**

Materials:

Middlebrook 7H9 Broth - Difco (Becton Dickinson) – Cat# 271310

Cation-adjusted Meuller-Hinton Broth – BBL (Becton Dickinson) – Cat# 211438

BSA - Sigma - Catalog #, A2153

Dextrose – Fisher – Catalog #, D16-500

Catalase - Sigma - Catalog #, C-40

Assays may be run plus or minus 0.05 %(v/v) tween-80.

Polysorbate-80 (Tween 80), Sigma # P-5188 (prepared as 5% (v:v) stock in $DDH_2O$ and passed through 0.2 µm filter).

Procedures for making 1 L (7H9):

1. 4.7 g of 7H9 powder
2. Place 7H9 powder in 1L flask with stir bar
3. Add 2 mL glycerol
4. Bring final volume to 900 mL with Milli-Q-water or equivalent
5. Place on stir plate and mix until dissolved
6. Autoclave

Procedures for making 100mL ADC:

1. 5g bovine serum albumin
2. 2g dextrose
3. 3mg catalase
4. Bring final volume to 100 mL Milli-Q-water or equivalent
5. Stir to dissolve
6. Filter sterilize (0.22 µM)

Procedures for making 1 L (7H9 complete medium):

1. Add 100 mL ADC (albumin, dextrose, catalase) to 900 mL 7H9 broth
2. Add 10 mL of sterile 5% [v:v] polysorbate 80 (tween 80) to yield 0.05% [v:v] tween 80 final
3. Do sterility test by incubating the bottles of 7H9 complete media for 24 hours at 37°C
4. Hold media at room temp < 2 months

**Bacteria inoculant preparations**

*NTM* strains:

*M. abscessus*; ATCC19977

*M. avium*; ATCC 700891 – MAC 101

Target adjusted final Concentration of NTM strains: 5 x $10^5$ CFU/mL

Verification of the inoculum:

NTMs are grown on solid medium and CFUs are taken from the plates and used to inoculate 7H9 broth with 0.05% tween-80 and grown at 35-37°C in ambient air until the optical density (OD) absorbance taken after days 2, 3, 5, 7, 12 and 14 is an (OD) 0.08 - 0.1 (0.5 McFarland Standard). The bacterial cell suspensions are then confirmed by preparing them in saline, matching the (OD) 0.08 - 0.1 (0.5 McFarland Standard). Plate 5 µL of the prepared inoculum and 100 µL of a 1:1000 dilution (5 µL into 4,995 µL of 7H9) of the prepared inoculum onto a MH or 7H9 medium (expect 50 CFU if the inoculum contained $5x10^5$CFU/mL).

Drug Preparation:

1. Compounds are made using 1.28 mg/mL in DMSO (or as recommended)
2. 20 µL of compound added to the first column of wells and serially diluted 1:2 drug dilutions by transferring 100 µL from well to well (L to R) to establish a test range 64-0.062 µg/mL
3. Finish the dilution sets in column 10 by mixing and retaining the 100 µl in the wells.

Assay Plate Preparation:

1. MIC testing is performed by microbroth dilution method using cation-adjusted Mueller-Hinton broth.
2. Add 100 µl of NTM bacterial suspension in all the wells except the media only control wells.
3. QC agents specific for each organism 1) bacteria only negative control 2) media only negative control 3) (CLA) clarithromycin and amikacin (AMI) positive drug control 4) optional *E. coli* control.
4. Seal Assay plates and incubate at 37ºC. Read plates at $OD_{600}$ per CLSI guidelines for NTMs (day 7). After the $OD_{600}$ reading, add 10 µL Alamar Blue dye or resazurin (7-Hydroxy-3*H*-phenoxazin-3-one 10-oxide) to each analytical well.
5. Scan all assay plates after Alamar Blue dye on a flatbed color scanner.

**1.2 Antimicrobial susceptibility test against non-mycobacteria strains** (data from NPNCR)

**Table S3**: Activity against non-mycobacteria strains

| **entry[a]** | **$IC_{50}$ µg/mL** | | | | | | | |
|---|---|---|---|---|---|---|---|---|
| | MRSA | *E. coli* | *P. aeruginosa* | *K. pneumoniae* | VRE | *A. fumigatus* | *C. albicans* | *C. neoforman* |
| **1** | >20 | >20 | >20 | >20 | >20 | >20 | >20 | 4.18 |
| **8** | >20 | >20 | >20 | >20 | >20 | >20 | >20 | >20 |
| **9** | >20 | >20 | >20 | >20 | >20 | >20 | >20 | >20 |

| | | | | | | | | |
|---|---|---|---|---|---|---|---|---|
| **18** | >20 | >20 | >20 | >20 | >20 | >20 | >20 | >20 |
| **20** | >20 | >20 | >20 | >20 | >20 | >20 | >20 | >20 |
| **21** | >20 | >20 | >20 | >20 | >20 | >20 | >20 | >20 |
| **24** | >20 | >20 | >20 | >20 | >20 | >20 | >20 | >20 |
| **34** | >20 | >20 | >20 | >20 | >20 | >20 | >20 | >20 |
| **36** | >16.6 | >20 | >20 | >20 | >20 | >20 | >20 | ND |
| **44** | >20 | >20 | >20 | >20 | >20 | >20 | >20 | >20 |
| **Cefotaxime** | 8.6 | - | 20 | - | - | ND | ND | ND |
| **Meropenem** | 3.5 | 3.6 | 5.5 | - | - | ND | ND | ND |
| **Methicillin** | 24 | - | - | - | - | ND | ND | ND |
| **Amphotericin B** | ND | ND | ND | ND | ND | 0.1 | 0.63 | <0.1 |

[a]The activity of compounds was tested against drug resistant bacterial including Methicillin-resistance *Staphylococcus aureus* (MRSA), *Escherichia coli*, *Pseudomonas aeruginosa*, *Klebsiella pneumoniae* and Vancomycin-resistant *Enterococci faecalis (VRE)*. The susceptibility testing was carried out at three concentrations (20, 4, and 0.8 µg/mL). Compounds were also tested against two fungi strains namely A. *fumigatus* and *C. neoformans*. ND = Not determined.

All microbial strains used in this study are obtained from the American Type Culture Collection (ATCC, Manassas, VA). The efficacy of compounds were evaluated against *Candida albicans* ATCC 90028, *Cryptococcus neoformans* ATCC 90113, *Aspergillus fumigatus* ATCC 204305, methicillin-resistant Staphylococcus aureus ATCC 1708 (MRS), *Escherichia coli* ATCC 2452, *Pseudomonas aeruginosa* ATCC BAA-2018, *Klebsiella pneumoniae* ATCC 2146 and vancomycin-resistant *Enterococcus faecium* (VRE) ATCC 700221. Susceptibility testing was performed using a modified version of the CLSI methods.[1–3]. Pure compounds were tested at 20, 4, 0.8 µg/mL in 384 well plate using high-throughput screening. The incubation broth RPMI 1640 (2% dextrose/0.03% glutamine/MOPS at pH 6.0) was used for *C. albicans*, Sabouraud Dextrose for *C. neoformans*, cation-adjusted Mueller-Hinton at pH 7.0 for MRS, VRE, *E. coli*, *K. pneumoniae* and *P. aeruginosa*, and RPMI 1640 broth (2% dextrose, 0.03% glutamine, buffered with 0.165M MOPS at pH 7.0) for *A. fumigatus* to afford recommended inocula as per CLSI protocol. 5% Alamar Blue™ was added in *A. fumigatus*, VRE and MRS. Drug controls for bacteria and fungi were included in each assay. All organisms were read, at either 530nm or 544ex/590em for *A. fumigatus*, VRE and MRS, using the Bio-Tek plate reader prior to and after incubation: MRS, VRE, *E. coli*, *K. pneumoniae* and *P. aeruginosa* at 35°C for 18-24h, *C. albicans* and *A. fumigatus* at 35°C for 48h and *C. neoformans* at 35°C for 68–72 h. The concentration of compounds for 50% growth inhibition ($IC_{50}$) was calculated using XLfit 4.2 software (IDBS, Alameda, CA) using fit model 201.

### 1.3 Protocol for activity against mono-resistant strains of *M. tuberculosis* (data from NIAID AToMIc)

MIC of compounds were determined against *Mtb* H37Rv and 3 clinically relevant resistant strains using the microbroth dilution method. MIC was determined using optical density and a calorimetric growth indicator. For optical density ($OD_{600nm}$), the MIC is considered the first concentration to inhibit growth when compared to untreated growth control. For the calorimetric growth indicator (i.e. Alamar Blue), the MIC is calculated as the first concentration for the observed color change from pink, indicating active growth to blue, indicating no active growth.

**Bacteria inoculant preparations**

***M. tuberculosis* strains**: (i) *Mtb* H37Rv, Lot VG 5/25/16 (ii) *Mtb* RIFr; RpoB(S450L); ATCC #35838VG 04/13/07 (iii) *Mtb* INHr; KatG(del); MS015 11/29/19 (iv) *Mtb* FQr: GyrA(D94K); MOX3 VG 03/15/07

Materials:

Middlebrook 7H9 Broth - Difco (Becton Dickinson) – Cat# 271310

Glycerol – Fisher –Catalog # G33-500

Bovine Serum Albumin (BSA) - Sigma - Catalog #, A2153

Dextrose – Fisher – Catalog #, D16-500

Catalase - Sigma - Catalog #, C-40

Assays may be run plus or minus 0.05 %(v/v) tween-80.

Polysorbate-80 (Tween 80), Sigma # P-5188 (prepared as 5% (v/v) stock in milli-Q water and passed through 0.2 µm filter).

BSA - Sigma - Catalog #, A2153

Dextrose – Fisher - Catalog #, D16-500

Catalase - Sigma - Catalog #, C-40

Procedures for making 1 L (7H9):

1. 4.7 g of 7H9 powder
2. Place 7H9 powder in 1L flask with stir bar
3. Add 2 mL glycerol
4. Bring final volume to 900 mL with Milli-Q-water or equivalent
5. Place on stir plate and mix until dissolved

Procedures for making 100mL ADC:

1. 5g bovine serum albumin
2. 2g dextrose
3. 3mg catalase

4. Bring final volume to 100 mL Milli-Q-water or equivalent
5. Stir to dissolve

<u>Procedures for making 1 L (7H9 complete medium):</u>

1. Add 100 mL ADC (albumin, dextrose, catalase) to 900 mL 7H9 broth
2. Add 10 mL of sterile 5% [v:v] polysorbate 80 (tween 80) to yield 0.05% [v:v] tween 80 final
3. Filter sterilize (0.22 µM)
4. Do sterility test by incubating the bottles of 7H9 complete media for 24 hours at 37°C
5. Hold media at room temp < 2 months

**Bacteria inoculant preparations**

*M. tuberculosis* strains:

*Mtb* H37Rv, Lot VG 5/25/16

*Mtb* $RIF^r$; $RpoB^{(S450L)}$; ATCC #35838VG 04/13/07

*Mtb* $INH^r$; $KatG^{(del)}$; MS015 11/29/19

*Mtb* FQr: $GyrA^{(D94K)}$; MOX3 VG 03/15/07

Target adjusted final Concentration of *M. tuberculosis*: 5 x $10^5$ CFU/mL

Verification of the inoculum:

Plate 5 µL of the prepared inoculum and 100 µL of a 1:1000 dilution (5 µL into 4,995 µL of 7H9) of the prepared inoculum onto a 7H11-OADC medium (expect 50 CFU if the inoculum contained $5x10^5$CFU/mL).

Drug Preparation:

4. After labeling plates (see example below), add 50 µL of 100% DMSO to each interior well in columns 3 through 11 (see section Plate Layout).
5. Add 100 µL of a test material/compound at the starting concentration to each well in column 2 designated as the start of each dilution series (see section Plate Layout).
6. Make 1:2 drug dilutions by transferring 50 µL from well to well (L to R). Change tips between dilutions.
7. Finish the dilution sets in column 10 by mixing and retaining the 100 µl in the wells. Note the wells in column 11 do not get any drugs, only 50 µL of diluent (diluent only control).

Assay Plate Preparation:

1. After labeling plates (see example below), add 100 µL of media into *all* wells.
2. Transfer 2.5 µL of each drug dilution from the Drug Plate to the corresponding wells in the Assay plates.
3. Seal Assay plates and incubate at 37ºC.
4. Read plates at $OD_{600}$ on day 7/8.
5. After the $OD_{600}$ reading, add 10 µL Alamar Blue dye to each analytical well.

6. Scan all assay plates after Alamar Blue dye on a flatbed color.

#### 1.4 *In vitro* ADME (data from Cyprotex)

**Table S4.** Caco-2 permeability of compound 20

| **entry** | test tonc (µM) | assay duration (hr) | mean A → B Papp $10^{-6}$ cm/s | mean B → A Papp $10^{-6}$ cm/s | efflux Ratio | comment |
|---|---|---|---|---|---|---|
| **20** | 10.0 | 2.00 | 0.456 | 0.699 | 1.54 | Test compound not detectable in receiver compartment of replicate 2 A2B [a] |
| **Atenolol** | 10.0 | 2.00 | 0.198 | 0.401 | 2.03 | Low Permeability Control |
| **Talinolol** | 10.0 | 2.00 | 0.138 | 10.70 | 77.5 | P-gp Efflux Control |
| **Antipyrine** | 10.0 | 2.00 | 38.2 | 42.9 | 1.12 | High Permeability Control |

[a]Low recovery may be due to poor aqueous solubility, poor stability, and/or non-specific binding (high lipophilicity) of the compound. Permeability rates for compounds with low/poor post assay recovery may be underestimated.

**Table S5**. Experimental summary for turbidimetric solubility

| Test Article | Test conc. (µM) | Medium | Incubation | Reference Compounds |
|---|---|---|---|---|
| TA | 1.6, 3.1, 6.25, 12.5, 25, 50, 100, 200 µM | PBS (pH 7.4) | 2 hr 37°C | Reserpine Tamoxifen Verapamil |

**Experimental procedure for solubility:** Serial dilutions of test articles were prepared in DMSO at 100x the final concentration. Test article solutions were then diluted 100-fold into buffer in a 96-well plate and mixed. After time, the presence of precipitate was then detected by turbidity (absorbance at 540 nm). An absorbance value of greater than 'mean + 3x standard deviation of the blank' (after subtracting the background) was considered an indication of turbidity. For brightly colored compounds, a visual inspection of the plate was performed to verify the solubility limit determined by UV absorbance.

**Data Analysis:** The solubility limit was reported as the highest experimental concentration with no evidence of turbidity.

**Table S6**: Experimental summary for plasma protein binding assay

| Test Article | Test conc. | Plasma Species | Incubation | Reference Compounds |
|---|---|---|---|---|
| TA | 5 µM | Mouse, Human | 4 hr<br>37 °C | Warfarin |

**Experimental procedure for plasma protein binding:** Test article was added in duplicate to plasma (pH 7.4, ± 0.1, adjusted if necessary). This mixture was dialyzed in a RED device (Rapid Equilibrium Dialysis, Pierce) per the manufacturers' instructions against PBS and incubated on an orbital shaker. At the end of the incubation, aliquots from both plasma and PBS sides was collected, and was matrix-matrix matched with an appropriate amount of PBS and blank plasma, respectively. Acetonitrile (three volumes) containing an analytical internal standard (IS) was added to precipitate the proteins and release the test article. After centrifugation, the supernatant was transferred to a new plate and analyzed by LC-MS/MS to obtain peak area ratios (analyte/IS) for determining the fraction unbound.

**Data Analysis:** The extent of binding was reported as a fraction unbound ($f_u$) value which is calculated as detailed below:

fu= PF/PC

Where: fu = fraction unbound PC = Test compound in protein-containing compartment

PF = Test compound in protein-free compartment.

**Table S7:** Experimental summary plasma stability half-life determination

| Test Article | Test conc. | Test Species | Incubation | Reference Compounds |
|---|---|---|---|---|
| TA | 5 µM | Mouse, Human | 0, 15, 30, 60, and 120 min: (37 °C) | Warfarin (all)<br>Propantheline (all, except rat)<br>Enalapril (rat) |

**Experimental procedure for plasma stability:** The test article was incubated in singlicate with plasma (pH 7.4, ± 0.1, adjusted if necessary) at 37 ºC. At the indicated times, an aliquot was removed from each experimental reaction and mixed with ice-cold Stop Solution (methanol containing an analytical internal standard). Stopped reactions was kept on ice for at least ten minutes. The sample was centrifuged to remove precipitated protein, and the supernatants analyzed by LC-MS/MS to quantitate the remaining parent.

**Data Analysis:** Data was calculated as % parent remaining by assuming zero-minute time point peak area ratio (analyte/Internal Standard) as 100 % and dividing remaining time point peak area ratios by zero-minute time point peak area ratio. Data was subjected to fit first-order decay model to calculate slope and thereby half-life.

**Table S8**: Experimental summary for microsomal stability (Intrinsic Clearance)

| Test Article | Test Conc. | Microsome Species | Protein Conc. | Incubation | Analytical Method |
|---|---|---|---|---|---|
| TA | 1 μM | Mouse, Human | 0.5 mg/mL | 0, 5, 15, 30 and 45 min (37°C) | LC-MS/MS |

**Experimental procedure for microsomal stability:** The test article was incubated in singlicate with liver microsomes at 37 ºC. The reaction contains microsomal protein in 100 mM potassium phosphate buffer (pH 7.4), 2 mM NADPH, 3 mM $MgCl_2$. A control was run for each test article omitting NADPH to detect NADPH-free degradation. At predetermined time points as mentioned above, an aliquot was removed from each experimental and control reaction and mixed with an equal volume of ice-cold acetonitrile containing internal standard to stop the reaction and precipitate proteins. The samples were centrifuged to remove precipitated protein, and the supernatants were then analyzed by LC-MS/MS to quantify % parent remaining.
**Data Analysis:** Data was calculated as % parent remaining by assuming zero-minute time point peak area ratio (analyte/IS) as 100% and dividing remaining time point peak area ratios by zero-minute time point peak area ratio. Data was subjected to fit first-order decay model to calculate slope and thereby half-life. Intrinsic clearance was calculated from the half-life and the human liver microsomal protein concentrations using following equations:

$t_{1/2} = \ln(2) / -k$
$CL_{int} = \ln(2) / (t_{1/2}\ [\text{microsomal protein}])$

Where

k = slope (elimination constant)
$CL_{int}$ = intrinsic clearance
$t_{1/2}$ = half-life

### 2. Chemistry Protocols

#### 2.1 General experimental conditions

All solvents and reagents were purchased from standard commercial vendors and used without further purification. Synthetic reactions were monitored using thin-layer chromatography (TLC) (Sorbtech silica XG TLC plates) and visualized under UV at 254 nm or with appropriate staining. Purification was undertaken using medium pressure liquid chromatography (MPLC) on Biotage

Isolera One with Biotage SNAP 10g–50g cartridges. NMR spectra were recorded on a Bruker Avance-500 or Bruker Avance-400 spectrometers at 298.15 K. Chemical shifts are reported in ppm using deuterated solvents ($CDCl_3$, $CD_3OD$, or DMSO-$d_6$) for $^1$H and $^{13}$C NMR. $CDCl_3$ (δ = 77.16 ppm), $CD_3OD$ (δ = 49.00 ppm) or DMSO-$d_6$ (δ = 39.52 ppm) were used as internal standards for $^{13}$C NMR. For $^1$H NMR, $CDCl_3$ (δ = 7.26 ppm), $CD_3OD$ (δ = 3.31 ppm) or DMSO-$d_6$ (δ = 2.50 ppm) or TMS (δ = 0 ppm) were used as internal standards. Data were reported as: s = singlet, br = broad singlet, d = doublet, t = triplet, q = quartet, p = pentet, m = multiplet, b = broad, ap = apparent; coupling constants, *J*, in Hz.For High-resolution mass spectrometry (HRMS), quadruple-TOF was used to obtain the data both in positive and negative modes. Attenuated total reflectance Infra-red (ATR-IR) was taken using an Agilent Technologies Cary 600 series FTIR Spectrometer. Purity (>95%) were determined using a Dionex Ultimate 3000 UPLC system (Thermo Fisher Scientific). Melting points were determined using an OptiMelt automated melting point system.

**2.2 Synthetic procedure for intermediates**

**Scheme S1.** Synthesis of compound **54**

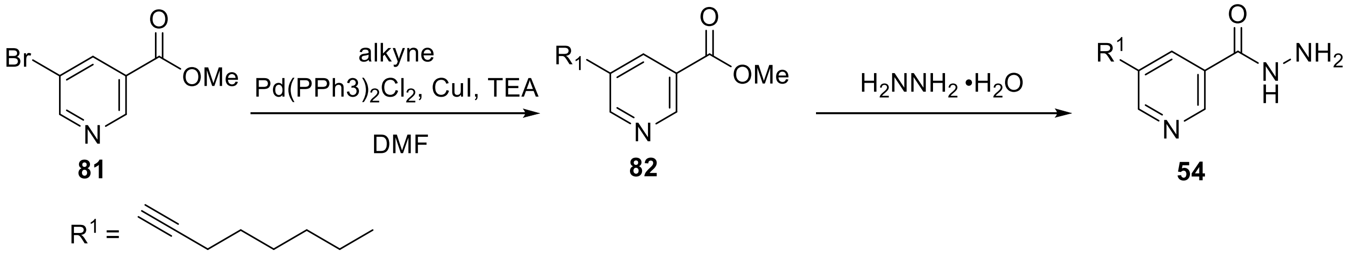

**General procedure B for Sonogashira coupling of bromide with corresponding alkyne**

A mixture of the appropriate aryl bromide (A) (1.0 equiv), copper(I) iodide (0.1.0 equiv), and bis(triphenylphosphine)palladium (II) chloride (0.05 equiv) in DMF (0.2 molar) was purged with argon for 10 min. To the above mixture (A) was added a solution of corresponding alkyne (1.0 equiv) that was previously stirred with triethylamine (2.0 equiv) under argon. The resultant mixture was then stirred at room temperature overnight. After completion of reaction as determined by TLC, the reaction mixture was diluted with a saturated solution of $NaHCO_3$ (10 mL) and filtered through a celite pad. The filtrate was then extracted twice with diethyl ether, washed with water and brine, dried with anhydrous $Na_2SO_4$, and purified with using MPLC.

**Methyl 5-(oct-1-yn-1-yl)nicotinate (82) .** Methyl 5-bromonicotinate, **82** (1.0 g, 1.0 equiv, 9.3 mmol), **81** was coupled with oct-1-yne (1.4 mL, 1.0 equiv, 9.3 mmol) using general procedure B. Final product was purified by MPLC (mobile phase: 2–20% EtOAc/hexane). Yield: 92% (2.1 g) as a yellow liquid. $^1$H NMR (500 MHz, $CDCl_3$) δ 9.05 (d, *J* = 2.1 Hz, 1H), 8.74 (d, *J* = 2.0 Hz, 1H), 8.26 (t, *J* = 2.1 Hz, 1H), 2.42 (t, *J* = 7.2 Hz, 2H), 1.65–1.55 (m, 2H), 1.49–1.38 (m, 2H), 1.37–1.25 (m, 4H), 0.90 (t, *J* = 6.8 Hz, 3H). $^{13}$C NMR (101 MHz, $CDCl_3$) δ 165.4, 155.7, 148.8, 139.4, 125.5, 121.3, 95.5, 76.5, 52.5, 31.4, 28.7, 28.5, 22.6, 19.5, 14.1. HRMS (ESI): m/z [M + H] Calcd for $[C_{15}H_{19}NO_2 + H]^+$ 246.1494, found 246.1476.

**General procedure C for synthesis of acyl hydrazide**

To the methyl or ethyl ester of the appropriate carboxylic acid (1.0 equiv) dissolved in EtOH (0.2 molar) was added 64% solution hydrazine hydrate (2.0 equiv) gradually and reflux for 5 h. Upon completion of reaction as determined by TLC, the reaction mixture was brought to room temperature and the precipitated product was filtered and rinsed with methanol and n-hexanes to obtain a white crystalline solid. Product was used for the next step without further purification.

**5-(Oct-1-yn-1-yl) nicotinohydrazide (54).** Methyl 5-(oct-1-yn-1-yl)nicotinate, **82** (3 g, 1.0 equiv, 12.23 mmol) was reacted with 64% hydrazine hydrate (1.23 g, 2.0 equiv, 24.46 mmol) using general procedure C**.** Yield: 93% (2.80 g) as white solid; mp 148–150 °C $^{1}$H NMR (400 MHz, $CD_3OD$) δ 8.81 (d, *J* = 2.1 Hz, 1H), 8.64 (d, *J* = 2.1 Hz, 1H), 8.14 (t, *J* = 2.1 Hz, 1H), 2.48 (t, *J* = 7.0 Hz, 2H), 1.67–1.57 (m, 2H), 1.54–1.42 (m, 2H), 1.42–1.30 (m, 4H), 0.97–0.88 (m, 3H). $^{13}$C NMR (101 MHz, $CD_3OD$) δ 166.6, 154.7, 147.2, 138.7, 130.3, 122.9, 96.6, 77.2, 32.5, 29.7, 29.5, 23.6, 20.0, 14.4. HRMS (ESI): m/z $[M - H]^-$ Calcd for $[C_{14}H_{19}N_3O + H]^+$ 245.1606, found 245.1586.

**Synthesis of compound 85–87; Sonogashira coupling of bromide with corresponding alkynes using general procedure B**

**Scheme S2.** Synthesis of **85–87**

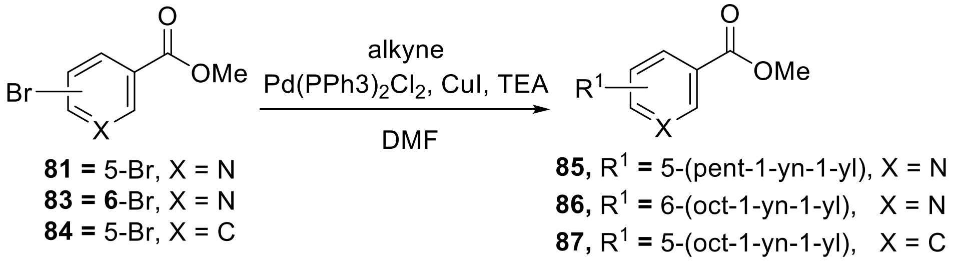

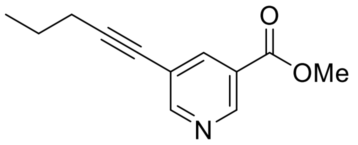

**Methyl 5-(pent-1-yn-1-yl) nicotinate (85)** Methyl 5-bromonicotinate **81** (1.0 g, 1.0 equiv, 5 mmol) was coupled with pent-1-yne (0.6 mL, 1.25 equiv, 6 mmol) using general procedure B. Product was purified by MPLC (mobile phase: 0–30% EtOAc/hexane). Yield: 90% (0.821 g) as a brown liquid. $^{1}$H NMR (400 MHz, $CDCl_3$) δ 8.97 (d, *J* = 2.1 Hz, 1H), 8.66 (d, *J* = 2.1 Hz, 1H), 8.16 (t, *J* = 2.1 Hz, 1H), 3.86 (s, 3H), 2.33 (t, *J* = 7.0 Hz, 2H), 1.57 (p, *J* = 7.3 Hz, 2H), 0.97 (t, *J* = 7.4 Hz, 3H). $^{13}$C NMR (101 MHz, $CDCl_3$) δ 165.2, 155.5, 148.7, 139.2, 125.4, 121.1, 95.1, 76.6, 52.4, 21.85, 21.3, 13.4. HRMS (ESI): m/z $[M + H]^+$ Calcd for $[C_{12}H_{13}NO_2 + H]^+$ 204.1024, found 204.1049.

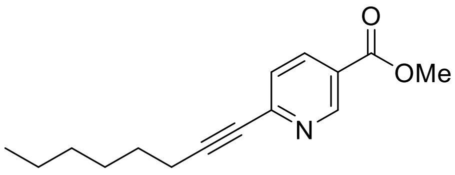

**Methyl 6-(oct-1-yn-1-yl)nicotinate (86).** Methyl 6-bromonicotinate **83** (1.0 g, 1.0 equiv, 5 mmol) was coupled with oct-1-yne (0.8 mL, 1.2.0 equiv, 6 mmol) using general procedure B. Final product was purified by MPLC (mobile phase: 2–10 % EtOAc/hexane). Yield 80% (0.907 g) as a yellow liquid. $^{1}$H NMR (400 MHz, $CDCl_3$) δ 9.07 (dd, *J* = 2.2, 0.9 Hz, 1H), 8.15 (dd, *J* = 8.2, 2.2 Hz, 1H), 7.37 (dd, *J* = 8.2, 0.9 Hz, 1H), 3.88 (s, 3H), 2.41 (t, *J* = 7.1 Hz, 2H), 1.63–1.53 (m, 2H),

1.46–1.34 (m, 2H), 1.31–1.21 (m, 4H), 0.84 (t, *J* = 6.9 Hz, 3H). $^{13}$C NMR (101 MHz, $CDCl_3$) δ 165.4, 150.9, 147.7, 137.1, 126.3, 124.1, 94.8, 80.3, 52.4, 31.3, 28.7, 28.2, 22.5, 19.5, 14.1. HRMS (ESI): m/z $[M + H]^+$ Calcd for $[C_{15}H_{19}NO_2 + H]^+$ 246.1494, found 246.1476.

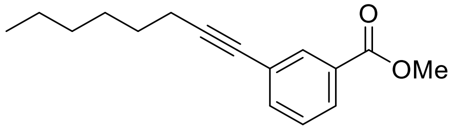

**Methyl 3-(oct-1-yn-1-yl)benzoate (87).** Methyl 3-bromobenzoate, **84** (1.0 g, 1.0 equiv, 4.65 mmol), was coupled with oct-1-yne (0.512 g, 687 µL, 1.0 equiv, 4.65 mmol) using general procedure B. Final product was purified by MPLC (mobile phase: 0–5% EtOAc/hexane) Yield: 85% (0.968 g) as a brown liquid. $^{1}$H NMR (400 MHz, $CDCl_3$) δ 8.06 (t, *J* = 1.8 Hz, 1H), 7.92 (dt, *J* = 7.8, 1.5 Hz, 1H), 7.55 (dt, *J* = 7.7, 1.5 Hz, 1H), 7.35 (t, *J* = 7.8 Hz, 1H), 3.91 (s, 3H), 2.40 (t, *J* = 7.1 Hz, 2H), 1.66–1.54 (m, 2H), 1.51–1.40 (m, 1H), 1.40–1.25 (m, 2H), 0.90 (t, *J* = 6.9 Hz, 3H). $^{13}$C NMR (101 MHz, $CDCl_3$) δ 166.7, 135.8, 132.8, 130.4, 128.6, 128.4, 124.7, 91.7, 79.8, 52.3, 31.5, 28.8, 28.7, 22.7, 19.5, 14.2. HRMS (ESI): m/z $[M + H]^+$ Calcd for $[C_{16}H_{20}O_2 + H]^+$ 245.1542, found 245.1542.

**Williamson ether synthesis of compound 89**

**Scheme S3.** Synthesis of compound **89**

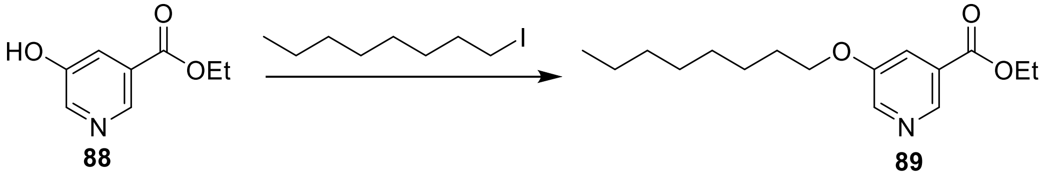

**Ethyl 5-(octyloxy) nicotinate (89).** To ethyl 5-hydroxynicotinate, **88** (500 mg, 1.0 equiv, 2.99 mmol) dissolved in acetonitrile (15 mL) was added $Cs_2CO_3$ (1170 mg, 1.2 equiv, 3.59 mmol) and *sec*-Octyl iodide (790 mg, 594 µL, 1.1.0 equiv, 3.29 mmol). The reaction mixture was stirred for 18 h 5 mL of water was added to the reaction mixture and then extracted three times with 50 mL of EtOAc. The combined organic phase was concentrated under vacuum and the crude was purified by MPLC (5–15% EtOAC/hex) to obtain a clear liquid product. Yield: 67% (563 mg). $^{1}$H NMR (400 MHz, $CDCl_3$) δ 8.78 (d, *J* = 1.7 Hz, 1H), 8.43 (d, *J* = 2.9 Hz, 1H), 7.72 (dd, *J* = 2.9, 1.7 Hz, 1H), 4.39 (q, *J* = 7.2 Hz, 2H), 4.02 (t, *J* = 6.5 Hz, 2H), 1.79 (dq, *J* = 7.9, 6.5 Hz, 2H), 1.50–1.21 (m, 13H), 0.91–0.82 (m, 3H). $^{13}$C NMR (101 MHz, $CDCl_3$) δ 165.4, 155.0, 142.8, 142.6, 126.7, 120.7, 68.6, 61.5, 31.8, 29.3, 29.2, 29.0, 25.9, 22.6, 14.3, 14.1. HRMS (ESI): m/z $[M + H]^+$ Calcd for $[C_{16}H_{25}NO_3Cs + H]^+$ 412.0889, found 412.0880.

**Synthesis of acyl hydrazide 61i–vi using general procedure C**

**Scheme S4.** Synthesis of acyl hydrazide **61i–vi**

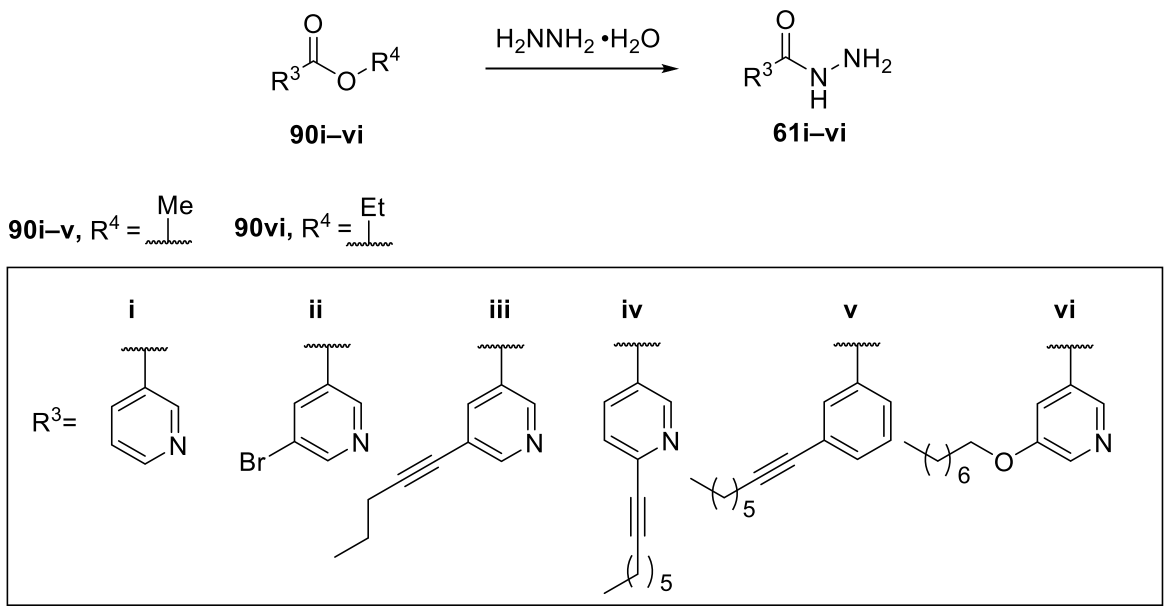

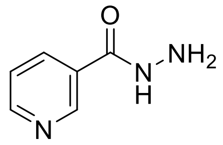

**Nicotinohydrazide (61i).** Methyl nicotinate **90i** (300 mg, 1.0 equiv, 2.19 mmol) was reacted with hydrazine hydrate (219 mg, 2.0 equiv, 4.38 mmol) using general procedure C. Yield: 99% (298 mg) as white solid; mp 159–161 °C. $^1$H NMR (400 MHz, DMSO-d6) δ 9.96 (s, 1H), 8.99–8.94 (m, 1H), 8.71–8.65 (m, 1H), 8.19–8.11 (m, 1H), 7.53–7.45 (m, 1H), 4.57 (s, 2H). $^{13}$C NMR (101 MHz, DMSO-d6) δ 164.3, 151.8, 148.1, 134.7, 128.9, 123.5. HRMS (ESI): m/z [M - H]$^-$ Calcd for [$C_6H_7N_3O$ - H]$^-$ 136.0511, found 136.0540.

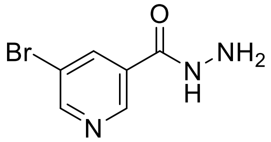

**5-Bromonicotinohydrazide (61ii).** Methyl 5-bromonicotinate, **90ii** (1.0 g, 1.0 equiv, 5 mmol) was reacted with 64% hydrazine hydrate (500 mg, 2.0 equiv, 9 mmol) using general procedure C. Yield: 90% (916 mg) as white solid; mp 188–190 °C. $^1$H NMR (400 MHz, DMSO-d6) δ 10.07 (s, 1H), 9.28–8.57 (m, 2H), 8.36 (s, 1H), 4.62 (s, 2H). $^{13}$C NMR (101 MHz, DMSO-d6) δ 162.8, 152.4, 146.7, 137.1, 130.6, 120.1. HRMS (ESI): m/z [M - H]$^-$ Calcd for [$C_6H_6BrN_3O$ - H]$^-$ 215.9596, found 215.9545.

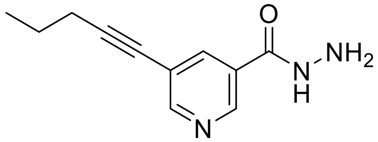

**5-(Pent-1-yn-1-yl)nicotinohydrazide (61iii).** Methyl 5-(pent-1-yn-1-yl)nicotinate, **90iii** (380 mg, 1.0 equiv, 1.87 mmol) was reacted with 64% hydrazine hydrate (187 mg, 2.0 equiv, 3.74 mmol) using general procedure C. Yield: 67% (253 mg) as a white solid; mp 199–180 °C. $^1$H NMR (400 MHz, DMSO-d6) δ 9.99 (s, 1H), 8.88 (d, *J* = 2.1 Hz, 1H), 8.69 (d, *J* = 2.0 Hz, 1H), 8.13 (t, *J* = 2.1 Hz, 1H), 4.62–4.52 (m, 2H), 2.46 (t, *J* = 7.0 Hz, 3H), 1.59 (h, *J* = 7.3 Hz, 2H), 1.02 (t, *J* = 7.4

Hz, 3H). $^{13}$C NMR (101 MHz, DMSO-d6) δ 163.4, 153.4, 146.7, 136.6, 128.4, 119.9, 94.9, 77.0, 21.4, 20.6, 13.3. HRMS (ESI): m/z [M - H]$^-$ Calcd for [$C_{12}H_{13}NO_2$ + H] $^+$ 204.1137, found 204.1164.

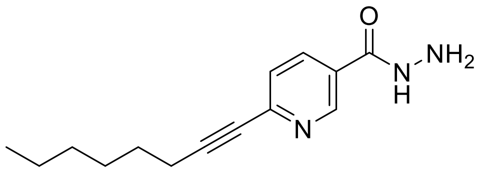

**6-Oct-1-yn-1-yl)nicotinohydrazide (61iv).** Methyl 6-(oct-1-yn-1-yl)nicotinate, **90iv** (345 mg, 1.0 equiv, 1.41 mmol) was reacted with 64% hydrazine hydrate (141 mg, 2.0 equiv, 2.81 mmol) using general procedure C. Yield: 54% (187 mg) as white solid; mp 70–72 °C. $^1$H NMR (400 MHz, DMSO-d6) δ 10.64 (s, 1H), 8.90 (s, 1H), 8.24–8.04 (m, 1H), 7.54 (d, *J* = 8.1 Hz, 1H), 2.54–2.43 (m, 3H), 1.62–1.51 (m, 2H), 1.48–1.37 (m, 2H), 1.37–1.22 (m, 4H), 0.94–0.80 (m, 3H). $^{13}$C NMR (101 MHz, DMSO-d6) δ 165.0, 133.6, 129.6, 128.7, 126.4, 123.4, 91.4, 80.0, 30.7, 28.1, 28.0, 22.0, 18.6, 13.9. HRMS (ESI): m/z [M - H]$^-$ Calcd for [$C_{14}H_{19}N_3O$ - H]$^-$ 244.1450, found 244.1429.

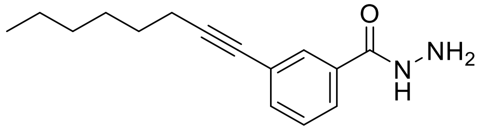

**3-(Oct-1-yn-1-yl) benzohydrazide (61v)**. Methyl 3-(oct-1-yn-1-yl)benzoate, **90v** (404 mg, 1.0 equiv, 1.65 mmol) was reacted with 64% hydrazine hydrate (166 mg, 2.0 equiv, 3.31 mmol) using general procedure C. Yield: 86% (347 mg) as a white solid; mp 122–124 °C. $^1$H NMR (400 MHz, DMSO-d6) δ 7.81(s, 1H), 7.79–7.74 (m, 1H), 7.54–7.46 (m, 1H), 7.41 (t, *J* = 7.7 Hz, 1H), 4.49 (s, 2H), 2.42 (t, *J* = 7.0 Hz, 2H), 1.61–1.48 (m, 2H), 1.47–1.36 (m, 2H), 1.36–1.20 (m, 4H), 0.94–0.79 (m, 3H). $^{13}$C NMR (101 MHz, DMSO-d6) δ 165.0, 133.6, 133.6, 129.7, 128.7, 126.4, 123.4, 91.4, 80.0, 30.8, 28.1, 28.0, 22.0, 18.6, 13.9. HRMS (ESI): m/z [M + H] Calcd for [$C_{15}H_{20}N_2OCs$ + H]$^+$ 377.0601, found 377.0630.

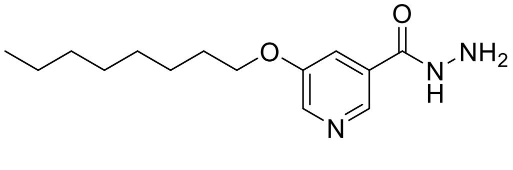

**5-(Octyloxy)nicotinohydrazide (61vi).** Ethyl 5-(octyloxy)nicotinate, **90vi** (380 mg, 1.0 equiv, 1.36 mmol) was reacted with 64% hydrazine hydrate (272 mg, 4.0 equiv, 5.44 mmol) using general procedure C. Yield: 88% (317 mg) as white solid; mp 97–99°C. $^1$H NMR (400 MHz, $CD_3OD$) δ 8.50 (d, *J* = 1.8 Hz, 1H), 8.34 (d, *J* = 2.9 Hz, 1H), 7.73 (dd, *J* = 2.9, 1.7 Hz, 1H), 4.09 (t, *J* = 6.4 Hz, 2H), 1.87–1.75 (m, 2H), 1.55–1.25 (m, 10H), 0.95–0.86 (m, 3H). $^{13}$C NMR (101 MHz, $CD_3OD$) δ 167.1, 157.0, 141.4, 140.6, 131.3, 120.9, 69.9, 33.0, 30.4, 30.4, 30.1, 27.0, 23.7, 14.4. HRMS (ESI): m/z [M - H]$^-$ Calcd for [$C_{14}H_{23}N_3O_2$ -H]$^-$ 264.1712, found 264.1698.

**Synthesis of 4,5-substituted-1,2,4-triazole-2-thiones (55–60, 62i–vi)**

**Scheme S5**. Synthesis of 4,5-substituted-1,2,4-triazole-2-thiones **55-60, 62i–vi**

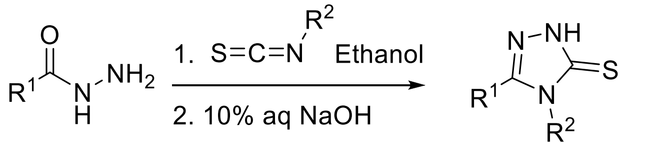

**54, 61i–vi** **55-60, 62i–vi**

**55, 63a–f**, **$R^2$** = $CH_3$-
**56**, **$R^2$** = $C_2H_5$-
**57**, **$R^2$** = $C_4H_9$-
**58**, **$R^2$** = Ph-
**59**, **$R^2$** = Bn-
**60**, **$R^2$** = cyclopropyl-

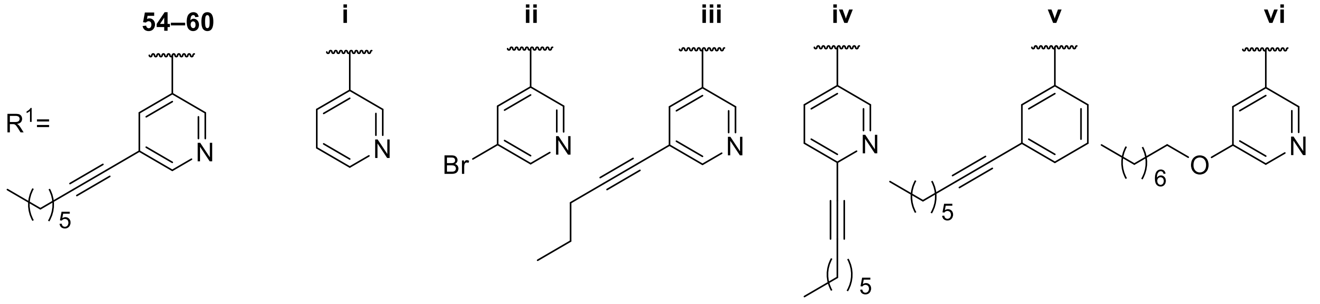

**General procedure D for synthesis of 4,5-substituted-1,2,4-triazole-2-thiones**

Step1: Acyl hydrazide (1.0 equiv) and corresponding isothiocyanate (1.0 equiv) dissolved in absolute ethanol (10 mL) was refluxed for 4 h. After completion of reaction, as confirmed by TLC, the solvent was removed under reduced pressure to obtain the corresponding 1,4-substituted thiosemicarbazides. Step 2: To the residue was then added 5 mL of 10% aqueous NaOH and stirred for 3 h at 60 °C. Upon completion, the reaction mixture was brought to room temperature and acidified with conc. HCl to a pH of 5–6. The precipitate formed was then filtered, rinsed with water (5 mL) to give the corresponding 4,5-substituted-1,2,4-triazole-2-thiol and was used without further purification. For compound that did not precipitate upon acidification, the reaction mixture was extracted (x 2) with EtOAc, the organic phase was then washed with brine, concentrated in *vacuo* and purified using MPLC.

**The synthesis of the following compounds was modified from the General Procedure C as needed.**

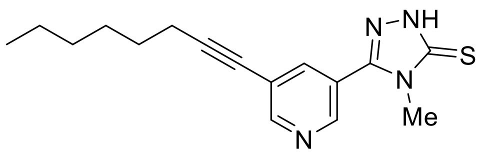

**4-Methyl-5-(5-(oct-1-yn-1-yl)pyridin-3-yl)-2,4-dihydro-3H-1,2,4-triazole-3-thione (55).** 5-(Oct-1-yn-1-yl) nicotinohydrazide, **54** (1.0 g, 1.0 equiv, 4 mmol) was reacted with isothiocyanatomethane (0.3 g, 0.3 mL, 1.0 equiv, 4 mmol) using general procedure C. Yield: 70% (0.810 g) as a white solid; mp 174–175 °C. $^1$H NMR (400 MHz, $CDCl_3$) δ 12.80 (s, 1H), 8.86–8.74 (m, 2H), 7.93 (t, $J$ = 2.1 Hz, 1H), 3.69 (s, 3H), 2.44 (t, $J$ = 7.2 Hz, 2H), 1.61 (q, $J$ = 7.4 Hz, 2H), 1.50–1.39 (m, 2H), 1.39–1.24 (m, 4H), 0.89 (t, $J$ = 6.8 Hz, 3H). $^{13}$C NMR (101 MHz, $CDCl_3$) δ 169.0, 154.1, 149.1, 146.6, 138.3, 122.1, 122.1, 96.9, 76.2, 32.4, 31.4, 28.7, 28.4, 22.6, 19.6, 14.2. HRMS (ESI): m/z [M + H] Calcd for $[C_{16}H_{20}N_4S + H]^+$ 301.1487, found 301.1482.

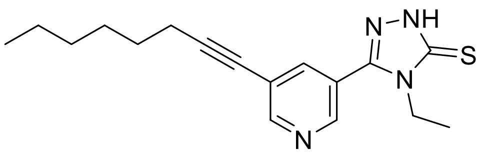

**4-Ethyl-5-(5-(oct-1-yn-1-yl)pyridin-3-yl)-2,4-dihydro-3H-1,2,4-triazole-3-thione (56).** 5-(Oct-1-yn-1-yl)nicotinohydrazide, **54** (80 mg, 1.0 equiv, 0.33 mmol) was reacted with isothiocyanatoethane (28 mg, 29 µL, 1.0 equiv, 0.33 mmol) using general procedure C. Yield: 73% (75 mg) as a white solid; mp 122–124 °C. $^1$H NMR (400 MHz, $CDCl_3$) δ 12.92 (s, 1H), 8.90–8.69 (m, 2H), 7.91 (t, *J* = 2.1 Hz, 1H), 4.17 (q, *J* = 7.2 Hz, 2H), 2.44 (t, *J* = 7.1 Hz, 2H), 1.61 (p, *J* = 7.2 Hz, 2H), 1.50–1.26 (m, 9H), 0.89 (t, *J* = 6.9 Hz, 3H). $^{13}$C NMR (101 MHz, $CDCl_3$) δ 168.1, 154.1, 148.8, 146.5, 138.4, 122.3,122.2, 96.9, 76.2, 40.4, 31.4, 28.7, 28.4, 22.6, 19.6, 14.2, 14.1. HRMS (ESI): m/z [M + H] Calcd for $[C_{17}H_{22}N_4S + H]^+$ 314.1565, found 314.1583.

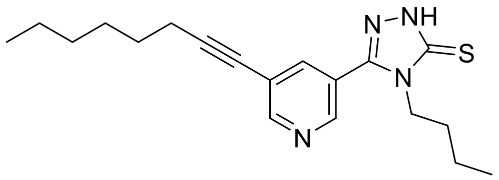

**4-Butyl-5-(5-(oct-1-yn-1-yl)pyridin-3-yl)-2,4-dihydro-3H-1,2,4-triazole-3-thione (57).** 5-(Oct-1-yn-1-yl)nicotinohydrazide, **54** (108 mg, 1.0 equiv, 440 µmol) was reacted with 1-isothiocyanatobutane (50.7 mg, 1.0 equiv, 440 µmol) using general procedure C. Yield: 88% (132 mg) as a white solid; mp 116–118 °C. $^1$H NMR (400 MHz, $CDCl_3$) δ 12.68 (s, 1H), 8.83–8.71 (m, 2H), 8.70 (s, 2H), 7.90 (s, 1H), 4.11 (t, *J* = 7.8 Hz, 2H), 2.44 (t, *J* = 7.1 Hz, 2H), 1.80–1.66 (m, 2H), 1.66–1.55 (m, 2H), 1.51–1.39 (m, 2H), 1.39–1.24 (m, 6H), 0.98–0.79 (m, 6H). $^{13}$C NMR (101 MHz, $CDCl_3$) δ 168.4, 154.1, 148.9, 146.6, 138.5, 122.4, 122.1, 96.8, 76.2, 44.9, 31.4, 30.5, 28.7, 28.4, 22.6, 19.8, 19.6, 14.1, 13.6. HRMS (ESI): m/z $[M + H]^+$ Calcd for $[C_{19}H_{26}N_4S + H]^+$ 343.1956, found 343.1965.

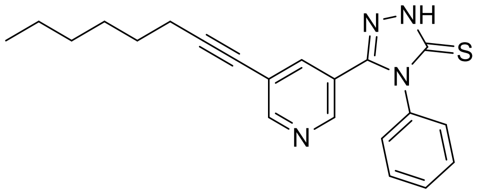

**5-(5-(Ooct-1-yn-1-yl)pyridin-3-yl)-4-phenyl-2,4-dihydro-3H-1,2,4-triazole-3-thione (58)** 5-(Oct-1-yn-1-yl)nicotinohydrazide, **54** (75 mg, 1.0 equiv, 0.31 mmol) was reacted with isothiocyanatobenzene (41 mg, 36 µL, 1.0 equiv, 0.31 mmol) using general procedure C. Final product was purified by MPLC (mobile phase: 20–50% EtOAc/hexane). Yield: 76% (84 mg) as a white solid; mp 168-170 °C. $^1$H NMR (400 MHz, $CDCl_3$) δ 12.5 (s, 1H), 8.6 (d, *J* = 2.0 Hz, 1H), 8.4 (d, *J* = 2.2 Hz, 1H), 7.7 (t, *J* = 2.1 Hz, 1H), 7.6–7.5 (m, 3H), 7.4–7.3 (m, 2H), 2.4 (t, *J* = 7.1 Hz, 2H), 1.6–1.5 (m, 2H), 1.5–1.2 (m, 5H), 0.9 (t, *J* = 6.8 Hz, 3H). $^{13}$C NMR (101 MHz, $CDCl_3$) δ 170.0, 153.6, 148.5, 146.4, 137.9, 133.8, 130.5, 130.3, 128.2, 121.7, 121.6, 96.2, 76.3, 31.4, 28.6, 28.4, 22.7, 19.5, 14.2. HRMS (ESI): m/z $[M-H]^-$ Calcd for $[C_{21}H_{22}N_4S - H]^-$ 631.1487, found 361.1478.

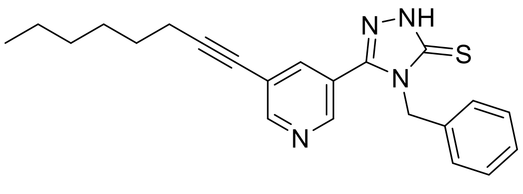

**4-Benzyl-5-(5-(oct-1-yn-1-yl)pyridin-3-yl)-2,4-dihydro-3H-1,2,4-triazole-3-thione (59).** 5-(Oct-1-yn-1-yl)nicotinohydrazide, **54** (100 mg, 1.0 equiv, 408 µmol) was reacted with (isothiocyanatomethyl)benzene (54 µL, 1.0 equiv, 408 µmol) . Final product was purified by MPLC (mobile phase: 20–50% EtOAc/hexane) using general procedure C. Yield: 99% (125 mg) as a white solid: mp 131–132 °C. $^{1}$H NMR (400 MHz, $CDCl_3$) δ 12.11 (s, 1H), 8.71 (d, *J* = 2.0 Hz, 1H), 8.53 (d, *J* = 2.2 Hz, 1H), 7.65 (s, 1H), 7.41–7.20 (m, 4H), 7.20–7.06 (m, 2H), 5.36 (s, 2H), 2.41 (t, *J* = 7.1 Hz, 2H), 1.64–1.55 (m, 2H), 1.48–1.38 (m, 2H), 1.38–1.27 (m, 4H), 0.91 (t, *J* = 6.9 Hz, 3H). $^{13}$C NMR (101 MHz, $CDCl_3$) δ169.2, 154.0, 149.6, 146.8, 138.7, 134.8, 129.2, 128.5, 127.2, 122.1, 121.8, 96.6, 76.1, 48.3, 31.4, 28.7, 28.4, 22.6, 19.6, 14.2. HRMS (ESI): m/z [M + H] Calcd for $[C_{22}H_{24}N_4Cs + H]^+$ 509.0776, found 509.0758.

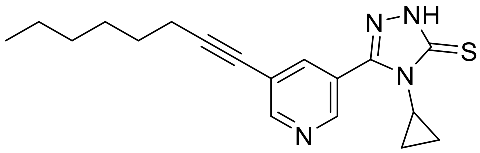

**4-Cyclopropyl-5-(5-(oct-1-yn-1-yl)pyridin-3-yl)-2,4-dihydro-3H-1,2,4-triazole-3-thione (60).** 5-(Oct-1-yn-1-yl)nicotinohydrazide, **54** (700 mg, 1.0 equiv, 2.85 mmol) was reacted with isothiocyanatocyclopropane (283 mg, 264 µL, 1.0 equiv, 2.85 mmol) using general procedure C. Final product was purified by MPLC (mobile phase: 20–50% EtOAc/hexane). Yield: 74% (685 mg) as a white solid: mp 133–135 °C. $^{1}$H NMR (400 MHz, $CDCl_3$) δ 13.14 (s, 1H), 8.93 (d, *J* = 2.2 Hz, 1H), 8.77 (d, *J* = 2.1 Hz, 1H), 8.02 (t, *J* = 2.1 Hz, 1H), 3.27–3.17 (m, 1H), 2.43 (t, *J* = 7.1 Hz, 2H), 1.60 (p, *J* = 7.1 Hz, 2H), 1.49–1.38 (m, 2H), 1.38–1.25 (m, 4H), 1.25–1.13 (m, 2H), 0.88 (t, *J* = 6.9 Hz, 3H), 0.83–0.74 (m, 2H). $^{13}$C NMR (101 MHz, $CDCl_3$) δ 170.2, 153.6, 149.7, 146.8, 138.2, 122.0, 121.7, 96.4, 76.3, 31.4, 28.7, 28.4, 27.1, 22.6, 19.6, 14.1, 9.8. HRMS (ESI): m/z [M + H] Calcd for $[C_{18}H_{22}N_4SCs + H]^+$ 459.0606, found 459.0620.

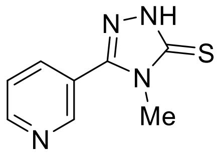

**4-Methyl-5-(pyridin-3-yl)-2,4-dihydro-3H-1,2,4-triazole-3-thione (62i).** Nicotinohydrazide, **61i** (300 mg, 1.0 equiv, 2.19 mmol) was reacted with isothiocyanatomethane (176 mg, 1.1.0 equiv, 2.41 mmol) using general procedure C. Yield: 99% (418 mg) as a white solid. $^{1}$H NMR (500 MHz, DMSO-d6) δ 14.08 (s, 1H), 8.91 (d, J = 2.6 Hz, 1H), 8.76 (dd, J = 4.9, 1.7 Hz, 1H), 8.18 (dt, J = 7.9, 2.0 Hz, 1H), 7.60 (dd, J = 8.0, 4.7 Hz, 1H), 3.54 (s, 3H). $^{13}$C NMR (126 MHz, DMSO-d6) δ 168.2, 151.8, 149.8, 149.4, 136.7, 124.3, 123.1, 32.0. HRMS m/z $[M + H]^+$ Calcd for $[C_8H_8N_4S + H]^+$ 193.0548, found 193.0555.

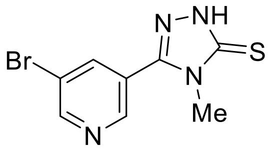

**5-(5-Bromopyridin-3-yl)-4-methyl-2,4-dihydro-3H-1,2,4-triazole-3-thione (62ii).** 5-Bromonicotinohydrazide, **61ii** (475 mg, 1.0 equiv, 2.20 mmol) was reacted with isothiocyanatomethane (161 mg, 150 µL, 1.0 equiv, 2.20 mmol) using general procedure C**.** Yield: 91% (542 mg) as a white solid: mp 255–275 °C. $^{1}$H NMR (400 MHz, DMSO-d6) δ 14.11 (s, 1H), 8.91 (dd, *J* = 9.9, 2.1 Hz, 2H), 8.46 (t, *J* = 2.1 Hz, 1H), 3.55 (s, 3H). HRMS (ESI): m/z [M - H]$^-$ Calcd for [$C_8H_7BrN_4$ - H]$^-$ 268.9496, found 268.9510.

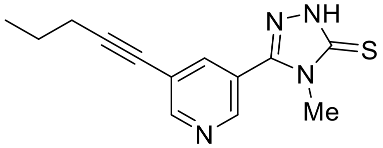

**4-Methyl-5-(5-(pent-1-yn-1-yl)pyridin-3-yl)-2,4-dihydro-3H-1,2,4-triazole-3-thione (62iii).** 5-(Pent-1-yn-1-yl)nicotinohydrazide, **61iii** (146 mg, 1.0 equiv, 718 µmol) was reacted with isothiocyanatomethane (53 mg, 1.0 equiv, 718 µmol) using general procedure C. Yield: 82% (153 mg) as an off-white solid: mp 146–148 °C. $^{1}$H NMR (400 MHz, $CDCl_3$) δ 12.99 (s, 1H), 8.93–8.70 (m, 2H), 7.95 (s, 1H), 3.69 (s, 3H), 2.42 (t, *J* = 7.0 Hz, 2H), 1.64 (h, *J* = 7.2 Hz, 2H), 1.04 (t, *J* = 7.4 Hz, 3H). $^{13}$C NMR (101 MHz, $CDCl_3$) δ 168.9, 153.9, 149.0, 146.5, 138.4, 122.1, 96.7, 76.3, 32.4, 21.9, 21.5, 13.7. HRMS (ESI): m/z [M + H]$^+$ Calcd for [$C_{13}H_{14}N_4S$ + H]$^+$ 259.1017, found 259.1005.

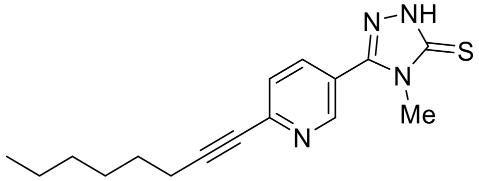

**4-Methyl-5-(6-(oct-1-yn-1-yl)pyridin-3-yl)-2,4-dihydro-3H-1,2,4-triazole-3-thione (62iv).** 6-(Oct-1-yn-1-yl)nicotinohydrazide, **61iv** (146 mg, 1.0 equiv, 595 µmol) was reacted with isothiocyanatomethane (44 mg, 1.0 equiv, 595 µmol) using general procedure C. Yield: 27% (48 mg) as a white solid; mp 141–144 °C. $^{1}$H NMR (400 MHz, $CDCl_3$) δ 12.78 (s, 1H), 8.91–8.86 (m, 1H), 7.91 (dd, *J* = 8.2, 2.3 Hz, 1H), 7.51 (dd, *J* = 8.2, 1.0 Hz, 1H), 3.69 (s, 3H), 2.46 (t, *J* = 7.2 Hz, 2H), 1.63 (p, *J* = 7.3 Hz, 2H), 1.50–1.37 (m, 2H), 1.36–1.22 (m, 4H), 0.87 (t, *J* = 6.9 Hz, 3H). $^{13}$C NMR (126 MHz, $CDCl_3$) δ 168.8, 149.3, 148.8, 146.1, 136.1, 126.9, 120.7, 95.2, 79.8, 32.4, 31.4, 28.7, 28.3, 22.6, 19.6, 14.1. HRMS (ESI): m/z [M + H] Calcd for [$C_{16}H_{20}N_4S$ + H]$^+$ 301.1487, found 301.1515.

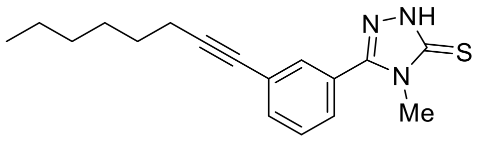

**4-Methyl-5-(3-(oct-1-yn-1-yl)phenyl)-2,4-dihydro-3H-1,2,4-triazole-3-thione (62v).** 3-(Oct-1-yn-1-yl)benzohydrazide, **61v** (100 mg, 1.0 equiv, 409 µmol) was reacted with isothiocyanatomethane (30 mg, 1.0 equiv, 409 μmol) using general procedure C**.** Yield: 86% (105 mg) as a light yellow solid; mp 79–80 °C. $^1$H NMR (400 MHz, $CDCl_3$) δ 12.39 (s, 1H), 7.64–7.38 (m, 4H), 3.65 (s, 3H), 2.40 (t, *J* = 7.1 Hz, 2H), 1.66 –1.54 (m, 2H), 1.50–1.38 (m, 2H), 1.37–1.25 (m, 5H), 0.89 (t, *J* = 6.9 Hz, 3H). $^{13}$C NMR (101 MHz, $CDCl_3$) δ 168.2, 151.7, 134.1, 131.5, 129.2, 127.5, 125.9, 125.6, 92.8, 79.4, 32.4, 31.4, 28.7, 28.6, 22.6, 19.5, 14.2. HRMS (ESI): m/z [M + H]$^+$ Calcd for $[C_{17}H_{21}N_3S + H]^+$ 300.1550, found 300.1534.

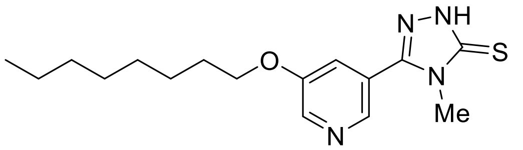

**4-Methyl-5-(5-(octyloxy)pyridin-3-yl)-2,4-dihydro-3H-1,2,4-triazole-3-thione (62vi).** 5-(Octyloxy)nicotinohydrazide, **61vi** (230 mg, 1.0 equiv, 867 μmol) was reacted with isothiocyanatomethane (63mg, 1.0 equiv, 867 μmol) using general procedure C. Yield: 92% (270 mg) as a white powder; mp 156–157 °C $^1$H NMR (400 MHz, $CDCl_3$) δ 13.1 (s, 1H), 8.5 (s, 2H), 7.5 (s, 1H), 4.1 (t, *J* = 6.5 Hz, 2H), 3.7 (s, 3H), 1.8 (q, *J* = 8.1, 7.3 Hz, 2H), 1.5–1.4 (m, 3H), 1.4–1.2 (m, 9H), 0.9 (t, *J* = 6.8 Hz, 3H). $^{13}$C NMR (101 MHz, $CDCl_3$) δ 168.9, 155.5, 149.6, 140.4, 140.3, 123.0, 120.9, 69.1, 32.5, 31.9, 29.4, 29.3, 29.1, 26.0, 22.7, 14.2. HRMS (ESI): m/z [M + H]$^+$ Calcd for $[C_{16}H_{24}N_4OS + H]^+$ 321.1749, found 321.1722.

**Synthesis of intermediate for compounds 64**
**Scheme S6.** Synthesis of **64**

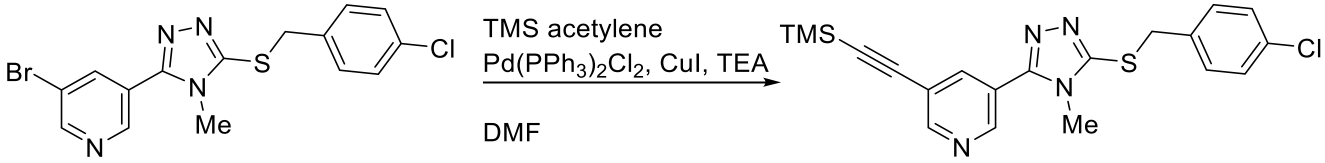

**3-(5-((4-Chlorobenzyl)thio)-4-methyl-4H-1,2,4-triazol-3-yl)-5-((trimethylsilyl)ethynyl)pyridine (63).** A mixture of 3-bromo-5-(5-((4-chlorobenzyl)thio)-4-methyl-4H-1,2,4-triazol-3-yl)pyridine, **38** (350 mg, 1.0 equiv, 885 μmol), bis(triphenylphosphine)palladium(II) chloride (31.0 mg, 0.05 equiv, 44.2 μmol) , and copper(I) iodide (17 mg, 0.1.0 equiv, 88.5 μmol) in DMF (5 mL) was purged with argon and treated with ethynyltrimethylsilane (174 mg, 248 μL, 2.0 equiv, 1.77 mmol) previously stirred with $Et_3N$ (179 mg, 0.25 mL, 2.0 equiv, 1.77 mmol) under argon. The resultant mixture was stirred at 50 °C for 18 h. Reaction mixture was diluted with saturated solution of $NaHCO_3$ and extracted with EtOAc (2x 15 mL), washed with water and brine, dried over $Na_2SO_3$ and purified by purified by MPLC (mobile phase: 100% EtOAc). Yield 53% (193 mg) as a brown solid. $^1$H NMR (500 MHz, $CDCl_3$) δ 8.80–8.74 (m, 2H), 7.99 (t, *J* = 2.1 Hz, 1H), 7.33–7.25 (m, 4H), 4.43 (s, 2H), 3.44 (s, 3H), 0.28 (s, 9H). $^{13}$C NMR (126 MHz, $CDCl_3$) δ 153.7, 152.9, 152.1, 147.6, 138.5, 135.4, 134.0, 130.6, 129.0, 123.1, 120.9, 100.4, 100.3, 37.5, 31.8. HRMS (ESI): m/z [M + H]$^+$ Calcd for $[C_{20}H_{21}N_4SSi + H]^+$ 413.0996, found 413.1023.
**Synthesis of intermediate 66**
**Scheme S7**. Synthesis of **66**

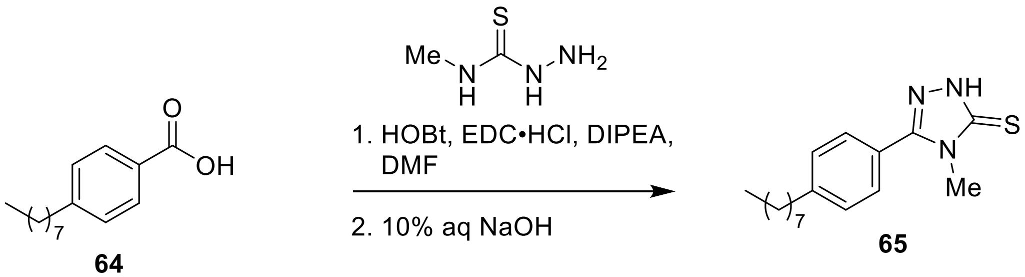

**4-Methyl-5-(4-octylphenyl)-4H-1,2,4-triazole-3-thione (64).** To a 50 mL round bottom flask equipped with a magnetic stir bar was added 4-Methyl-3-thiosemicarbazide (89.7 mg, 1.0 equiv, 853 µmol) , 4-octylbenzoic acid, **65** (200 mg, 1.0 equiv, 853 µmol) 1H-benzo[d][1,2,3]triazol-1-ol hydrate (157 mg, 1.2.0 equiv, 1.02 mmol), EDC hydrochloride (196 mg, 1.2.0 equiv, 1.02 mmol) and DIPEA (221 mg, 0.30 mL, 2.0 equiv, 1.71 mmol). The vessel was evacuated and refilled with argon three-time. DMF (10 mL) was added, and the reaction mixture stirred at room temperature under argon for two hours. On the consumption of 4-Methyl-3-thiosemicarbazide, the reaction mixture was diluted with 50 mL of EtOAc, washed sequentially with 50 mL 10% HCl, sat $Na_2CO_3$, water, and then brine. The organic phase was concentrated in-vivo to obtain the crude, white solid. The residue was refluxed in 10mL 10% NaOH was at 75 °C for 4 h. The reaction mixture was then cooled to room temperature, then 37% HCl was added gradually to reduce the pH to about 5. The precipitate formed was filtered, washed with water, and dried to obtain a white solid which was used for the next step without further purification. Yield 89% (231 mg) as a white solid: mp 102–104 °C. The compound $^1$H NMR (400 MHz, $CDCl_3$) δ 12.44 (s, 1H), 7.54–7.45 (m, 2H), 7.36–7.27 (m, 2H), 3.65 (s, 3H), 2.70–2.62 (m, 2H), 1.71–1.53 (m, 2H), 1.38–1.16 (m, 10H), 0.93–0.77 (m, 3H). $^{13}$C NMR (101 MHz, $CDCl_3$) δ 167.9, 152.4, 146.6, 129.3, 128.5, 123.0, 36.0, 32.4, 31.9, 31.3, 29.5, 29.3, 29.3, 22.7, 14.2. HRMS (ESI): m/z $[M+H]^+$ Calcd for [C17H25N3S + H]$^+$ 303.1848, found 303.1850.

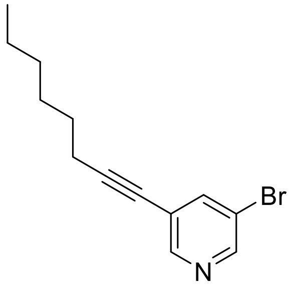

**3-Bromo-5-(oct-1-yn-1-yl)pyridine** (**67**) A mixture (A) of the 3,5-dibromopyridine (1000 mg, 1.0 equiv, 4.2 mmol) , copper(I) iodide (80 mg, 0.1.0 equiv, 0.42 mmol) , and bis(triphenylphosphine)palladium(II) chloride (0.15 g, 0.05 equiv, 0.21 mmol) in DMF (20 mL) was purged with argon for 10 min. Oct-1-yne (0.47 g, 0.62 mL, 1.0 Eq, 4.2 mmol), previously stirred with triethylamine (0.85 g, 1.2 mL, 2 Eq, 8.4 mmol) under argon, was added to mixture (A) dropwise over 10 mins. The resultant mixture was stirred at room temperature overnight. After completion of reaction, the mixture was diluted with $NaHCO_3$ and extracted with diethyl ether, washed with water and brine, dried over $Na_2SO_4$ and purified by MPLC (mobile phase: 0–2% EtOAc). Yield 78% (880 mg) as a light yellowish-green oil. $^1$H NMR (400 MHz, $CDCl_3$) δ 8.57–8.49 (m, 2H), 7.80 (t, $J$ = 2.0 Hz, 1H), 2.40 (t, $J$ = 7.2 Hz, 2H), 1.64–1.52 (m, 2H), 1.48–1.38 (m, 2H), 1.36–1.23 (m, 4H), 0.88 (t, $J$

= 6.9 Hz, 3H). $^{13}C$ NMR (101 MHz, $CDCl_3$) δ 150.5, 149.2, 140.9, 122.8, 120.1, 96.0, 76.1, 31.4, 28.7, 28.5, 22.6, 19.6, 14.2. Spectroscopic data agrees with reported compound.[4]

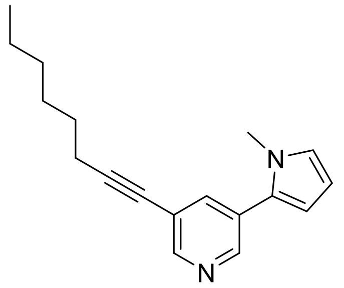

**3-(1-Methyl-1H-pyrrol-2-yl)-5-(oct-1-yn-1-yl)pyridine (91)** 1H-Pyrrole,1-methyl-2-(4,4,5,5-tetramethyl-1,3,2-dioxaborolan-2-yl) **67** (50 mg, 51 µL, 1.0 equiv, 0.24 mmol), 3-bromo-5-(oct-1-yn-1-yl)pyridine (77 mg, 1.2.0 equiv, 0.29 mmol), tetrakis (2.8 mg, 0.01.0 equiv, 2.4 µmol) $K_2CO_3$ (33 mg, 1 Eq, 0.24 mmol) were added to a 1 dram vial under argon. The vial was evacuated and flushed with argon. Water (2 mL) and dioxane (0.5 mL) were added to the reaction mixture after which the reaction mixture was stirred at 80 °C for 12 h. On completion of the reaction, the reaction mixture was diluted with water and extracted with EtOAC (2 x 10 mL). The organic layer was separated, dried ($Na_2SO_4$), and concentrated in vacuo. The residue was then purified by purified by MPLC *(*mobile phase: 0–20% EtOAc). Yield 33 % (21 mg) as a brown semi-solid. $^1H$ NMR (400 MHz, $CDCl_3$) δ 8.54 (dd, *J* = 5.5, 2.0 Hz, 2H), 7.68 (t, *J* = 2.1 Hz, 1H), 6.81 – 6.71 (m, 1H), 6.29 (dd, *J* = 3.7, 1.8 Hz, 1H), 6.22 (dd, *J* = 3.7, 2.7 Hz, 1H), 3.68 (s, 3H), 2.44 (t, *J* = 7.1 Hz, 2H), 1.69–1.57 (m, 2H), 1.52–1.40 (m, 2H), 1.39–1.29 (m, 4H), 0.91 (t, *J* = 6.8 Hz, 3H). $^{13}C$ NMR (101 MHz, $CDCl_3$) δ 150.2, 147.5, 137.8, 130.3, 128.8, 125.1, 120.9, 110.2, 108.4, 94.5, 77.4, 35.3, 31.5, 28.7, 28.6, 22.7, 19.6, 14.2. HRMS (ESI): m/z $[M + H]^+$ Calcd for $[C_{18}H_{22}N_2 + H]^+$ 267.1888, found 267.1861.

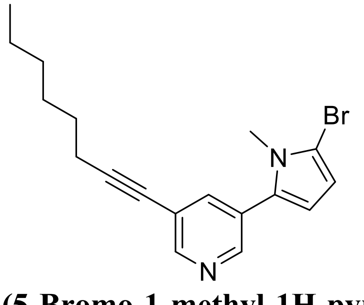

**3-(5-Bromo-1-methyl-1H-pyrrol-2-yl)-5-(oct-1-yn-1-yl)pyridine (68)** To 3-(1-methyl-1H-pyrrol-2-yl)-5-(oct-1-yn-1-yl)pyridine, **91** (20 mg, 1.0 equiv, 75 µmol) dissolved in $CH_2Cl_2$ (2 mL) was added NBS (8.6 mg, 2.0 equiv, 150 µmol). The reaction mixture was then stirred for 12 h at room temperature. The reaction mixture was diluted with water and extracted with $CH_2Cl_2$. The organic layer was separated, dried ($Na_2SO_4$), and concentrated in vacuo. The residue was then purified by purified by MPLC (mobile phase: 0-20% EtOAc). Yield 85% (22 mg) as a brown sticky solid. $^1H$ NMR (400 MHz, $CDCl_3$) δ 8.58–8.52 (m, 1H), 8.52–8.47 (m, 1H), 7.64 (t, *J* = 2.1 Hz, 1H), 6.29 – 6.24 (m, 2H), 3.59 (s, 3H), 2.43 (t, *J* = 7.1 Hz, 2H), 1.67–1.56 (m, 2H), 1.51–1.41 (m, 2H), 1.38–1.28 (m, 4H), 0.90 (t, *J* = 6.9 Hz, 3H). $^{13}C$ NMR (101 MHz, $CDCl_3$) δ 150.8, 147.5, 138.0, 131.5, 128.7, 121.1, 111.1, 110.6, 105.7, 94.8, 77.2, 34.0, 31.5, 28.7, 28.6, 22.7, 19.6, 14.2. HRMS (ESI): m/z $[M + H]^+$ Calcd for $[C_{18}H_{21}BrN_2 + H]^+$ 345.0955, found 345.0966.

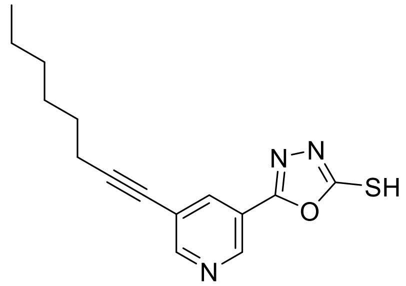

**5-(5-(Oct-1-yn-1-yl)pyridin-3-yl)-1,3,4-oxadiazole-2-thiol** (**69**) Carbon disulfide (217 mg, 172 µL, 1.0 equiv, 2.85 mmol) was added slowly to a solution of 5-(oct-1-yn-1-yl)nicotinohydrazide (700 mg, 1.0 equiv, 2.85 mmol) and KOH (160 mg, 1.0 equiv, 2.85 mmol) in 10 mL EtOH and water (4:1). The reaction mixture was refluxed for 12 h. Upon completion, the remaining ethanol was evaporated under reduced pressure. The residue was cooled to 0 °C and then acidified to pH 2 with 1N HCl. The was product was then filtered and washed with water to obtain a neutral pH compound.[5,6] Yield 80 % (652 mg) as off-white solid. mp 117–119 ºC $^1$H NMR (400 MHz, DMSO) δ 14.78 (br, 1H), 8.97 – 8.92 (m, 1H), 8.78 – 8.73 (m, 1H), 8.16 – 8.10 (m, 1H), 2.47 (t, *J* = 7.6, 7.1 Hz, 2H), 1.57 (p, *J* = 7.0 Hz, 2H), 1.48 – 1.36 (m, 2H), 1.35 – 1.21 (m, 4H), 0.91 – 0.84 (m, 3H). $^{13}$C NMR (101 MHz, DMSO) δ 177.6, 158.1, 154.0, 145.1, 135.3, 120.7, 119.1, 96.1, 76.2, 30.7, 27.9, 27.8, 22.0, 18.7, 13.9. HRMS (ESI): m/z [M + H]$^+$ Calcd for [$C_{15}H_{17}N_{23}OS$ + H]$^+$ 288.1171, found 288.1141

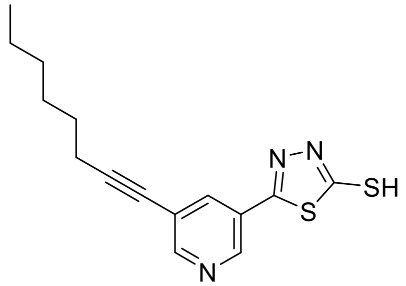

**5-(5-(Oct-1-yn-1-yl)pyridin-3-yl)-1,3,4-thiadiazole-2-thiol** (**70**) Carbon disulfide (93.1 mg, 73.9 µL, 2 Eq, 1.22 mmol) was added to a solution of the corresponding 5-(oct-1-yn-1-yl)nicotinohydrazide (150 mg, 1 Eq, 611 µmol) and KOH (68.6 mg, 2 Eq, 1.22 mmol) in ethanol (5 mL). The reaction mixture was stirred at room temperature for 12 h and then $Et_2O$ (5 mL) was added. The precipitate was filtered, washed with diethyl ether, and dried to give 194 mg potassium aroyl dithiocarbazate which was used for the next step without further purification or characterization. The precipitate was added in small portions to concentrated $H_2SO_4$ (1 mL) at 0 °C with vigorous stirring. The reaction mixture was then stirred in ice bath for 4 h and then poured into cold water (5 mL). The precipitate obtained was filtered and dissolved in an 10 % aqueous solution of sodium hydroxide and the insolubilities filtered. The solution was then acidified with HCl, and the final product was filtered and washed with water. Yield 47 % (88 mg) as a yellow solid. mp 157–158 ºC $^1$H NMR (400 MHz, $CDCl_3$) δ 12.36 (s, 1H), 8.88 (d, *J* = 2.2 Hz, 1H), 8.76 (d, *J* = 2.0 Hz, 1H), 7.96 (t, *J* = 2.1 Hz, 1H), 2.45 (t, *J* = 7.2 Hz, 2H), 1.63 (p, *J* = 7.2 Hz, 2H), 1.52–1.41 (m, 2H), 1.41–1.27 (m, 4H), 0.91 (t, *J* = 6.9 Hz, 3H). $^{13}$C NMR (101 MHz, $CDCl_3$) δ 188.8, 157.2, 154.0, 144.8, 136.5, 125.2, 122.4, 97.1, 76.1, 31.4, 28.8, 28.5, 22.7, 19.7, 14.2. HRMS (ESI): m/z [M + H]$^+$ Calcd for [$C_{15}H_{17}N_3S_2$ + H]$^+$ 304.0942, found 304.0930.

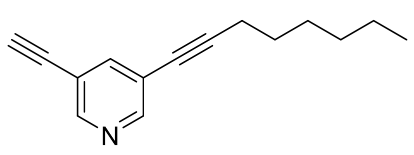

**3-Ethynyl-5-(oct-1-yn-1-yl)pyridine** (**71**). A mixture (A) 3-bromo-5-(oct-1-yn-1-yl)pyridine, **67** (350 mg, 494 µL, 1.0 equiv, 1.31 mmol), copper(I) iodide (25.0 mg, 0.1.0 equiv, 131 µmol) , and bis(triphenylphosphine)palladium(II) chloride (46.1 mg, 0.05 equiv, 65.7 µmol) DMF (5 mL) was purged with argon for 10 min. A solution of ethynyltrimethylsilane (129 mg, 1.0 equiv, 1.31 mmol) previously stirred with triethylamine (399 mg, 0.55 mL, 3 equivq, 3.94 mmol) under argon, was added to mixture (A) dropwise. The resultant mixture was stirred at room temperature for 12 h. The mixture then was diluted with $NaHCO_3$ and filtered. The filtrate was then extracted with diethyl ether, washed with water, dried over $Na_2SO_4$ and concentrated in vacuo. The residue was run through short column (20% EtOAc) and used for the next step without further characterization. Next, the residue dissolved in MeOH (5 mL) was added $K_2CO_3$ (763 mg, 5 equiv, 5.52 mmol) and stirred in room temperature for 12 h. On completion, the reaction mixture was evaporated in vacuo, and the residue dissolved in water and extracted with EtOAc (2 x 10 mL). The organic layer was separated, dried ($Na_2SO_4$), and concentrated in vacuo. The residue was then purified by purified by MPLC (mobile phase: 2–10% EtOAc). Yield 47 % (109 mg) as a brown semi-solid. $^{1}H$ NMR (400 MHz, $CDCl_3$) δ 8.55 (dd, $J$ = 5.1, 2.0 Hz, 2H), 7.73 (t, $J$ = 2.0 Hz, 1H), 3.19 (s, 1H), 2.40 (t, $J$ = 7.1 Hz, 2H), 1.64 – 1.54 (m, 2H), 1.40 (s, 2H), 1.38 – 1.26 (m, 4H), 0.89 (t, $J$ = 6.9 Hz, 3H). $^{13}C$ NMR (101 MHz 13C NMR (101 MHz, CDCl3) δ 151.7, 150.8, 141.4, 121.0, 118.8, 95.2, 81.0, 79.9, 76.6, 31.4, 28.7, 28.5, 22.7, 19.6, 14.2., $CDCl_3$) δ 151.7, 150.8, 141.4, 121.0, 118.8, 95.2, 81.0, 79.9, 76.6, 31.4, 28.7, 28.5, 22.7, 19.6, 14.2. HRMS (ESI): m/z $[M + H]^+$ Calcd for $[C_{15}H_{17}N + H]^+$ 212.1458, found 212.1439.

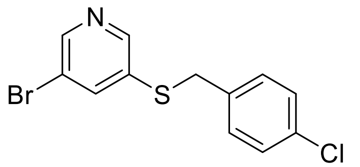

**3-Bromo-5-((4-chlorobenzyl)thio)pyridine (92)**. To (4-chlorophenyl)methanethiol (310 mg, 0.26 mL, 1.0 equiv, 1.95 mmol) dissolved in DMF (2 mL) and brought to 0 °C, was added NaH (93.8 mg, 60% Wt, 1.2.0 equiv, 2.34 mmol) and stirred for 2 h. 3,5-dibromopyridine (555 mg, 1.2 Eq, 2.34 mmol) was then added and the reaction mixture allowed to gradually warm up to room temperature. The reaction mixture was then subsequently heated at 80 °C overnight. On completion of the reaction, the reaction mixture was diluted with water and extracted with EtOAc (2 x 10 mL). The organic layer was separated, dried ($Na_2SO_4$), and concentrated in vacuo. The residue was then purified by purified by MPLC (mobile phase: 0–20% EtOAc). Yield 76 % (469 mg) as a brown solid. to give a light yellow oil.[7] $^{1}H$ NMR (400 MHz, $CDCl_3$) δ 8.48 (d, $J$ = 2.1 Hz, 1H), 8.39 (d, $J$ = 2.0 Hz, 1H), 7.69 (t, $J$ = 2.1 Hz, 1H), 7.30–7.16 (m, 5H), 4.07 (s, 2H). $^{13}C$ NMR (101 MHz, $CDCl_3$) δ 148.9, 148.9, 139.9, 134.9, 134.5, 133.8, 130.3, 129.1, 120.7, 38.4. HRMS (ESI): m/z $[M + H]^+$ Calcd for $[C_{12}H_9N + H]^+$ 313.9405, found 313.9406.

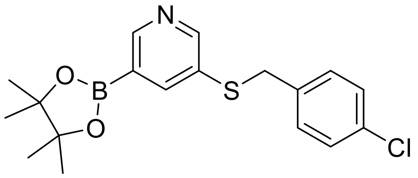

**3-((4-Chlorobenzyl)thio)-5-(4,4,5,5-tetramethyl-1,3,2-dioxaborolan-2-yl)**pyridine (**74**). 3-Bromo-5-((4-chlorobenzyl)thio)pyridine, **92** (150 mg, 1.0 equiv, 477 µmol), bis(pinacolato)diboron (133 mg, 1.1.0 equiv, 524 µmol), Pd(dppf)$Cl_2$ (34.9 mg, 0.1.0 equiv, 47.7 µmol) was added to a round bottom flask and flushed with argon for 20 min. Anhydrous toluene (2.5 mL) was then added. The reaction mixture then was heated to 100 °C and stirred for 18 hours. On completion, the reaction mixture was cooled to room temperature, and water (5 mL) was added. The mixture was then extracted with EtOAc (10 mL × 3). The combined organic layers was washed with water (10 mL) and brine (10 mL), dried over anhydrous $Na_2SO_4$, and the solvent was evaporated in vacuo to give brown solid product which was used without further purification.[7]

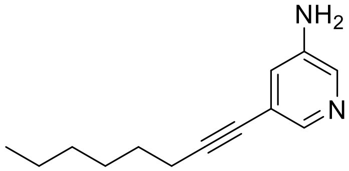

**5-(oct-1-yn-1-yl)pyridin-3-amine (76).** A mixture (A) of 5-bromopyridin-3-amine, **75** (1 g, 1 Eq, 6 mmol), bis(triphenylphosphine)palladium(II) chloride (0.2 g, 0.05 Eq, 0.3 mmol) , and copper(I) iodide (0.1 g, 0.1 Eq, 0.6 mmol) in DMF (15 mL)was purged with argon for 10 min. Oct-1-yne (0.8 g, 1 mL, 1.25 Eq, 7 mmol), previously stirred with triethylamine (2 mL, 2 Eq, 0.01 mol) under argon, was added to mixture (A) dropwise over 10 mins. The resultant mixture was stirred at room temperature overnight. After completion of reaction, the mixture was diluted with $NaHCO_3$ and extracted with EtOAc, washed with water and brine, dried over $Na_2SO_4$ and concentrated in vacuo. The residue was then purified by MPLC (mobile phase: 10–50% EtOAc). Yield 60% (727 mg) as a brown oil. $^1$H NMR (400 MHz, $CDCl_3$) δ 8.02 (s, 1H), 7.98–7.93 (m, 1H), 6.95 (dd, *J* = 2.7, 1.7 Hz, 1H), 3.70 (s, 2H), 2.38 (t, *J* = 7.1 Hz, 2H), 1.64–1.52 (m, 2H), 1.48–1.37 (m, 2H), 1.36–1.24 (m, 4H), 0.89 (t, *J* = 6.9 Hz, 3H).$^{13}$C NMR (101 MHz, $CDCl_3$) δ 142.8, 141.9, 136.1, 123.7, 121.1, 93.4, 77.7, 31.4, 28.7, 28.7, 22.7, 19.5, 14.2. HRMS (ESI): m/z $[M + H]^+$ Calcd for $[C_{13}H_{18}N_2 + H]^+$ 203.1548, found 203.1561.

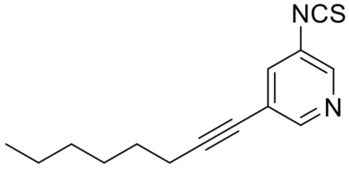

**3-isothiocyanato-5-(oct-1-yn-1-yl)pyridine (77).** To a solution of 5-(oct-1-yn-1-yl)pyridin-3-amine, **76** (205 mg, 1 Eq, 1.01 mmol) dissolved in 3 mL $CH_2Cl_2$ cooled to 0 °C was added NaOH (81.1 mg, 2 Eq, 2.03 mmol) and thiophosgene (175 mg, 117 µL, 1.5 Eq, 1.52 mmol) dropwise. The reaction mixture was stirred for an hour. On completion of the reaction, as determined by TLC, water (5 mL) was added. The mixture was then extracted with (10 mL × 3). The combined organic layer was washed with water (10 mL) and brine (10 mL), dried over anhydrous Na2SO4, and the solvent was concentrated in vacuo. The residue purified by MPLC (mobile phase: 0–10% EtOAc). Yield 78 % (193 mg) as a light dark brown liquid. $^1$H NMR (500 MHz, $CDCl_3$) δ 8.46 (d,

$J$ = 1.9 Hz, 1H), 8.37 (d, $J$ = 2.4 Hz, 1H), 7.47 (t, $J$ = 2.1 Hz, 1H), 2.41 (t, $J$ = 7.2 Hz, 2H), 1.60 (p, $J$ = 7.2 Hz, 2H), 1.48–1.39 (m, 2H), 1.36–1.27 (m, 4H), 0.90 (t, $J$ = 6.9 Hz, 3H). $^{13}$C NMR (126 MHz, $CDCl_3$) δ 150.2, 145.1, 140.0, 134.6, 129.2, 122.1, 96.3, 76.1, 31.4, 30.1, 28.7, 28.5, 22.7, 19.6, 14.2. HRMS (ESI): m/z $[M + H]^+$ Calcd for $[C_{14}H_{16}N_2S + H]^+$ 245.1113, found 245.1122.

**Synthesis of intermediates 93**

**Scheme S8**. Synthesis of 93

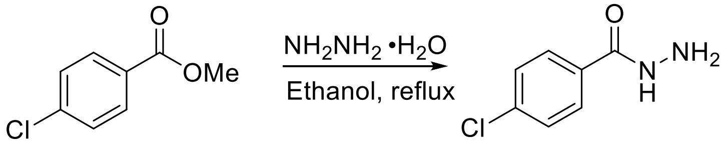

**92** **93**

**4-Chlorobenzohydrazide (93).** Methyl 4-chlorobenzoate, **92** (500 mg, 1.0 equiv, 2.93 mmol) was reacted with hydrazine hydrate (294 mg, 2.0 equiv, 5.86 mmol) using general procedure B. Yield: 98% (490 mg) as a white solid; 160–162 °C. (Lit[8] mp 162–163 °C). $^{1}$H NMR (400 MHz, $CD_3OD$) δ 7.81–7.73 (m, 2H), 7.51–7.43 (m, 2H). $^{13}$C NMR (101 MHz, $CD_3OD$) δ 168.5, 138.8, 133.0, 129.9, 129.8.

**Scheme S9.** Synthesis of 60

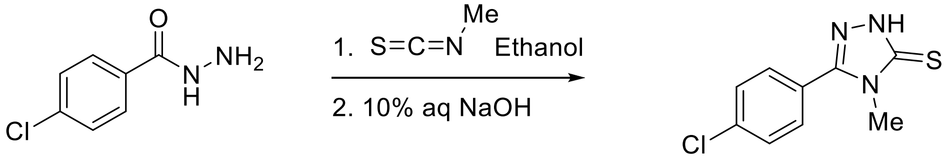

**94** **78**

**5-(4-Chlorophenyl)-4-methyl-2,4-dihydro-3H-1,2,4-triazole-3-thione (78).** 4-Chlorobenzohydrazide, **94** (324 mg, 1.0 equiv, 1.90 mmol) was reacted with isothiocyanatomethane (139 mg, 1.0 equiv, 1.90 mmol) using general procedure C**.** Yield: 89% (381 mg) as a white solid: mp 210–213 °C. (Lit[9] mp 210–212 °C) $^{1}$H NMR (400 MHz, DMSO) δ 13.98 (s, 1H), 7.83–7.70 (m, 2H), 7.69–7.57 (m, 2H), 3.52 (s, 3H). $^{13}$C NMR (101 MHz, DMSO) δ 167.6, 150.5, 135.6, 130.4, 129.1, 125.0, 31.6.

**Scheme S10.** Synthesis of 62

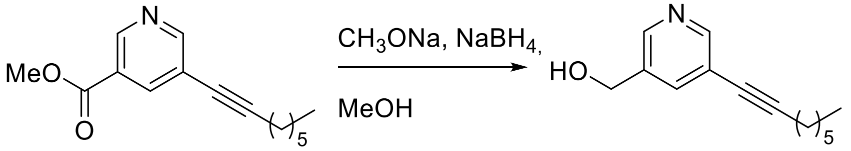

**95** **79**

**(5-(Oct-1-yn-1-yl)pyridin-3-yl)methanol (79).** To methyl 5-(oct-1-yn-1-yl)nicotinate, **95** (100 mg, 1.0 equiv, 408 µmol) dissolved in methanol (5 mL) was added sodium methoxide (1.10 mg, 0.05 equiv, 20.4 µmol) at room temperature and stirred for 5 min. $NaBH_4$ (30.8 mg, 2.0 equiv, 815 µmol) was added in one portion at 0 °C while constantly stirring. The reaction mixture was then brought to room temperature and stirred for 5 hours. The progress of the reaction was monitored by TLC. The reaction was quenched by adding excess methanol. The solution was then evaporated at reduced pressure, and the residue obtained was extracted with $CH_2Cl_2$ (3 × 10 mL). The combined extracts were dried over anhydrous sodium sulfate and the solvent was evaporated under reduced pressure. The residue was purified using MPLC (mobile phase:10% EtOAc/hexane) to obtain a yellow liquid. Yield: 58% (51 mg). $^1$H NMR (400 MHz, $CDCl_3$) δ 8.56 (d, *J* = 2.1 Hz, 1H), 8.47 (d, *J* = 2.2 Hz, 1H), 7.82 (t, *J* = 2.1 Hz, 1H), 4.78 (s, 2H), 2.52 (t, *J* = 7.2 Hz, 2H), 1.76–1.66 (m, 2H), 1.61–1.50 (m, 2H), 1.49–1.38 (m, 4H), 1.06–0.97 (m, 3H). $^{13}$C NMR (101 MHz, $CDCl_3$) δ 151.0, 146.3, 137.5, 136.3, 121.3, 94.6, 77.2, 62.1, 31.4, 28.7, 28.6, 22.6, 19.5, 14.1. HRMS (ESI): m/z $[M-H]^-$ Calcd for $[C14H19NO - H]^-$ 217.1467, found 217.1487

### 2.3 Synthetic Procedure for final analogs (in Tables 1–3)

The procedures and characterization data for the analogs listed in Table 4 are include in the main paper under methods section.

#### 2.3.1 General procedures for alkylation of 4,5-substituted-1,2,4-triazole-2-thione intermediates

*Method A1:* The appropriate alkylating agent (bromide or chloride) (1.2 equiv) and the corresponding 1,2,4-triazole-3-thione (1.0 equiv) were dissolved in 1:2 parts of MeOH/acetone (0.1 M) at room temperature. Upon addition of $K_2CO_3$ (1.5 equiv), the mixture was stirred overnight at room temperature. The completion of the reaction was monitored by TLC, after which the solvent was evaporated under reduced pressure. The residue was dissolved with EtOAc, washed with water (x2), brine. The extracted organic layer was dried over anhydrous $Na_2SO_4$, and then concentrated under low pressure. The resultant residue was then purified by MPLC.
*Method A2*[10]*:* The appropriate halide alkylating agent (1.2 equiv) was added to a solution of the corresponding 1,2,4-triazole-3-thiol (1.0 equiv) and $Et_3N$ (2.0 equiv) in $CH_3CN$ (0.1 M) and stirred for 4 h at room temperature. Upon completion of the reaction, the solvent was removed under reduced pressure. The residue was then dissolved in EtOAc and washed with water and brine. The extracted organic layer was dried over anhydrous $Na_2SO_4$, and concentrated under low pressure. The resultant residue was purified by MPLC.

**3-(4-Methyl-5-(methylthio)-4H-1,2,4-triazol-3-yl)-5-(oct-1-yn-1-yl)pyridine (2).** Methyl iodide (40 mg, 18 μL, 1.2 equiv, 0.28 mmol) was reacted with **55** (71 mg, 1.0 equiv, 0.24 mmol) according to general procedure A2. Final product was purified by MPLC (mobile phase: 5–14% MeOH/$CH_2Cl_2$). Yield 87% (65 mg) as a white solid; mp 61–63 ºC. $^1$H NMR (400 MHz, $CDCl_3$) δ 8.77–8.67 (m, 2H), 7.95 (t, *J* = 2.1 Hz, 1H), 3.60 (s, 3H), 2.78 (s, 3H), 2.43 (t, *J* = 7.1 Hz, 2H), 1.67–1.55 (m, 2H), 1.49–1.40 (m, 3H), 1.38–1.26 (m, 4H), 0.89 (t, *J* = 7.1 Hz, 3H). $^{13}$C NMR (101 MHz, $CDCl_3$) δ 154.0, 153.4, 152.9, 146.9, 138.3, 123.1, 121.8, 96.0, 76.6, 31.7, 31.4, 28.7, 28.5, 22.7, 19.6, 15.2, 14.2. HRMS (ESI): m/z $[M+H]^+$ Calcd for $[C_{17}H_{22}N_4S + H]^+$ 315.1643, found 315.1667.

**3-(5-(Benzylthio)-4-methyl-4H-1,2,4-triazol-3-yl)-5-(oct-1-yn-1-yl)pyridine (3).** Bromomethylbenzene (20 mg, 14 μL, 1.2 equiv, 0.12 mmol) was reacted with 4-methyl-5-(5-(oct-1-yn-1-yl)pyridin-3-yl)-4,5-dihydro-3H-1,2,4-triazole-3-thione (30 mg, 1.0 equiv, 99 μmol) according to general procedure A1. Final product was purified by flash MPLC (mobile phase: 40–80% EtOAc/Hex). Yield 46% (18 mg) as an off-white solid; mp 114–116 ºC. $^1$H NMR (400 MHz, $CDCl_3$) δ 8.70 (d, *J* = 2.0 Hz, 1H), 8.66 (d, *J* = 2.2 Hz, 1H),7.92 (t, *J* = 2.1 Hz, 1H), 7.29 (s, 4H), 4.41 (s, 2H), 3.31 (s, 3H), 2.44 (t, *J* = 7.1 Hz, 2H), 1.66–1.57 (m, 3H), 1.50–1.41 (m, 2H), 1.39–1.27 (m, 4H), 0.91 (t, *J* = 7.1 Hz, 3H). $^{13}$C NMR (101 MHz, $CDCl_3$) δ 153.5, 153.0, 152.1, 146.8,

138.3, 136.9, 129.2, 128.9, 128.1, 123.2, 121.8, 96.1, 76.6, 39.0, 31.7, 31.5, 28.7, 28.5, 22.7, 19.6, 14.2. HRMS (ESI): m/z $[M+H]^+$ Calcd for $[C_{23}H_{26}N_4S + H]^+$ 391.1956, found 391.1967.

**3-(5-((4-Bromobenzyl)thio)-4-methyl-4H-1,2,4-triazol-3-yl)-5-(oct-1-yn-1-yl)pyridine (4).** 1-Bromo-4-(bromomethyl)benzene (30 mg, 1.2 equiv, 0.12 mmol) was reacted with **55** (30 mg, 1.0 equiv, 0.10 mmol) according to general procedure A1. Final product was purified by flash MPLC (mobile phase: 40–80% EtOAc/Hex). Yield 81% (38 mg) as a yellowish solid; mp 87–89 ºC. $^1$H NMR (400 MHz, $CDCl_3$) δ 8.71 (d, *J* = 2.0 Hz, 1H), 8.69 (d, *J* = 2.2 Hz, 1H), 7.92 (t, *J* = 2.1 Hz, 1H), 7.46–7.39 (m, 2H), 7.24 (dd, *J* = 8.4, 6.4 Hz, 2H), 4.41 (s, 2H), 3.42 (s, 2H), 2.44 (t, *J* = 7.1 Hz, 2H), 1.66–1.56 (m, 2H), 1.51–1.39 (m, 2H), 1.38–1.26 (m, 4H), 0.90 (t, *J* = 7.0 Hz, 3H). $^{13}$C NMR (126 MHz, $CDCl_3$) δ 153.5, 153.0, 151.9, 146.8, 138.2, 136.0, 132.0, 130.9, 123.0, 122.1, 121.8, 96.1, 37.5, 31.8, 31.4, 28.7, 28.5, 22.7, 19.6, 14.2. HRMS (ESI): m/z $[M+H]^+$ Calcd for $[C_{23}H_{25}BrN_4S + H]^+$ 469.1062, found 469.1027.

**3-(5-((4-Fluorobenzyl)thio)-4-methyl-4H-1,2,4-triazol-3-yl)-5-(oct-1-yn-1-yl)pyridine (5)**. 1-(Bromomethyl)-4-fluorobenzene (23 mg, 15 µL, 1.2 equiv, 0.12 mmol) was reacted with **55** (30 mg, 1.0 equiv, 0.10 mmol). Final product was purified by MPLC (mobile phase: 50–80% EtOAc/Hex) according to general procedure A1. Yield 69% (28 mg) as white solid; mp 76–79 ºC. $^1$H NMR (400 MHz, $CDCl_3$) δ 8.71–8.62 (m, 1H), 7.91 (t, *J* = 2.1 Hz, 1H), 7.35–7.25 (m, 2H), 7.02–6.91 (m, 2H), 4.41 (s, 2H), 3.41 (s, 3H), 2.42 (t, *J* = 7.1 Hz, 2H), 1.66–1.54 (m, 2H), 1.49–1.39 (m, 2H), 1.43–1.24 (m, 4H), 0.89 (t, *J* = 6.9 Hz, 3H). $^{13}$C NMR (101 MHz, $CDCl_3$) δ 162.5 (d, *J* = 247.3 Hz), 153.4, 152.9, 152.1, 146.7, 138.2, 132.6 (d, *J* = 3.5 Hz), 130.9 (d, *J* = 8.2 Hz), 123.0, 121.7, 115.7 (d, *J* = 21.5 Hz), 96.1, 76.5, 37.5, 31.7, 31.4, 28.7, 28.5, 22.6, 19.6, 14.1. HRMS (ESI): m/z $[M+H]^+$ Calcd for $[C_{23}H_{25}FN_4S + H]^+$ 409.1862, found 409.1874.

**3-(4-Methyl-5-((4-methylbenzyl)thio)-4H-1,2,4-triazol-3-yl)-5-(oct-1-yn-1-yl)pyridine (6).** 1-(Bromomethyl)-4-methylbenzene (34 mg, 1.2 equiv, 0.18 mmol) was reacted with **55** (46 mg, 1.0 equiv, 0.15 mmol) according to general procedure A1. Final product was purified by MPLC (mobile phase: 20–60% EtOAc/Hex). Yield 60% (37 mg) as a light brown solid; mp 100–103 ºC. $^1$H NMR (400 MHz, $CDCl_3$) δ 8.77 (d, *J* = 2.0 Hz, 1H), 8.74 (d, *J* = 2.1 Hz, 1H), 7.97 (t, *J* = 2.1 Hz, 1H), 7.28–7.11 (m, 4H), 4.45 (s, 2H), 3.40 (s, 3H), 2.51 (t, *J* = 7.1 Hz, 2H), 2.40 (s, 3H), 1.76–1.64 (m, 2H), 1.58–1.48 (m, 2H), 1.46–1.35 (m, 5H), 0.98 (t, *J* = 7.0 Hz, 4H). $^{13}$C NMR (101 MHz, $CDCl_3$) δ 153.4, 152.9, 152.2, 146.8, 138.3, 137.9, 133.7, 129.5, 129.1, 123.2, 121.7, 96.0, 76.6, 38.7, 31.7, 31.4, 28.2, 28.1, 22.7, 21.3, 19.6, 14.2. HRMS (ESI): m/z $[M+H]^+$ Calcd for $[C_{24}H_{28}N_5S + H]^+$ 405.2113, found 405.2105.

**3-(5-((4-Methoxybenzyl)thio)-4-methyl-4H-1,2,4-triazol-3-yl)-5-(oct-1-yn-1-yl)pyridine (7).** 1-(Bromomethyl)-4-methoxybenzene (92.4 mg, 1.2 equiv, 459 µmol) was reacted with **55** (115 mg, 1.0 equiv, 383 µmol) according to general procedure A1. Final product was purified by MPLC (mobile phase: 20–60% EtOAc/Hex). Yield 67% (108 mg) as a white solid; mp 91–93 ºC. $^1$H NMR (400 MHz, $CDCl_3$) δ 8.70 (d, *J* = 2.0 Hz, 1H), 8.68 (d, *J* = 2.1 Hz, 1H), 7.91 (t, *J* = 2.1 Hz, 1H), 7.24–7.16 (m, 2H), 6.85–6.79 (m, 2H), 4.38 (s, 2H), 3.78 (s, 3H), 3.35 (s, 3H), 2.44 (t, *J* = 7.1 Hz, 2H), 1.62 (p, *J* = 7.4, 7.0 Hz, 2H), 1.51–1.38 (m, 2H), 1.39–1.28 (m, 4H), 0.90 (t, *J* = 7.0 Hz, 3H). $^{13}$C NMR (101 MHz, $CDCl_3$) δ 159.5, 153.4, 152.9, 152.3, 146.8, 138.3, 130.4, 128.7, 123.2, 121.8, 114.2, 96.1, 76.6, 55.4, 38.5, 31.8, 31.5, 28.7, 28.5, 22.7, 19.6, 14.2. HRMS (ESI): m/z $[M+H]^+$ Calcd for $[C_{24}H_{28}N_4OS + Cs]^+$ 553.1038, found 553.1006.

**3-(4-Methyl-5-((4-(trifluoromethyl)benzyl)thio)-4H-1,2,4-triazol-3-yl)-5-(oct-1-yn-1-yl)pyridine (10).** 1-(Bromomethyl)-4-(trifluoromethyl)benzene (41 mg, 1.2 equiv, 0.17 mmol) was reacted with **55** (43 mg, 1.0 equiv, 0.14 mmol) $K_2CO_3$ (40 mg, 2.0 equiv, 0.29 mmol) according to general procedure A1. Final product was purified by MPLC (mobile phase: 20–60% EtOAc/Hex). Yield 62% (41 mg) as a white solid; mp 118–120 ºC. $^1$H NMR (400 MHz, $CDCl_3$) δ 8.77–8.64 (m, 2H), 7.91 (t, *J* = 2.1 Hz, 1H), 7.64–7.41 (m, 4H), 4.52 (s, 2H), 3.45 (s, 3H), 2.44 (t, *J* = 7.1 Hz, 2H), 1.66–1.57 (m, 2H), 1.50–1.40 (m, 2H), 1.38–1.27 (m, 4H), 0.90 (t, *J* = 7.0 Hz, 3H). $^{13}$C NMR (126 MHz, $CDCl_3$) δ 153.5, 153.1, 151.8, 146.8, 141.1, 138.2, 130.2 (q, *J* = 32.6 Hz), 129.6, 125.8 (q, *J* = 3.5 Hz), 124.1 (q, *J* = 272.6 Hz), 122.93, 121.8, 96.2, 76.5, 37.2, 31.8, 31.4, 28.7, 28.5, 22.7, 19.6, 14.2. HRMS (ESI): m/z $[M-H]^-$ Calcd for $[C_{24}H_{25}FN_4S - H]^-$ 457.1674, found 457.1671.

**3-(5-(([1,1'-Biphenyl]-4-ylmethyl)thio)-4-methyl-4H-1,2,4-triazol-3-yl)-5-(oct-1-yn-1-yl)pyridine (11).** 4-(Bromomethyl)-1,1'-biphenyl (36 mg, 1.2 equiv, 0.14 mmol) was reacted with **55** (36 mg, 1.0 equiv, 0.12 mmol) (EtoAc 3 x 10 mL) according to general procedure A1. Final product was purified by was purified by MPLC (mobile phase: 20-60% EtOAc/Hex). Yield 60% (43 mg) as a white solid; mp 98–100 ºC. $^1$H NMR (400 MHz, CDCl3) δ 8.75–8.65 (m, 2H), 7.92 (t, *J* = 2.1 Hz, 1H), 7.60–7.51 (m, 4H), 7.48–7.31 (m, 5H), 4.49 (s, 2H), 3.40 (s, 3H), 2.43 (t, *J* = 7.1 Hz, 2H), 1.68–1.55 (m, 2H), 1.45 (p, *J* = 6.9 Hz, 2H), 1.39–1.28 (m, 4H), 0.91 (t, *J* = 7.0 Hz, 3H). $^{13}$C NMR (101 MHz, $CDCl_3$) δ 153.5, 153.0, 152.2, 146.8, 141.1, 140.5, 138.3, 135.8, 131.4, 129.7, 129.0, 127.7, 127.5, 127.2, 123.2, 121.8, 96.1, 76.8, 38.4, 31.79, 31.5, 28.7, 28.5, 22.7, 19.6, 14.2. HRMS (ESI): m/z $[M+H]^+$ Calcd for $[C_{29}H_{30}N_4S + H]^+$ 467.2269, found 467.2297.

**3-(5-((4-Chlorophenethyl)thio)-4-methyl-4H-1,2,4-triazol-3-yl)-5-(oct-1-yn-1-yl)pyridine (12).** 1-(2-Bromoethyl)-4-chlorobenzene (50 mg, 33 µL, 1.0 equiv, 0.23 mmol) was reacted with **55**(82 mg, 1.2 equiv, 0.27 mmol) according general procedure A2. Final product was purified by MPLC (mobile phase: 20-50 EtOAc/Hex). Yield 67% (67 mg) as a off-white solid; mp 75–77 ºC. $^1$H NMR (400 MHz, $CDCl_3$) δ 8.80–8.70 (m, 2H), 7.96 (t, *J* = 2.1 Hz, 1H), 7.32–7.15 (m, 4H), 3.60–3.50 (m, 5H), 3.12 (t, *J* = 7.4 Hz, 2H), 2.45 (t, *J* = 7.1 Hz, 2H), 1.68–1.57 (m, 2H), 1.52–1.40 (m, 2H), 1.40–1.28 (m, 4H), 0.91 (t, *J* = 7.1 Hz, 3H). $^{13}$C NMR (101 MHz, $CDCl_3$) δ 153.4, 152.8, 146.8, 138.2, 137.9, 132.6, 130.2, 128.7, 123.0, 121.7, 96.0, 76.6, 35.3, 34.2, 31.8, 31.4, 28.7, 28.5, 22.6, 19.6, 14.2. HRMS (ESI): m/z $[M+H]^+$ Calcd for $[C_{24}H_{27}ClN_4S + H]^+$ 439.1723, found 439.1715.

**3-(5-((4-Chlorophenyl)thio)-4-methyl-4H-1,2,4-triazol-3-yl)-5-(oct-1-yn-1-yl)pyridine (13).** To a 1 dram vial was added 1-bromo-4-chlorobenzene (35 mg, 1.0 equiv, 0.18 mmol), DIPEA (47 mg, 64 µL, 2.0 equiv, 0.37 mmol), dry 1,4-Dioxane (1 mL). The mixture was then evacuated and backfilled with argon three times. $Pd_2(dba)_3$ (8.4 mg, 0.05 equiv, 9.1 µmol), xantphos (11 mg, 0.1 equiv, 18 µmol) and **55** (55 mg, 1.0 equiv, 0.18 mmol) were added to the reaction mixture. The mixture was then degassed twice and refluxed overnight. On completion of reaction, as determined by TLC, the reaction mixture was allowed to reach ambient temperature, filtered and concentrated under reduced pressure. The crude mixture was purified by MPLC (mobile phase: 10–20 EtOAc/Hex) to obtain final product. Yield 33% (25 mg) as a white solid; mp 53–55 ºC. $^1$H NMR (400 MHz, $CDCl_3$) δ 8.75 (d, *J* = 2.2 Hz, 1H), 8.72 (d, *J* = 2.0 Hz, 1H), 7.98 (t, *J* = 2.1 Hz, 1H), 7.41–7.29 (m, 4H), 3.66 (s, 3H), 2.43 (t, *J* = 7.1 Hz, 2H), 1.67–1.55 (m, 2H), 1.49–1.40 (m, 2H),

1.37–1.27 (m, 4H), 0.90 (t, *J* = 7.0 Hz, 3H). $^{13}$C NMR (101 MHz, $CDCl_3$) δ 153.9, 153.7, 150.0, 146.8, 138.3, 134.7, 131.8, 130.0, 129.5, 122.9, 121.8, 96.3, 76.50, 32.5, 31.4, 28.7, 28.5, 22.7, 19.6, 14.2. HRMS (ESI): m/z [M+H]$^+$ Calcd for [$C_{22}H_{23}N_4S$ + H]$^+$ 411.1410, found 411.1398.

**3-(5-((3-Chlorobenzyl)thio)-4-methyl-4H-1,2,4-triazol-3-yl)-5-(oct-1-yn-1-yl)pyridine (14).** 1-(Bromomethyl)-3-chlorobenzene (58 mg, 37 µL, 1.2 equiv, 0.28 mmol) was reacted with **55** (71 mg, 1.0 equiv, 0.24 mmol) according to general procedure A1. Final product was purified by MPLC (mobile phase: 20–80% EtOAc/Hex). Yield 75% (75 mg) as yellow solid; mp 78–80 ºC. $^1$H NMR (400 MHz, $CDCl_3$) δ 8.73–8.67 (m, 2H), 7.93 (t, *J* = 2.1 Hz, 1H), 7.33–7.19 (m, 4H), 4.40 (s, 2H), 3.41 (s, 3H), 2.44 (t, *J* = 7.1 Hz, 2H), 1.68–1.58 (m, 2H), 1.52–1.40 (m, 2H), 1.39–1.26 (m, 4H), 0.91 (t, *J* = 6.7 Hz, 3H). $^{13}$C NMR (101 MHz, $CDCl_3$) δ 153.2, 153.0, 151.9, 146.4, 138.9, 138.5, 134.6, 130.1, 129.2, 128.2, 127.4, 123.1, 122.0, 96.4, 76.5, 37.9, 31.8, 31.4, 28.7, 28.5, 22.7, 19.6, 14.2. HRMS (ESI): m/z [M+H]$^+$ Calcd for [$C_{23}H_{25}ClN_4S$ + H]$^+$ 425.1567, found 425.1565.

**3-(4-Methyl-5-((3-nitrobenzyl)thio)-4H-1,2,4-triazol-3-yl)-5-(oct-1-yn-1-yl)pyridine (15).** 1-(Bromomethyl)-3-nitrobenzene (52 mg, 1.2 equiv, 0.24 mmol) was reacted with **55** (60 mg, 1.0 equiv, 0.20 mmol) according to general procedure A1. Final product was purified by MPLC (mobile phase: 100% EtOAc). Yield 54% (47 mg) as light brown solid; mp 82–84 ºC. $^1$H NMR (400 MHz, $CDCl_3$) δ 8.74–8.67 (m, 1H), 8.28 (t, *J* = 2.0 Hz, 1H), 8.16–8.08 (m, 1H), 7.95 (t, *J* = 2.0 Hz, 1H), 7.79 (d, *J* = 7.7 Hz, 1H), 7.48 (t, *J* = 8.0 Hz, 1H), 4.59 (s, 1H), 3.53 (s, 2H), 2.43 (t, *J* = 7.1 Hz, 1H), 1.67–1.54 (m, 1H), 1.49–1.38 (m, 1H), 1.37–1.26 (m, 2H), 0.89 (t, *J* = 7.0 Hz, 2H). $^{13}$C NMR (101 MHz, $CD_3OD$) δ 154.6, 154.1, 153.1, 149.7, 147.9, 140.9, 139.7, 136.4, 131.0, 124.8, 124.4, 123.7, 123.2, 97.1, 77.2, 37.9, 32.5, 32.5, 29.7, 29.5, 23.6, 20.1, 14.4. HRMS (ESI): m/z [M+H]$^+$ Calcd for [$C_{23}H_{25}N_5O_2S$ + H]$^+$ 436.1807, found 436.1805.

**3-(5-((3,4-Dichlorobenzyl)thio)-4-methyl-4H-1,2,4-triazol-3-yl)-5-(oct-1-yn-1-yl)pyridine (16)**. 4-(Bromomethyl)-1,2-dichlorobenzene (38 mg, 23 µL, 1.2 equiv, 0.16 mmol) was reacted with **55** (40 mg, 1.0 equiv, 0.13 mmol according to general procedure A1. Final product was purified by MPLC (mobile phase: 20–60% EtOAc/Hex). Yield 65% (40 mg) as white solid; mp 85–88 ºC. $^1$H NMR (500 MHz, $CDCl_3$) δ 8.74–8.69 (m, 2H), 7.93 (t, *J* = 2.1 Hz, 1H), 7.48 (d, *J* = 2.2 Hz, 1H), 7.40–7.35 (m, 1H), 7.28–7.22 (m, 2H), 4.43 (s, 2H), 3.48 (s, 3H), 2.44 (t, *J* = 7.1 Hz, 2H), 1.67 – 1.57 (m, 2H), 1.50 – 1.41 (m, 2H), 1.38 – 1.28 (m, 4H), 0.91 (t, *J* = 7.0 Hz, 3H). $^{13}$C NMR (126 MHz, $CDCl_3$) δ 153.6, 153.2, 151.6, 146.8, 138.3, 137.2, 132.9, 132.2, 131.1, 130.8, 128.7, 122.9, 121.8, 96.2, 76.9, 36.7, 31.8, 31.5, 28.7, 28.5, 22.7, 19.6, 14.2. HRMS (ESI): m/z [M+H]$^+$ Calcd for [$C_{23}H_{24}Cl_2N_4S$ + H]$^+$ 459.1177, found 459.1143.

**3-(5-((3,5-Difluorobenzyl)thio)-4-methyl-4H-1,2,4-triazol-3-yl)-5-(oct-1-yn-1-yl)pyridine (17).** 1-(Bromomethyl)-3,5-difluorobenzene (0.10 g, 65 µL, 1.2 equiv, 0.50 mmol) was reacted with **55** (80 mg, 1.0 equiv, 0.27 mmol) according to general procedure A1. Final product was purified by MPLC (mobile phase: 40–100% EtOAc/Hex). Yield 91 % (103 mg) as off-white solid; mp 69–71 ºC. $^1$H NMR (400 MHz, $CDCl_3$) δ 8.71 (s, 2H), 7.97 (t, *J* = 2.0 Hz, 1H), 6.96–6.88 (m, 2H), 6.8–6.7 (m, 1H), 4.45 (s, 2H), 3.50 (s, 3H), 2.44 (t, *J* = 7.1 Hz, 2H), 1.61 (p, *J* = 7.2 Hz, 2H), 1.50–1.39 (m, 2H), 1.37–1.27 (m, 4H), 0.90 (t, *J* = 7.1 Hz, 3H). $^{13}$C NMR (101 MHz, $CDCl_3$) δ 163.1 (dd, *J* = 249.5, 12.7 Hz), 153.1, 153.0, 151.8, 146.3, 140.6 (t, *J* = 9.5 Hz), 138.6, 123.1,

122.0, 113.94–110.4 (m), 103.6 (t, $J$ = 25.2 Hz), 96.5, 76.4, 37.0, 31.8, 31.4, 28.7, 28.5, 22.7, 19.6, 14.2. HRMS (ESI): m/z $[M-H]^-$ Calcd for $[C_{23}H_{24}F_2N_4S - H]^-$ 425.1612, found 425.1631.

**3-(5-((3,5-Bis(trifluoromethyl)benzyl)thio)-4-methyl-4H-1,2,4-triazol-3-yl)-5-(oct-1-yn-1-yl)pyridine (19).** 1-(Bromomethyl)-3,5-bis(trifluoromethyl)benzene (58.0 mg, 34.7 µL, 1.2 equiv, 189 µmol) was reacted with **55** (47.3 mg, 1.0 equiv, 157 µmol) according to general procedure A1. Final product was purified by flash chromatography (mobile phase: 40–80% EtOAc/Hex). Yield 42% (35 mg) as a white solid; mp 101–103 ºC. $^1$H NMR (400 MHz, CDCl3) δ 8.71 (s, 2H), 7.95–7.87 (m, 3H), 7.79 (s, 1H), 4.61 (s, 2H), 3.50 (s, 3H), 2.43 (t, $J$ = 7.1 Hz, 2H), 1.66–1.54 (m, 2H), 1.50–1.38 (m, 2H), 1.37–1.24 (m, 4H), 0.90 (t, $J$ = 7.0 Hz, 3H). $^{13}$C NMR (126 MHz, $CDCl_3$) δ 153.6, 153.3, 151.3, 146.8, 139.8, 138.2, 132.1 (q, $J$ = 33.4 Hz), 129.4 (q, $J$ = 4.3 Hz), 123.2 (q, $J$ = 273.1 Hz), 122.8, 122.1–121.8 (m), 96.2, 76.5, 36.4, 31.7, 31.4, 28.7, 28.5, 22.7, 19.6, 14.2. HRMS (ESI): m/z $[M+H]^+$ Calcd for $[C_{25}H_{24}F_6N_4S + H]^+$ 526.1609, found 526.1626.

**3-(4-Methyl-5-((2-(2-methyl-5-nitro-1H-imidazol-1-yl)ethyl)thio)-4H-1,2,4-triazol-3-yl)-5-(oct-1-yn-1-yl)pyridine (22).** 1-(2-Bromoethyl)-2-methyl-5-nitro-1H-imidazole (49 mg, 1.1 equiv, 0.21 mmol) was reacted with **55**(57 mg, 1.0 equiv, 0.19 mmol) according to general procedure A1. Final product was purified by MPLC (mobile phase: 2–10% MeOH/ $CH_2Cl_2$). Yield 59% (51 mg) as a yellow solid; mp 115–117 ºC. $^1$H NMR (400 MHz, CDCl3) δ 8.82–8.66 (m, 2H), 7.96 (d, $J$ = 2.2 Hz, 2H), 4.88–4.81 (m, 2H), 3.68–3.59 (m, 5H), 2.62 (s, 3H), 2.44 (t, $J$ = 7.1 Hz, 2H), 1.67–1.56 (m, 2H), 1.51–1.39 (m, 2H), 1.38–1.27 (m, 4H), 0.89 (t, $J$ = 6.9 Hz, 3H). $^{13}$C NMR (101 MHz, $CDCl_3$) δ 153.7, 153.3, 151.9, 151.7, 146.8, 138.5, 138.1, 133.6, 122.7, 121.9, 96.3, 76.5, 45.5, 31.8, 31.7, 31.4, 28.7, 28.5, 22.7, 19.6, 14.7, 14.2. HRMS (ESI): m/z $[M+H]^+$ Calcd for $[C_{22}H_{27}N_7O_2S + Cs]^+$ 586.1002, found 586.0984.

**3-(4-Methyl-5-(((1-methyl-5-nitro-1H-imidazol-2-yl)methyl)thio)-4H-1,2,4-triazol-3-yl)-5-(oct-1-yn-1-yl)pyridine (23).** To (1-methyl-5-nitro-1H-imidazo-2-yl)-methanol (31 mg, 1.2 equiv, 0.20 mmol) dissolved in anhydrous THF (2 mL) was added triphenylphosphine (1.2 equiv) and **55**(50 mg, 1.0 equiv, 0.17 mmol) at 0 °C under argon. After stirring for 30 min, DIAD (34 mg, 32 µL, 1.0 equiv, 0.17 mmol) was added dropwise. The mixture was then warmed to room temperature and stirred for 3 h. Solvent was removed in vacuo, and the residue was purified by MPLC (0–5% MeOH/$CH_2Cl_2$) to afford the final product.[11] Yield 62% (45 mg) as a white solid; mp 112–113 ºC. $^1$H NMR (400 MHz, $CD_3OD$) δ 8.76 (d, $J$ = 2.2 Hz, 1H), 8.69 (d, $J$ = 2.0 Hz, 1H), 8.12 (t, $J$ = 2.1 Hz, 1H), 7.83 (s, 1H), 4.50 (s, 2H), 4.06 (s, 3H), 3.76 (s, 3H), 2.48 (t, $J$ = 7.0 Hz, 2H), 1.69–1.57 (m, 2H), 1.55–1.43 (m, 2H), 1.41–1.28 (m, 4H), 0.92 (t, $J$ = 7.0 Hz, 3H). $^{13}$C NMR (101 MHz, $CD_3OD$) δ 155.0, 154.1, 151.4, 149.3, 148.0, 141.1, 139.7, 131.9, 124.4, 123.1, 97.1, 77.2, 34.4, 32.8, 32.5, 30.7, 29.7, 29.5, 23.6, 20.1, 14.4. HRMS (ESI): m/z $[M–H]^-$ Calcd for $[C_{21}H_{25}N_7O_2S – H]^-$ 438.1712, 438.1734.

**3-(5-((4-Fluoro-3-nitrobenzyl)thio)-4-methyl-4H-1,2,4-triazol-3-yl)-5-(oct-1-yn-1-yl)pyridine (25).** 4-(bromomethyl)-1-fluoro-2-nitrobenzene was reacted with **55** (49.5 mg, 1.0 equiv, 165 µmol) according to general procedure A1. Final product was purified by flash chromatography (40–80% EtOAc/Hex). Yield 35% (26 mg) as a white solid; mp 77–78 ºC. $^1$H NMR (400 MHz, $CDCl_3$) δ 8.71 (s, 2H), 8.14 (dd, $J$ = 7.0, 2.4 Hz, 1H), 7.93 (t, $J$ = 2.0 Hz, 1H), 7.82–7.75 (m, 1H), 7.22 (dd, $J$ = 10.5, 8.6 Hz, 1H), 4.54 (s, 2H), 3.55 (s, 3H), 2.43 (t, $J$ = 7.1 Hz,

2H), 1.66–1.55 (m, 2H), 1.50–1.37 (m, 2H), 1.37–1.24 (m, 4H), 0.89 (t, *J* = 7.0 Hz, 3H). $^{13}$C NMR (126 MHz, $CDCl_3$) δ 155.0 (d, *J* = 265.7 Hz), 153.6, 153.2, 151.5, 146.8, 138.2, 137.4 (d, *J* = 7.9 Hz), 136.6 (d, *J* = 8.6 Hz), 134.4 (d, *J* = 4.1 Hz), 126.6 (d, *J* = 2.3 Hz), 122.8, 121.8, 118.7 (d, *J* = 20.9 Hz), 96.2, 76.5, 35.4, 31.8, 31.4, 28.7, 28.5, 22.7, 19.6, 14.2. HRMS (ESI): m/z $[M+H]^+$ Calcd for $[C_{23}H_{24}FN_5O_2S + H]^+$ 454.1713, found 454.1704.

**3-(5-((4-Chloro-2-nitrobenzyl)thio)-4-methyl-4H-1,2,4-triazol-3-yl)-5-(oct-1-yn-1-yl)pyridine (26).** 4-Chloro-1-(chloromethyl)-2-nitrobenzene (33 mg, 1.2 equiv, 0.16 mmol) was reacted with **55** (40 mg, 1.0 equiv, 0.13 mmol) according to general procedure A1. Final product was purified by flash chromatography (mobile phase: 60–100 EtOAc/Hex). Yield 66% (41 mg) as a yellow solid; mp 76–78 ºC, $^{1}$H NMR (400 MHz, $CDCl_3$) δ 8.76–8.66 (m, 2H), 8.09 (d, *J* = 2.2 Hz, 1H), 7.92 (t, *J* = 2.1 Hz, 1H), 7.83 (d, *J* = 8.3 Hz, 1H), 7.53 (dd, *J* = 8.3, 2.2 Hz, 1H), 4.84 (s, 2H), 3.52 (s, 3H), 2.43 (s, 2H), 1.67–1.54 (m, 2H), 1.51–1.39 (m, 2H), 1.38–1.26 (m, 4H), 0.90 (t, *J* = 6.9 Hz, 3H). $^{13}$C NMR (101 MHz, $CDCl_3$) δ 153.5, 153.2, 152.1, 148.1, 146.8, 138.1, 135.0, 134.6, 134.0, 131.9, 125.7, 122.9, 121.8, 96.2, 76.6, 33.8, 31.8, 31.4, 28.7, 28.5, 22.7, 19.6, 14.2. HRMS (ESI): m/z $[M+H]^+$ Calcd for $[C_{23}H_{24}ClN_5O_2S + H]^+$ 470.1418, found 470.1413.

**3-(4-Methyl-5-((3-nitro-5-(trifluoromethyl)benzyl)thio)-4H-1,2,4-triazol-3-yl)-5-(oct-1-yn-1-yl)pyridine (27).** 1-(Chloromethyl)-3-nitro-5-(trifluoromethyl)benzene (37 mg, 1.2 equiv, 0.17 mmol) was reacted with **55** (42 mg, 1.0 equiv, 0.14 mmol) according to general procedure A1. Final product was purified by MPLC (mobile phase:40–80% EtOAc/Hex). Yield 68% (48 mg) as an off-white solid; mp 96–98 ºC. $^{1}$H NMR (400 MHz, $CDCl_3$) δ 8.71 (s, 2H), 8.55 (s, 1H), 8.39 (s, 1H), 8.09 (s, 1H), 7.93 (s, 1H), 4.67 (s, 2H), 3.57 (s, 3H), 2.43 (t, *J* = 7.1 Hz, 2H), 1.67–1.55 (m, 2H), 1.49–1.39 (m, 2H), 1.37–1.26 (m, 4H), 0.90 (t, *J* = 7.1 Hz, 3H). $^{13}$C NMR (126 MHz, DMSO-d6) δ 152.8, 152.5, 150.3, 148.0, 147.2, 142.3, 137.5, 132.0 (q, *J* = 3.7 Hz), 130.2 (q, *J* = 33.5 Hz), 127.7, 124.0, 123.1, 122.9 (q, *J* = 272.1 Hz), 120.3, 119.4 (q, *J* = 4.0 Hz), 95.5, 76.7, 35.1, 31.7, 30.7, 28.0, 27.9, 22.0, 18.7, 13.9.HRMS (ESI): m/z $[M+H]^+$ Calcd for $[C_{24}H_{24}F_3N_5O_2S + H]^+$ 504.1681, found 504.1670.

**4-((4-Methyl-5-(5-(oct-1-yn-1-yl)pyridin-3-yl)-4H-1,2,4-triazol-3-yl)thio)-7-nitrobenzo[c][1,2,5]oxadiazole (28).** NBD-chloride (41 mg, 37 µL, 1 equiv, 0.20 mmol) was reacted with **55** (61 mg, 1.0 equiv, 0.20 mmol) according to general procedure A1. Final product was purified by MPLC (50–80% EtOAc/Hex). Yield 55% (52 mg) as a yellow solid; mp 139–141 ºC. $^{1}$H NMR (400 MHz, $CDCl_3$) δ 8.82 (d, *J* = 2.3 Hz, 1H), 8.76 (d, *J* = 2.1 Hz, 1H), 8.39 (d, *J* = 7.7 Hz, 1H), 8.04 (t, *J* = 2.1 Hz, 1H), 7.54 (d, *J* = 7.7 Hz, 1H), 3.90 (s, 3H), 2.45 (t, *J* = 7.1 Hz, 2H), 1.68–1.56 (m, 2H), 1.52–1.39 (m, 2H), 1.37–1.26 (m, 3H), 0.89 (t, *J* = 7.0 Hz, 2H). $^{13}$C NMR (101 MHz, $CDCl_3$) δ 154.7, 154.1, 149.0, 146.8, 145.0, 142.7, 138.4, 135.4, 133.3, 130.4, 126.9, 122.4, 122.0, 96.7, 76.3, 33.0, 31.4, 28.7, 28.5, 22.7, 19.6, 14.2. HRMS (ESI): m/z $[M+H]^+$ Calcd for $[C_{22}H_{21}N_7O_3S + H]^+$ 464.1505, found 464.1471.

**5-(((4-Methyl-5-(5-(oct-1-yn-1-yl)pyridin-3-yl)-4H-1,2,4-triazol-3-yl)thio)methyl)benzo[c][1,2,5]oxadiazole (29).** 5-(Bromomethyl)benzo[c][1,2,5]oxadiazole (45 mg, 1 equiv, 0.21 mmol) was reacted with **55**(64 mg, 1.0 equiv, 0.21 mmol) according to general procedure A2. Final product was purified by MPLC (mobile phase:40-70 EtOAc/Hex). Yield 87% (80 mg) as a white solid; mp 106–107ºC. $^{1}$H NMR (400 MHz, $CDCl_3$) δ 8.72–8.66 (m, 2H), 7.91 (t, *J* = 2.1 Hz, 1H), 7.84 (s, 1H), 7.80 (d, *J* = 9.3, 1.0 Hz, 1H), 7.53 (dd, *J* = 9.3, 1.4 Hz, 1H), 4.59

(d, $J$ = 0.9 Hz, 2H), 3.54 (s, 3H), 2.42 (t, $J$ = 7.1 Hz, 2H), 1.66–1.54 (m, 2H), 1.48–1.38 (m, 2H), 1.35–1.26 (m, 4H), 0.89 (t, $J$ = 7.0 Hz, 3H). $^{13}$C NMR (101 MHz, $CDCl_3$) δ 153.5, 153.3, 151.4, 149.2, 148.6, 146.7, 140.8, 138.2, 133.5, 122.8, 121.8, 117.1, 115.7, 96.2, 76.5, 37.1, 31.81, 31.4, 28.7, 28.5, 22.6, 19.6, 14.2. HRMS (ESI): m/z [M+H]$^+$ Calcd for [$C_{23}H_{24}N_6OS$ + H]$^+$ 433.1811, found 433.1825.

**3-(5-((4-Chlorobenzyl)thio)-4-ethyl-4H-1,2,4-triazol-3-yl)-5-(oct-1-yn-1-yl)pyridine (30).** 1-(Bromomethyl)-4-chlorobenzene (47 mg, 1.2 equiv, 0.23 mmol) was reacted with 4-Ethyl-5-(5-(oct-1-yn-1-yl)pyridin-3-yl)-2,4-dihydro-3H-1,2,4-triazole-3-thione, **56** (60 mg, 1.0 equiv, 0.19 mmol) according to general procedure A1. Final product was purified by MPLC (mobile phase: 20–50% EtOAc/Hex). Yield 61% (51 mg) as a yellowish white solid; mp 89–91 ºC. $^1$H NMR (400 MHz, $CDCl_3$) δ 8.85–8.55 (m, 2H), 7.96 (s, 1H), 7.35–7.25 (m, 4H), 4.50 (s, 2H), 3.87 (q, $J$ = 7.2 Hz, 2H), 2.44 (t, $J$ = 7.1 Hz, 2H), 1.62 (p, $J$ = 7.2 Hz, 2H), 1.49–1.40 (m, 2H), 1.36–1.28 (m, 4H), 1.23 (t, $J$ = 7.2 Hz, 4H), 0.90 (t, $J$ = 6.9 Hz, 3H). $^{13}$C NMR (126 MHz, $CDCl_3$) δ 153.5, 152.5, 151.4, 146.7, 138.4, 135.4, 133.9, 130.6, 129.0, 123.3, 121.8, 96.1, 76.6, 39.9, 37.2, 31.4, 28.7, 28.5, 22.7, 19.6, 15.6, 14.2. HRMS (ESI): m/z [M+H]$^+$ Calcd for [$C_{24}H_{27}ClN_4S$ + H]$^+$ 439.1718, found 439.1723.

**3-(4-Butyl-5-((4-chlorobenzyl)thio)-4H-1,2,4-triazol-3-yl)-5-(oct-1-yn-1-yl)pyridine (31).** 1-(Bromomethyl)-4-chlorobenzene (36 mg, 1.2 equiv, 0.18 mmol) was reacted with 4-butyl-5-(5-(oct-1-yn-1-yl)pyridin-3-yl)-2,4-dihydro-3H-1,2,4-triazole-3-thione, **57** (50 mg, 1.0 equiv, 0.15 mmol) according to general procedure A1. Final product was purified by MPLC (mobile phase: 40-70 % EtOAc/Hex). Yield 75% (51 mg) as a yellow sticky solid. $^1$H NMR (400 MHz, $CDCl_3$) δ 8.71 (d, $J$ = 2.0 Hz, 1H), 8.66 (d, $J$ = 2.1 Hz, 1H), 7.90 (t, $J$ = 2.1 Hz, 1H), 7.36–7.24 (m, 5H), 4.48 (s, 2H), 3.83–3.75 (m, 2H), 2.44 (t, $J$ = 7.1 Hz, 2H), 1.68–1.56 (m, 2H), 1.56–1.40 (m, 5H), 1.40–1.29 (m, 5H), 1.24–1.08 (m, 2H), 0.96–0.87 (m, 3H), 0.82 (t, $J$ = 7.3 Hz, 3H). $^{13}$C NMR (126 MHz, $CDCl_3$) δ 153.5, 152.6, 151.7, 146.8, 138.5, 135.4, 133.9, 130.6, 129.0, 123.5, 121.8, 96.1, 76.6, 44.7, 37.4, 32.1, 31.5, 28.7, 28.5, 22.7, 19.7, 19.6, 14.2, 13.5. HRMS (ESI): m/z [M+H]$^+$ Calcd for [$C_{26}H_{31}ClN_4S$ + H]$^+$ 467.2036, found 467.2034.

**3-(5-((4-Chlorobenzyl)thio)-4-phenyl-4H-1,2,4-triazol-3-yl)-5-(oct-1-yn-1-yl)pyridine (32).** 1-(Bromomethyl)-4-chlorobenzene (73.5 mg, 1.2 equiv, 358 µmol) was reacted with 5-(5-(oct-1-yn-1-yl)pyridin-3-yl)-4-phenyl-2,4-dihydro-3H-1,2,4-triazole-3-thione, **58** (108 mg, 1.0 equiv, 298 µmol) according to general procedure A1. Final product was purified by MPLC (mobile phase: 20–70% EtOAc/Hex). Yield 39% (57 mg) as an off-white solid; mp 116–118 ºC. $^1$H NMR (400 MHz, $CDCl_3$) δ 8.53 (d, $J$ = 2.1 Hz, 1H), 8.31 (d, $J$ = 2.2 Hz, 1H), 7.84 (t, $J$ = 2.1 Hz, 1H), 7.55–7.42 (m, 3H), 7.34–7.19 (m, 4H), 7.15–7.06 (m, 2H), 4.44 (s, 2H), 2.38 (t, $J$ = 7.0 Hz, 2H), 1.62–1.51 (m, 2H), 1.47–1.36 (m, 2H), 1.36–1.25 (m, 3H), 0.90 (t, $J$ = 6.9 Hz, 3H). $^{13}$C NMR (101 MHz, $CDCl_3$) δ 153.3, 152.9, 152.2, 146.3, 137.8, 135.1, 133.8, 133.5, 130.7, 130.5, 130.4, 128.8, 127.2, 122.6, 121.4, 95.5, 76.6, 36.4, 31.4, 28.6, 28.5, 22.7, 19.5, 14.2. HRMS (ESI): m/z [M+H]$^+$ Calcd for [$C_{24}H_{27}ClN_4S$ + H]$^+$ 487.1723, found 487.1725.

**3-(4-Benzyl-5-((4-chlorobenzyl)thio)-4H-1,2,4-triazol-3-yl)-5-(oct-1-yn-1-yl)pyridine (33).** 1-Chloro-4-(chloromethyl)benzene (42 mg, 1.2 equiv, 0.26 mmol) was reacted with 4-benzyl-5-(5-(oct-1-yn-1-yl)pyridin-3-yl)-2,4-dihydro-3H-1,2,4-triazole-3-thione, **59** (82 mg, 1.0 equiv, 0.22 mmol) according to general procedure A1. Final product was purified by MPLC (mobile phase:

20–50 EtOAc/Hex). Yield 76% (83 mg) as a sticky solid. $^1$H NMR (400 MHz, $CDCl_3$) δ 8.65 (d, $J$ = 2.0 Hz, 1H), 8.53 (d, $J$ = 2.2 Hz, 1H), 7.79 (t, $J$ = 2.1 Hz, 1H), 7.36–7.19 (m, 9H), 6.89–6.80 (m, 2H), 5.04 (s, 2H), 4.44 (s, 2H), 2.41 (t, $J$ = 7.0 Hz, 2H), 1.66–1.53 (m, 2H), 1.50–1.38 (m, 2H), 1.37–1.27 (m, 4H), 0.92 (t, $J$ = 7.0 Hz, 3H). $^{13}$C NMR (101 MHz, $CDCl_3$) δ 153.5, 153.2, 152.3, 146.8, 138.5, 135.3, 134.4, 133.9, 130.7, 129.3, 129.0, 128.6, 126.2, 123.0, 121.7, 96.0, 76.5, 48.2, 37.4, 31.4, 28.7, 28.5, 22.7, 19.6, 14.2. HRMS (ESI): m/z [M+H]$^+$ Calcd for [$C_{29}H_{29}ClN_4S$ + H]$^+$ 501.1880, found 501.1878.

**3-(4-Cyclopropyl-5-((3,5-dinitrobenzyl)thio)-4H-1,2,4-triazol-3-yl)-5-(oct-1-yn-1-yl)pyridine (35).** 1-(Chloromethyl)-3,5-dinitrobenzene (49 mg, 1.2 equiv, 0.22 mmol) was reacted with **60** (61 mg, 1.0 equiv, 0.19 mmol) according to general procedure A2. Final product was purified by MPLC (mobile phase: 20–50 EtOAc/Hex). Yield 67% (63 mg) as a white solid; mp 119–121 ºC. $^1$H NMR (400 MHz, $CDCl_3$) δ 8.93 (t, $J$ = 2.1 Hz, 1H), 8.85 (d, $J$ = 2.1 Hz, 1H), 8.79–8.74 (m, 2H), 8.68 (d, $J$ = 2.0 Hz, 1H), 8.06 (t, $J$ = 2.1 Hz, 1H), 4.73 (s, 2H), 3.26–3.17 (m, 1H), 2.42 (t, $J$ = 7.1 Hz, 2H), 1.66–1.55 (m, 2H), 1.50–1.39 (m, 2H), 1.38–1.27 (m, 4H), 1.20–1.12 (m, 2H), 0.89 (t, $J$ = 7.0 Hz, 3H), 0.79–0.70 (m, 2H). $^{13}$C NMR (101 MHz, $CDCl_3$) δ 153.9, 153.3, 153.1, 148.6, 146.9, 142.3, 138.0, 129.6, 122.85, 121.5, 118.2, 95.9, 76.6, 34.4, 31.4, 28.7, 28.5, 25.8, 22.7, 19.6, 14.2, 9.3. HRMS (ESI): m/z [M+H]$^+$ Calcd for [$C_{25}H_{26}N_6O_4S$ + H]$^+$ 507.1815, found 507.1813.

**3-(5-((4-Chlorobenzyl)thio)-4-methyl-4H-1,2,4-triazol-3-yl)pyridine (37).** 1-(Bromomethyl)-4-chlorobenzene (69 mg, 1.2 equiv, 0.34 mmol) was reacted with 4-Methyl-5-(pyridin-3-yl)-2,4-dihydro-3H-1,2,4-triazole-3-thione (54 mg, 1.0 equiv, 0.28 mmol) according to general procedure A1. Final product was purified by MPLC (mobile phase: 2–5% MeOH/ $CH_2Cl_2$). Yield 90% (80 mg) as a white solid; mp 138–140 ºC. $^1$H NMR (400 MHz, $CDCl_3$) δ 9.04–8.65 (m, 2H), 8.06 (dt, $J$ = 8.0, 1.8 Hz, 1H), 7.56–7.48 (m, 1H), 7.34–7.23 (m, 4H), 4.43 (s, 2H), 3.46 (s, 3H). $^{13}$C NMR (101 MHz, $CDCl_3$) δ 153.2, 152.1, 150.45, 148.1, 136.9, 135.4, 134.0, 130.6, 129.0, 124.3, 124.1, 37.5, 31.8. HRMS (ESI): m/z [M+H]$^+$ Calcd for [$C_{23}H_{25}ClN_4S$ + H]$^+$ 317.0631, found 317.0627.

**3-Bromo-5-(5-((4-chlorobenzyl)thio)-4-methyl-4H-1,2,4-triazol-3-yl)pyridine (38).** To 1-(bromomethyl)-4-chlorobenzene (0.9 g, 1.2 equiv, 4 mmol) and 1,2,4-triazole-3-thiol (1.0 equiv) dissolved in 15 mL MeOH/acetone (1:2) was added $K_2CO_3$ (1.52.0 equiv). The mixture was stirred overnight at room temperature. Upon completion of the reaction, as determined by TLC, 5 mL of water was added, leading to the formation of white precipitates. The precipitate was dissolved with EtOAc (30 mL), washed with water brine, and dried over $Na_2SO_4$, and then concentrated under low pressure. The resultant residue was then purified by MPLC. Yield 70% (1 g) as white solid; mp 160–161 ºC. $^1$H NMR (400 MHz, $CDCl_3$) δ 8.86 – 8.71 (m, 2H), 8.16 (s, 1H), 7.40 – 7.18 (m, 4H), 4.44 (s, 2H), 3.47 (s, 3H). $^{13}$C NMR (101 MHz, $CDCl_3$) δ 152.3, 152.3, 152.2, 146.7, 138.5, 135.3, 134.0, 130.6, 129.0, 124.9, 121.2, 37.4, 31.8. Calcd for [$C_{15}H_{12}BrClN_4S$ + H]$^+$ 395.9733, found 394.9704.

**3-(5-((4-Chlorobenzyl)thio)-4-methyl-4H-1,2,4-triazol-3-yl)-5-ethynylpyridine (39).** To 3-(5-((4-chlorobenzyl)thio)-4-methyl-4H-1,2,4-triazol-3-yl)-5-((trimethylsilyl)ethynyl)pyridine (20 mg, 1.0 equiv, 48 μmol) dissolved in MeOH (1.5 mL) was added $K_2CO_3$ (67 mg, 10 equiv, 0.48 mmol) and stirred at room temperature for 4 h. On completion of the reaction, the solvent was evaporated under reduced pressure and the residue was dissolved in EtOAc and then washed with water and brine. The organic layer was dried under $Na_2SO_4$ and concentrated under reduced pressure. The residue was purified by MPLC (50–80% EtOAc/Hex). Yield 91% (15 mg) as a white solid; mp 107–109 ºC. $^{1}$H NMR (400 MHz, $CD_3OD$ ) δ 8.83–8.77 (m, 1H), 8.18 (t, *J* = 2.1 Hz, 1H), 7.36–7.26 (m, 2H), 4.37 (s, 1H), 3.54 (s, 2H). $^{13}$C NMR (126 MHz, $CD_3OD$) δ 154.6, 154.2, 153.5, 149.1, 140.3, 137.1, 134.8, 131.7, 129.8, 124.5, 121.5, 84.3, 79.9, 38.5, 32.5. HRMS (ESI): m/z $[M+H]^+$ Calcd for $[C_{17}H_{13}ClN4S + H]^+$ 341.0628, found 341.0657.

**3-(5-((4-Chlorobenzyl)thio)-4-methyl-4H-1,2,4-triazol-3-yl)-5-(pent-1-yn-1-yl)pyridine (40).** 1-(Bromomethyl)-4-chlorobenzene (76 mg, 1.2 equiv, 0.37 mmol) was reacted with 4-methyl-5-(5-(pent-1-yn-1-yl)pyridin-3-yl)-2,4-dihydro-3H-1,2,4-triazole-3-thione (80 mg, 1.0 equiv, 0.31 mmol) according to general procedure A1. Final product was purified by recrystallization in methanol. Yield 92% (109 mg) as light-yellow crystal; mp 141–142 ºC. $^{1}$H NMR (400 MHz, $CDCl_3$) δ 8.72 (s, 2H), 7.94 (d, *J* = 2.0 Hz, 1H), 7.34–7.24 (m, 4H), 4.43 (s, 2H), 3.44 (s, 3H), 2.43 (t, *J* = 7.0 Hz, 2H), 1.66 (h, *J* = 7.2 Hz, 2H), 1.06 (t, *J* = 7.4 Hz, 3H). $^{13}$C NMR (101 MHz, $CDCl_3$) δ 153.3, 152.9, 152.0, 146.6, 138.3, 135.4, 133.9, 130.6, 129.0, 123.0, 121.8, 96.0, 76.6, 37.5, 31.8, 22.0, 21.5, 13.7. HRMS (ESI): m/z $[M+H]^+$ Calcd for $[C_{20}H_{19}ClN4S + H]^+$ 383.1097, found 383.1106.

**5-(5-((4-Chlorobenzyl)thio)-4-methyl-4H-1,2,4-triazol-3-yl)-2-(oct-1-yn-1-yl)pyridine (41).** 1-(Bromomethyl)-4-chlorobenzene (33 mg, 1.2 equiv, 0.16 mmol) was reacted with 4-Methyl-5-(6-(oct-1-yn-1-yl)pyridin-3-yl)-2,4-dihydro-3H-1,2,4-triazole-3-thione (40 mg, 1.0 equiv, 0.13 mmol) according to general procedure A1. Final product was purified by MPLC (mobile phase: 50–100% EtOAc/Hex). Yield 87% (49 mg) as a white solid; mp 140–141 ºC $^{1}$H NMR (500 MHz, $CDCl_3$) δ 8.76 (d, *J* = 2.4 Hz, 1H), 7.93 (dd, *J* = 8.1, 2.3 Hz, 1H), 7.50 (d, *J* = 8.1 Hz, 1H), 7.37–7.19 (m, 5H), 4.42 (s, 2H), 3.43 (s, 3H), 2.47 (t, *J* = 7.2 Hz, 2H), 1.71–1.59 (m, 2H), 1.52–1.40 (m, 2H), 1.37–1.26 (m, 4H), 0.90 (t, *J* = 7.1 Hz, 3H). $^{13}$C NMR (101 MHz, $CDCl_3$) δ 153.3, 151.9, 148.7, 145.4, 136.0, 135.4, 133.9, 130.6, 128.9, 126.8, 121.8, 94.0, 80.1, 37.5, 31.8, 31.4, 28.8, 28.3, 22.6, 19.6, 14.2. HRMS (ESI): m/z $[M+H]^+$ Calcd for $[C_{23}H_{25}ClN_4S + H]^+$ 425.1567, found 425.1565.

**3-((4-Chlorobenzyl)thio)-4-methyl-5-(3-(oct-1-yn-1-yl)phenyl)-4H-1,2,4-triazole (42).** 1-(Bromomethyl)-4-chlorobenzene (29 mg, 1.2 equiv, 0.14 mmol) was reacted with 4-methyl-5-(3-(oct-1-yn-1-yl)phenyl)-2,4-dihydro-3H-1,2,4-triazole-3-thione (35 mg, 1.0 equiv, 0.12 mmol) according to general procedure A1. Final product was purified by MPLC (mobile phase: 20–50% EtOAc/Hex). Yield 61% (30 mg) as a white solid; mp 58–60 ºC. $^{1}$H NMR (400 MHz, $CDCl_3$) δ 7.60–7.55 (m, 1H), 7.54–7.37 (m, 3H), 7.27 (d, *J* = 0.6 Hz, 4H), 4.40 (s, 2H), 3.38 (s, 3H), 2.41 (t, *J* = 7.1 Hz, 2H), 1.66–1.55 (m, 2H), 1.51–1.40 (m, 2H), 1.40–1.27 (m, 4H), 0.90 (t, *J* = 7.1 Hz, 3H). $^{13}$C NMR (101 MHz, $CDCl_3$) δ 155.7, 151.0, 135.7, 133.9, 133.3, 131.5, 130.6, 129.1, 129.0, 127.7, 127.2, 125.3, 92.2, 79.7, 37.7, 31.7, 31.5, 28.8, 28.7, 22.7, 19.6, 14.2. HRMS (ESI): m/z $[M+H]^+$ Calcd for $[C_{24}H_{26}ClN_3S + Cs]^+$ 556.0590, found 556.0590.

**3-(5-((4-Chlorobenzyl)thio)-4-methyl-4H-1,2,4-triazol-3-yl)-5-(octyloxy)pyridine (43).** 1-(Bromomethyl)-4-chlorobenzene (59 mg, 1.1 equiv, 0.29 mmol) was reacted with 4-methyl-5-(5-(octyloxy)pyridin-3-yl)-2,4-dihydro-3H-1,2,4-triazole-3-thione (84 mg, 1.0 equiv, 0.26 mmol) according to general procedure A2. Final product was purified by MPLC (mobile phase: 50–80 EtOAc/Hex). Yield 61% (71 mg) as a white solid; mp 76–78 ºC. $^{1}$H NMR (400 MHz, $CDCl_3$) δ 8.47–8.29 (m, 2H), 7.54–7.45 (m, 1H), 7.35–7.21 (m, 4H), 4.42 (s, 2H), 4.05 (t, *J* = 6.5 Hz, 2H),

3.44 (s, 3H), 1.90–1.73 (m, 2H), 1.53–1.41 (m, 2H), 1.41–1.21 (m, 8H), 0.94–0.83 (m, 3H). $^{13}$C NMR (101 MHz, $CDCl_3$) δ 155.4, 153.5, 151.7, 140.2, 140.1, 135.5, 133.8, 130.5, 128.9, 123.8, 120.4, 68.8, 37.5, 31.8, 31.8, 29.3, 29.3, 29.1, 26.0, 22.7, 14.2. HRMS (ESI): m/z [M+H]$^+$ Calcd for [$C_{23}H_{29}ClN_4OS$ + H]$^+$ 445.1829, found 445.1827.

**3-((4-Chlorobenzyl)thio)-4-methyl-5-(4-octylphenyl)-4H-1,2,4-triazole (45).** 1-(Bromomethyl)-4-chlorobenzene (41 mg, 1.2 equiv, 0.20 mmol) was reacted with 4-Methyl-5-(4-octylphenyl)-4H-1,2,4-triazole-3-thione, **66** (50 mg, 1.0 equiv, 0.16 mmol) according to general procedure A1. Final product was purified by flash chromatography (mobile phase: 20–50 EtOAc/Hex). Yield 92% (65 mg) as a white solid; mp 136–138 ºC. $^1$H NMR (400 MHz, $CDCl_3$) δ 7.52–7.45 (m, 2H), 7.33–7.22 (m, 6H), 4.38 (s, 2H), 3.36 (s, 3H), 2.71–2.61 (m, 2H), 1.70–1.56 (m, 2H), 1.41–1.19 (m, 10H), 0.92–0.83 (m, 3H). $^{13}$C NMR (101 MHz, $CDCl_3$) δ 156.3, 150.6, 145.5, 135.7, 133.8, 130.6, 129.1, 128.9, 128.5, 124.4, 37.8, 35.9, 32.0, 31.7, 31.4, 29.5, 29.36, 29.35, 22.8, 14.2. HRMS (ESI): m/z [M+H]$^+$ Calcd for [$C_{24}H_{30}ClN_3S$ + H]$^+$ 428.1927, found 428.1949.

**3-(5-((4-Chlorobenzyl)thio)-1-methyl-1H-pyrrol-2-yl)-5-(oct-1-yn-1-yl)pyridine (46).** To a 1 dram vial was added 3-(5-bromo-1-methyl-1H-pyrrol-2-yl)-5-(oct-1-yn-1-yl)pyridine, **69** (18 mg, 1 equiv, 52 µmol), DIPEA (13 mg, 18 µL, 2.0 equiv, 0.10 mmol), dry 1,4-dioxane (1 mL). The mixture was then evacuated and backfilled with argon three times. Pd2(dba)3 (2.4 mg, 0.05 equiv, 2.6 µmol), xantphos (3.0 mg, 0.1 equiv, 5.2 µmol) and the 4-Chlorobenzyl mercaptan (8.3 mg, 6.9 µL, 1.0 equiv, 52 µmol) were added to the reaction mixture. The mixture was then degassed twice and reflux for 15 h. On completion of the reaction, as determined by TLC, the reaction mixture was allowed to reach ambient temperature, filtered, and concentrated under reduced pressure. The crude was subjected to MPLC (mobile phase: 2–20 EtOAc/Hex) to obtain final product. Yield 59% (13 mg) as a brown viscous oil. $^1$H NMR (500 MHz, $CDCl_3$) δ 8.54 (d, $J$ = 2.0 Hz, 1H), 8.44 (d, $J$ = 2.2 Hz, 1H), 7.58 (t, $J$ = 2.1 Hz, 1H), 7.2 –7.20 (m, 2H), 7.02–6.95 (m, 2H), 6.43 (d, $J$ = 3.7 Hz, 1H), 6.26 (d, $J$ = 3.7 Hz, 1H), 3.77 (s, 2H), 3.27 (s, 3H), 2.44 (t, $J$ = 7.1 Hz, 2H), 1.71–1.55 (m, 2H), 1.52–1.40 (m, 2H), 1.40–1.27 (m, 4H), 0.91 (t, $J$ = 6.9 Hz, 3H). $^{13}$C NMR (126 MHz, $CDCl_3$) δ 150.7, 147., 138.0, 136.9, 133.7, 133.3, 130.4, 128.9, 128.7, 123.4, 121.1, 118.7, 110.2, 94.8, 77.2, 42.7, 32.3, 31.5, 28.8, 28.6, 22.7, 19.5, 14.2. HRMS (ESI): m/z [M+H]$^+$ Calcd for [$C_{25}H_{27}ClN_2S$ + H]$^+$ 423.1662, found 423.1654.

**2-((4-Chlorobenzyl)thio)-5-(5-(oct-1-yn-1-yl)pyridin-3-yl)-1,3,4-oxadiazole (47).** 1-(Bromomethyl)-4-chlorobenzene (50 mg, 1.5 equiv, 0.29 mmol) was reacted with 5-(5-(oct-1-yn-1-yl)pyridin-3-yl)-1,3,4-oxadiazole-2-thiol, **70** (70 mg, 1.0 equiv, 0.24 mmol) $K_2CO_3$ (50 mg, 1.5 equiv, 0.37 mmol) according to general procedure A1. Final product was purified by MPLC (mobile phase: 10–50 EtOAc/Hex). Yield 76% (76 mg) as a white solid; mp 75–76 ºC. $^1$H NMR (400 MHz, $CDCl_3$) δ 9.04 (d, $J$ = 2.1 Hz, 1H), 8.71 (d, $J$ = 2.0 Hz, 1H), 8.22 (t, $J$ = 2.1 Hz, 1H), 7.45–7.36 (m, 2H), 7.35–7.26 (m, 2H), 4.49 (s, 2H), 2.44 (t, $J$ = 7.1 Hz, 2H), 1.69 – 1.55 (m, 2H), 1.52–1.39 (m, 2H), 1.42–1.25 (m, 4H), 0.95–0.84 (m, 3H). $^{13}$C NMR (101 MHz, $CDCl_3$) δ 164.7, 163.5, 154.7, 145.5, 136.0, 134.3, 134.1, 130.7, 129.1, 121.9, 119.6, 77.5, 76.4, 36.2, 31.4, 28.7, 28.5, 22.7, 19.6, 14.2. HRMS (ESI): m/z [M+H]$^+$ Calcd for [$C_{22}H_{22}ClN_3OS$ + H]$^+$ 412.1250, found 412.1216.

**2-((4-Chlorobenzyl)thio)-5-(5-(oct-1-yn-1-yl)pyridin-3-yl)-1,3,4-thiadiazole (48).**To a solution of 1-chloro-4-(chloromethyl)benzene (20 mg, 1.0 equiv, 0.12 mmol) and tetrabutylammonium bromide (2.0 mg, 0.05 equiv, 6.2 µmol) dissolved in $CH_2Cl_2$ (1.5 mL) was added a solution of 5-(5-(oct-1-yn-1-yl)pyridin-3-yl)-1,3,4-thiadiazole-2-thiol, **71** (41 mg, 1.1 equiv, 0.14 mmol) with NaOH (5.0 mg, 1.0 equiv, 0.12 mmol) dissolved in $H_2O$ (1 mL). The reaction was stirred at room temperature for 12 h. The organic phase was separated and washed with water 2 x 10 mL), dried over $Na_2SO_4$ and then concentrated under reduced pressure. The residue was purified by flash chromatography (mobile phase: 10–50 EtOAc/Hex). Yield 76% (76 mg) as a white solid; mp 72–74 ºC. $^1$H NMR (400 MHz, $CDCl_3$) δ 8.94–8.89 (m, 1H), 8.70–8.65 (m, 1H), 8.16 (t, *J* = 2.1 Hz, 1H), 7.45–7.36 (m, 2H), 7.35–7.27 (m, 2H), 4.58 (s, 2H), 2.45 (t, *J* = 7.1 Hz, 2H), 1.72–1.57 (m, 2H), 1.52–1.40 (m, 2H), 1.41–1.27 (m, 4H), 0.91 (t, *J* = 7.0 Hz, 3H). $^{13}$C NMR (101 MHz, $CDCl_3$) δ 165.5, 164.9, 154.2, 146.5, 136.9, 134.5, 134.1, 130.7, 129.1, 125.7, 122.0, 96.2, 76.5, 37.5, 31.5, 28.7, 28.5, 22.7, 19.6, 14.2. HRMS (ESI): m/z $[M + Cs]^+$ Calcd for $[C_{22}H_{22}ClN_3S_2 + H]^+$ 559.9998.1250, found 559.9962.

**3-(1-(4-Chlorophenethyl)-1H-1,2,3-triazol-4-yl)-5-(oct-1-yn-1-yl)pyridine (49).** A solution of 3-ethynyl-5-(oct-1-yn-1-yl)pyridine (60 mg, 1 equiv, 0.28 mmol) 1-(2-azidoethyl)-4-chlorobenzene (52 mg, 1 equiv, 0.28 mmol), sodium ascorbate (8.4 mg, 0.15 equiv, 43 µmol) and $CuSO_4\cdot 5H_2O$ (3.8 mg, 0.015 mmol) in $CH_3CN$ (1 mL) was stirred at room temperature under the argon protection for 16 h. The mixture was poured into water (5mL) and with short celite pad. The filtrate was then extracted with EtOAc (2 x 10 mL). The combined organic phase was washed with water, brine, dried over $Na_2SO4$, and concentrated under reduced pressure. The final product was purified by flash chromatography (mobile phase: 10–50% EtOAc/Hex). Yield 91% (102 mg) as a white solid; mp 132–134 ºC. $^1$H NMR (400 MHz, $CDCl_3$) δ 8.80 (d, *J* = 2.1 Hz, 1H), 8.55 (d, *J* = 2.0 Hz, 1H), 8.12 (t, *J* = 2.1 Hz, 1H), 7.56 (s, 1H), 7.30–7.23 (m, 2H), 7.08–7.00 (m, 2H), 4.63 (t, *J* = 7.1 Hz, 2H), 3.24 (t, *J* = 7.1 Hz, 2H), 2.43 (t, *J* = 7.1 Hz, 2H), 1.67–1.56 (m, 2H), 1.52–1.40 (m, 2H), 1.40–1.25 (m, 4H), 0.91 (t, *J* = 7.1 Hz, 3H). $^{13}$C NMR (101 MHz, $CDCl_3$) δ 151.7, 145.1, 144.2, 135.4, 135.4, 133.3, 130.2, 129.2, 126.1, 121.5, 120.5, 94.9, 51.8, 36.2, 31.5, 28.7, 28.6, 22.7, 19.6, 14.2. HRMS (ESI): m/z $[M+H]^+$ Calcd for $[C_{23}H_{25}ClN_4 + H]^+$ 393.1846, found 393.1834.

**5-((4-Chlorobenzyl)thio)-5'-(non-1-yn-1-yl)-3,3'-bipyridine (50).** To a round bottom flask was added 3-bromo-5-(non-1-yn-1-yl)pyridine (90.4 mg, 1.0 equiv, 323 µmol), 3-((4-chlorobenzyl)thio)-5-(4,4,5,5-tetramethyl-1,3,2-dioxaborolan-2-yl)pyridine (175 mg, 1.5 equiv, 484 µmol), potassium carbonate (44.6 mg, 1.0 equiv, 323 µmol) ,1,4-dioxane (2.5 mL) and water (0.5 mL). The reaction mixture was then evacuated and backfilled with argon three times. Subsequently, the reaction mixture was stirred at 85℃ under argon atmosphere for 18 h. Thereafter, the reaction was allowed to cool down to room temperature and diluted with water. The mixture was then extracted with EtOAc (2 x10 mL). The combined organic layer was subsequently washed with brine, dried over $Na_2SO_4$, and concentrated under reduced pressure. The crude was subjected to flash chromatography (mobile phase: 10-50 EtOAc/Hex) to obtain final product. Yield 63% (88 mg) as a white solid; mp 53-55 ºC. $^1$H NMR (400 MHz, $CDCl_3$) δ 8.69–8.58 (m, 3H), 8.54 (d, *J* = 2.1 Hz, 1H), 7.75 (t, *J* = 2.1 Hz, 1H), 7.67 (t, *J* = 2.1 Hz, 1H), 7.32–7.17 (m, 4H), 4.12 (s, 2H), 2.46 (t, *J* = 7.1 Hz, 2H), 1.69–1.59 (m, 2H), 1.53–1.42 (m, 2H), 1.40–1.27 (m, 4H), 0.92 (t, *J* = 6.9 Hz, 3H). $^{13}$C NMR (101 MHz, $CDCl_3$) δ 152.1, 150.6, 146.3, 146.2, 136.9, 136.5, 135.3, 133.7, 133.1, 132.3, 130.4, 129.0, 121.7, 95.4, 77.0, 38.6, 31.5, 28.8,

28.6, 22.7, 19.6, 14.2. HRMS (ESI): m/z $[M+H]^+$ Calcd for $[C_{26}H_{27}ClN_2S + H]^+$ 423.1662, found 423.1654.

**S-(4-Chlorobenzyl) (5-(oct-1-yn-1-yl)pyridin-3-yl)carbamothioate (51).** To a solution of 5-(oct-1-yn-1-yl)pyridin-3-amine, **77** (55 mg, 1.0 equiv, 0.27 mmol) in $CH_2Cl_2$ (2 mL) was added dropwise a solution of triphosgene (81 mg, 1.0 equiv, 0.27 mmol) in $CH_2Cl_2$ at 0 °C while stirring under argon. $Et_3N$ (28 mg, 38 µL, 1 equiv, 0.27 mmol) in $CH_2Cl_2$ was then added dropwise. The reaction mixture was stirred for 30 mins at 0 °C and then brought to room temperature and stirred for another 4 h. On the consumption of the starting material as determined by TLC, 4-Chlorobenzyl mercaptan (43 mg, 35 µL, 1.0 equiv, 0.27 mmol), and $Et_3N$ (28 mg, 38 µL, 1.0 equiv, 0.27 mmol) dissolved in $CH_2Cl_2$ were added dropwise. The reaction was stirred at room temperature overnight. The reaction was quenched with the addition of water (5) mL and the mixture extracted with $CH_2Cl_2$ (10 mL x 3). The combined organic phase was washed with water, brine, dried over $Na_2SO_4$, and concentrated under reduced pressure. The crude was subjected to MPLC (mobile phase: 0–5% MeOH/$CH_2Cl_2$) to obtain the final product. Yield 36% (38 mg) as a white solid; mp 137–139 ºC. $^1$H NMR (400 MHz, $CDCl_3$) δ 8.44 (s, 1H), 8.40–8.31 (m, 2H), 8.09 (t, $J$ = 2.2 Hz, 1H), 7.31–7.23 (m, 4H), 4.17 (s, 2H), 2.40 (t, $J$ = 7.1 Hz, 2H), 1.65–1.54 (m, 2H), 1.48–1.38 (m, 2H), 1.37–1.27 (m, 4H), 0.90 (t, $J$ = 7.2 Hz, 3H). $^{13}$C NMR (101 MHz, $CDCl_3$) δ 165.8, 147.6, 138.7, 136.5, 134.6, 133.4, 130.3, 129.7, 128.9, 121.9, 95.2, 76.9, 33.9, 31.5, 28.8, 28.6, 22.7, 19.6, 14.2. HRMS (ESI): m/z $[M+H]^+$ Calcd for $[C_{21}H_{23}ClN_2OS + H]^+$ 387.1298, found 387.1290.

**1-(4-Chlorobenzyl)-3-(5-(oct-1-yn-1-yl)pyridin-3-yl)thiourea (52).** To a solution of 3-isothiocyanato-5-(oct-1-yn-1-yl)pyridine (160 mg, 1 equiv, 655 µmol) in $CH_2Cl_2$ (1.5 mL) was added (4-chlorophenyl)methanamine (92.7 mg, 77 µL, 1.0 equiv, 655 µmol) in $CH_2Cl_2$ (0.5 mL) dropwise at 0 °C. Subsequently, $Et_3N$ (66.3 mg, 91.3 µL, 1.0 equiv, 655 µmol) in $CH_2Cl_2$ (0.5 mL) was added dropwise to the reaction mixture. The mixture was then stirred for 30 min and then for another 4 h at room temperature. On the completion of the reaction as determined by TLC, the reaction mixture was diluted with 4 mL of water and extracted with $CH_2Cl_2$ (2x 10 mL). The combined organic layer was subsequently washed with brine, dried over $Na_2SO_4$, and concentrated under reduced pressure. The crude was subjected to flash chromatography (mobile phase: 5% MeOH/ $CH_2Cl_2$). Yield 68% (172 mg) as a white solid; mp 144–145 ºC. $^1$H NMR (400 MHz, $CDCl_3$) δ 8.43 (d, $J$ = 1.8 Hz, 1H), 8.34 (d, $J$ = 2.5 Hz, 1H), 8.09 (s, 1H), 7.61 (t, $J$ = 2.2 Hz, 1H), 7.34–7.21 (m, 4H), 6.36 (s, 1H), 4.82 (d, $J$ = 5.6 Hz, 2H), 2.42 (t, $J$ = 7.1 Hz, 2H), 1.66–1.55 (m, 2H), 1.49–1.39 (m, 2H), 1.38–1.26 (m, 4H), 0.91 (t, $J$ = 7.1 Hz, 3H). $^{13}$C NMR (101 MHz, $CDCl_3$) δ 181.6, 150.6, 144.5, 135.5, 134.9, 134.0, 132.8, 129.3, 129.2, 122.5, 96.5, 76.3, 48.8, 31.4, 28.8, 28.5, 22.7, 19.6, 14.2. HRMS (ESI): m/z $[M+H]^+$ Calcd for $[C_{21}H_{24}ClN_3S + H]^+$ 386.1458, found 386.1433.

**3-(((5-(4-Chlorophenyl)-4-methyl-4H-1,2,4-triazol-3-yl)thio)methyl)-5-(oct-1-yn-1-yl)pyridine (53).** To (5-(Oct-1-yn-1-yl)pyridin-3-yl)methanol (30 mg, 1 equiv, 0.14 mmol) dissolved in anhydrous THF (30 mL) was added triphenylphosphine (43 mg, 1.2 equiv, 0.17 mmol) and 5-(4-chlorophenyl)-4-methyl-2,4-dihydro-3H-1,2,4-triazole-3-thionethiol (31 mg, 1.0 equiv, 0.14 mmol) at 0 °C under argon. After stirring for 30 min, DIAD (28 mg, 27 µL, 1.0 equiv, 0.14 mmol) was added dropwise. The mixture was then warmed to room temperature and stirred for 3 h. Solvent was removed in vacuo, and the residue was purified by MPLC (40–70% EtOAc/Hex) to afford the final product. Yield 29% (17 mg) as a white solid; mp 146–148 ºC. $^1$H NMR (400

MHz, $CDCl_3$) δ 8.51 (d, *J* = 2.0 Hz, 1H), 8.47 (d, *J* = 2.2 Hz, 1H), 7.64 (t, *J* = 2.1 Hz, 1H), 7.60–7.43 (m, 4H), 4.41 (s, 2H), 3.38 (s, 3H), 2.39 (t, *J* = 7.1 Hz, 2H), 1.64–1.52 (m, 2H), 1.48–1.38 (m, 2H), 1.37–1.26 (m, 4H), 0.90 (t, *J* = 6.9 Hz, 3H). $^{13}$C NMR (126 MHz, $CDCl_3$) δ 155.4, 151.7, 150.9, 148.4, 138.9, 136.7, 132.5, 130.0, 129.4, 125.5, 121.2, 95.1, 77.0, 35.0, 31.7, 31.5, 28.8, 28.6, 22.7, 19.6, 14.2. HRMS (ESI): m/z $[M+H]^+$ Calcd for $[C_{23}H_{25}ClN_4S + H]^+$ 425.1567, found 425.1567.

#### 3. $^{1}$H and $^{13}$C NMR spectrum of the selected compounds (1, 8, 9, 18, 20, 21, 24, 34, 36, 44 and 80)

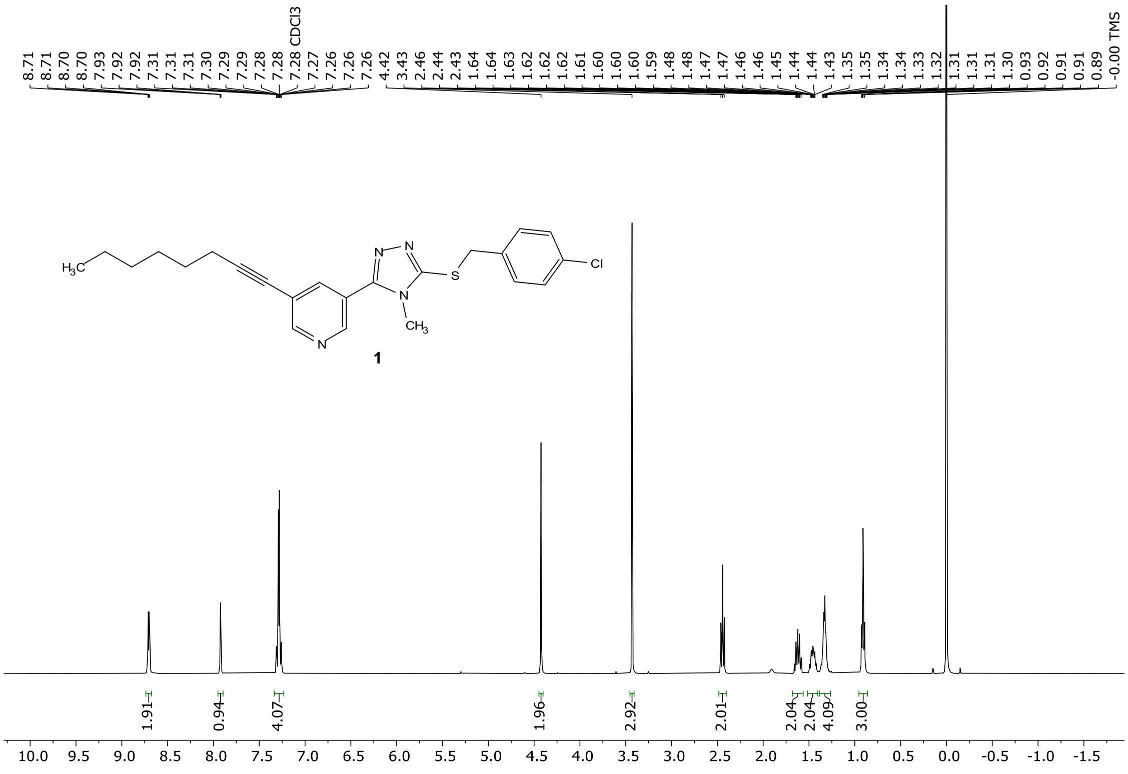

The $^{1}$H NMR spectrum of compound **1**

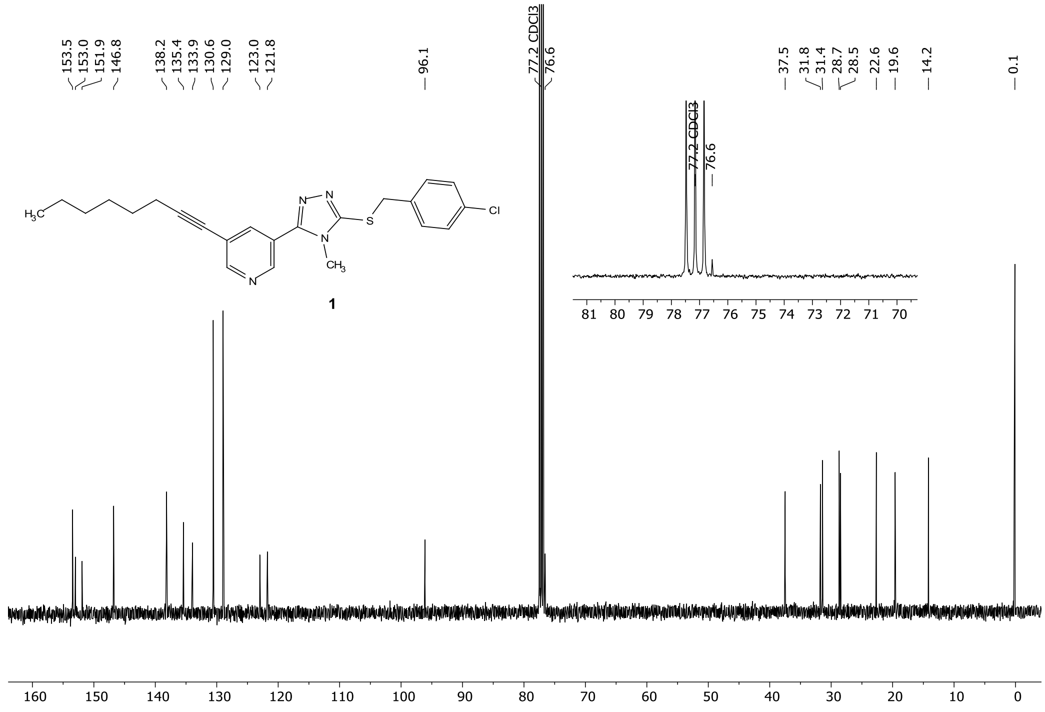

The $^{13}$C NMR spectrum of compound **1**

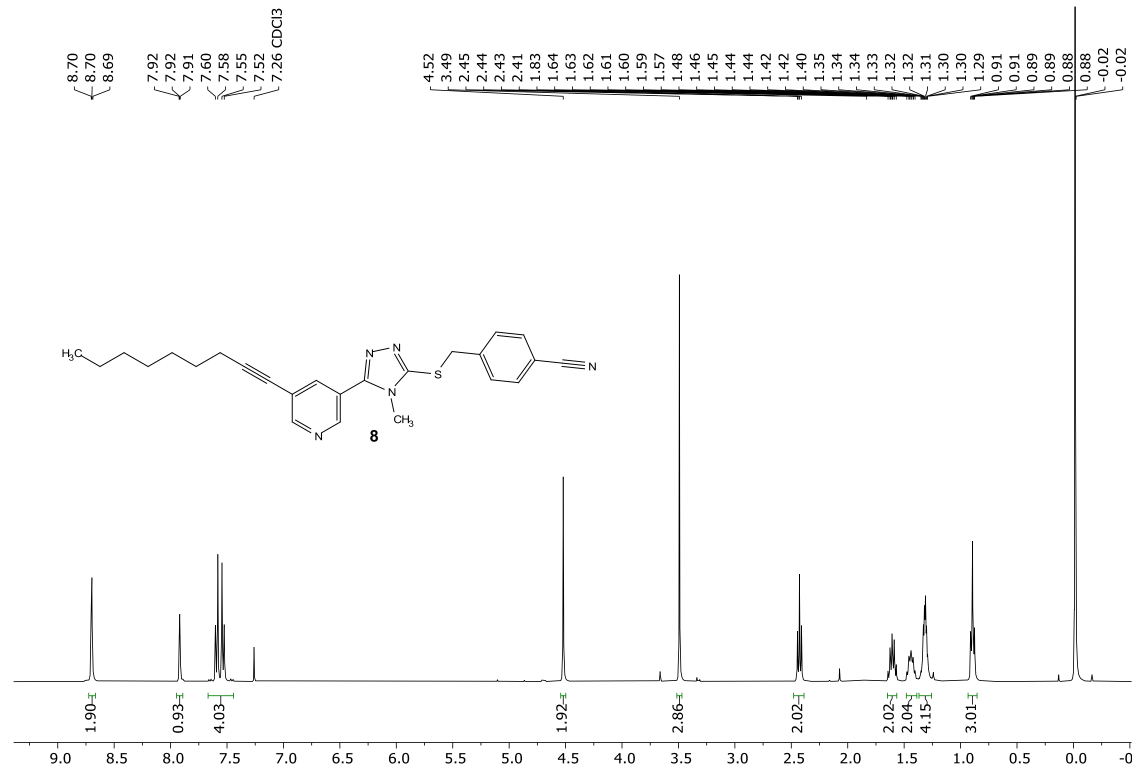

The $^{1}H$ NMR spectrum of compound **8**

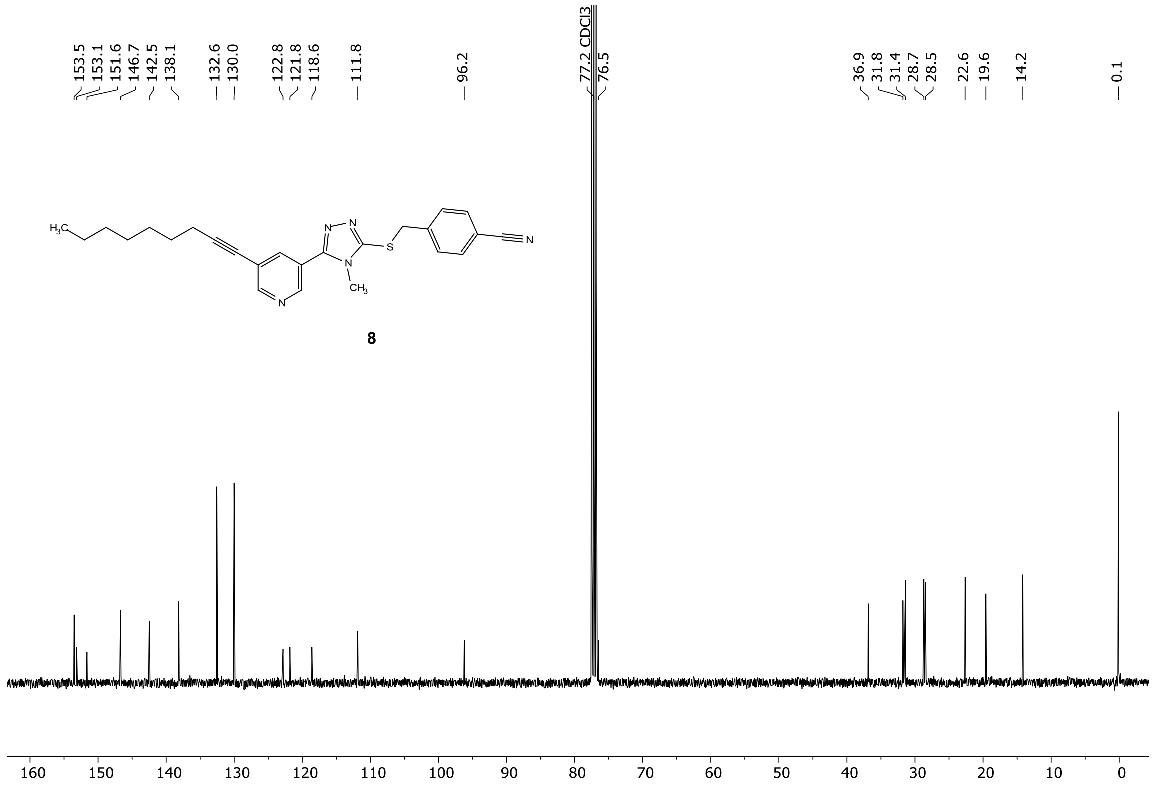

The $^{13}C$ NMR spectrum of compound **8**

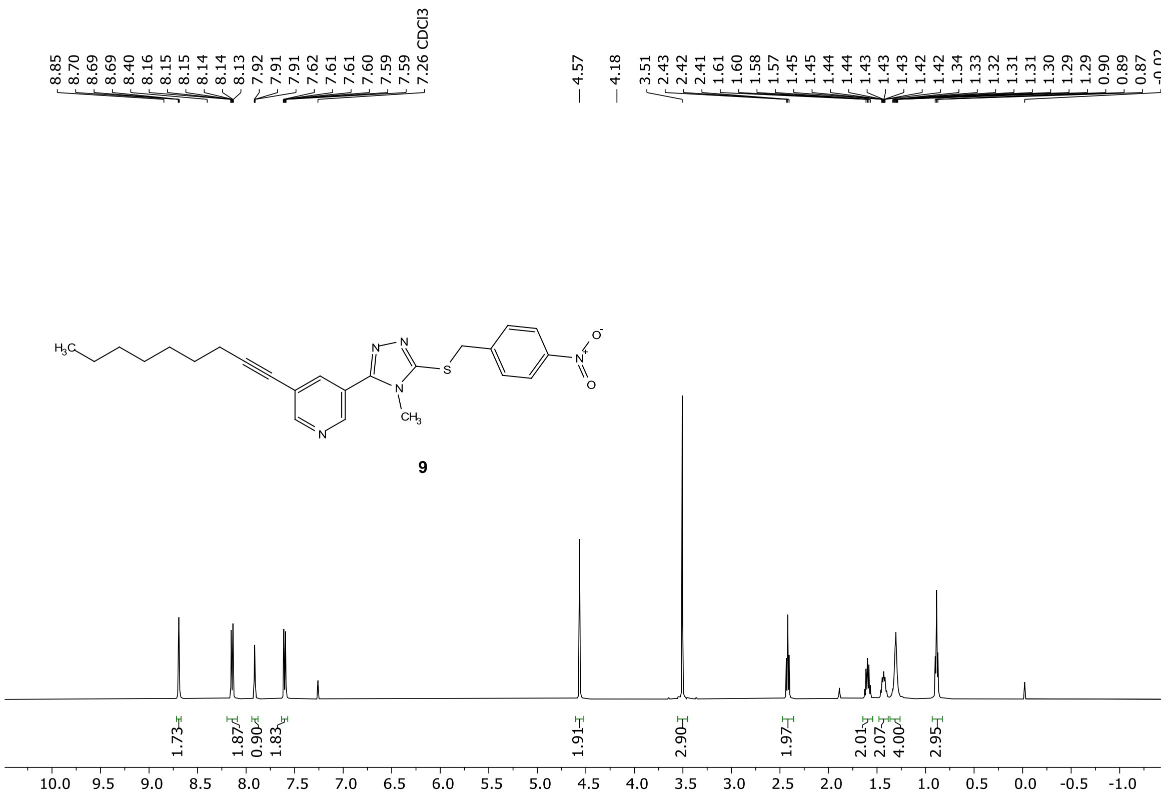

The $^{1}H$ NMR spectrum of compound **9**

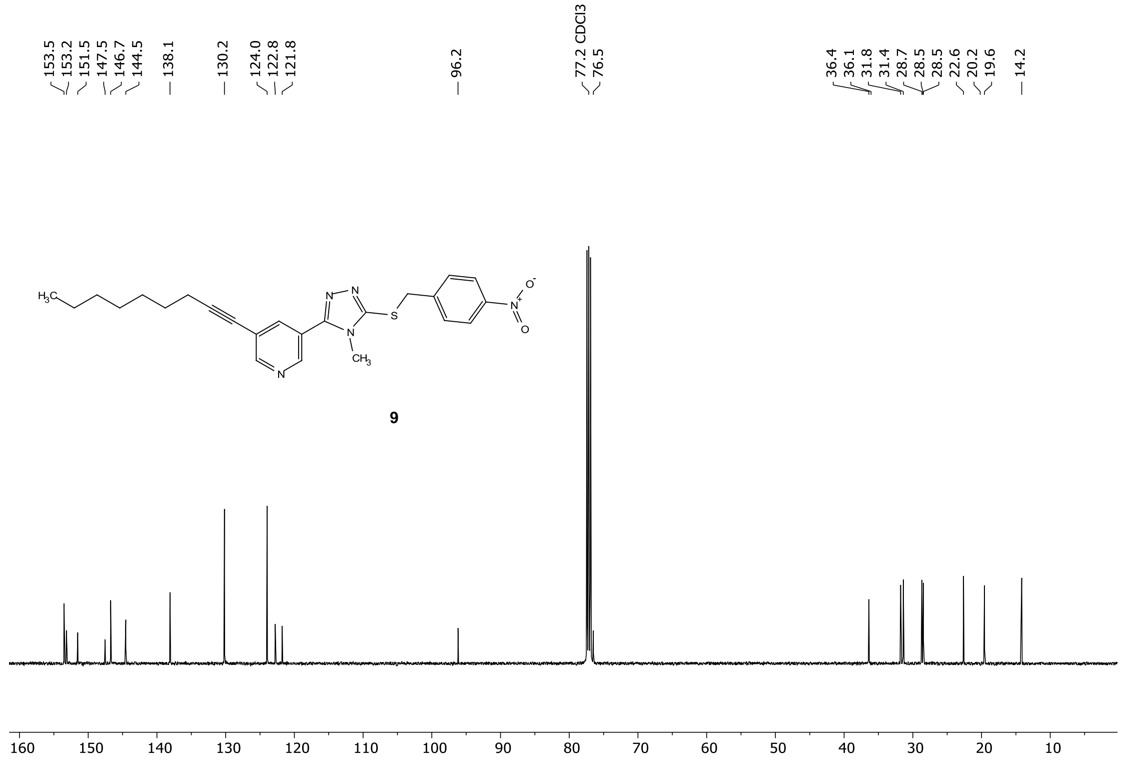

The $^{13}C$ NMR spectrum of compound **9**

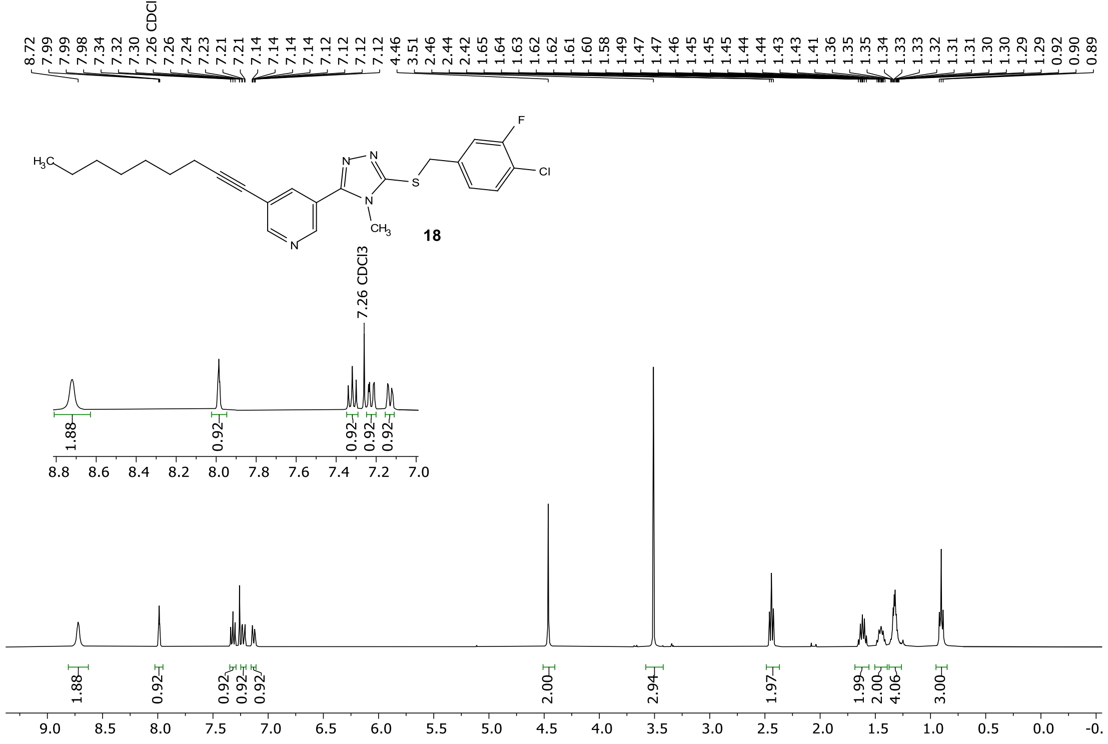

The $^{1}H$ NMR spectrum of compound **18**

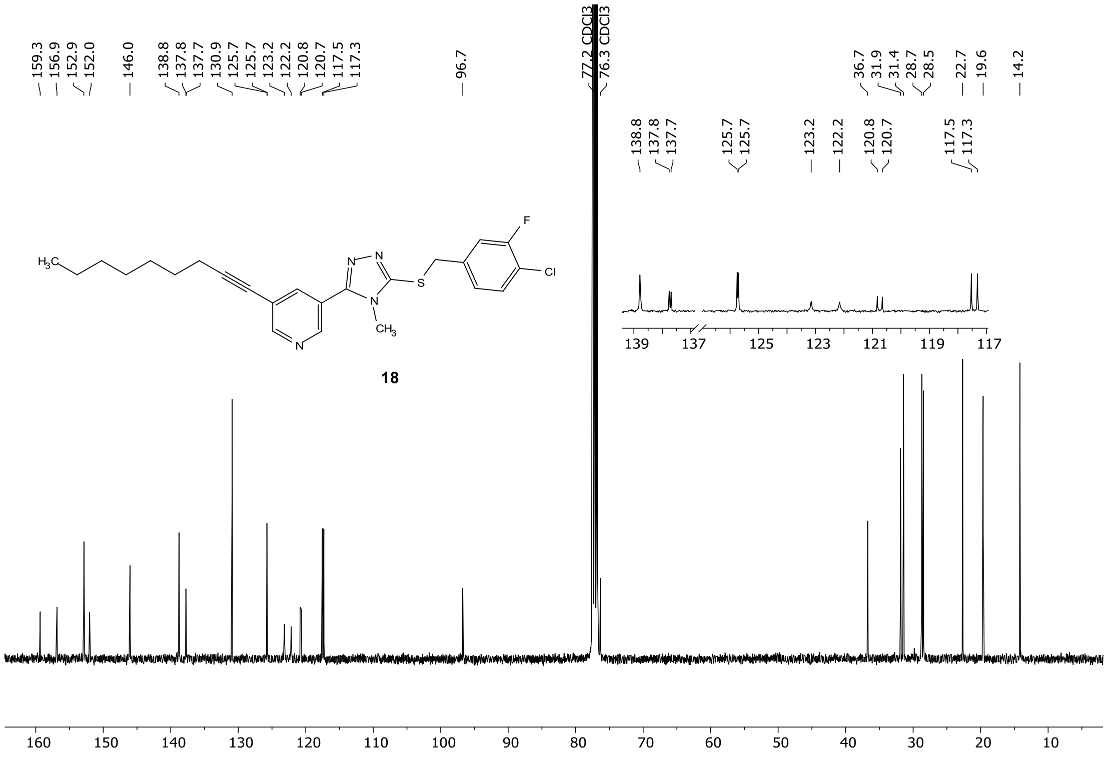

The $^{13}C$ NMR spectrum of compound **18**

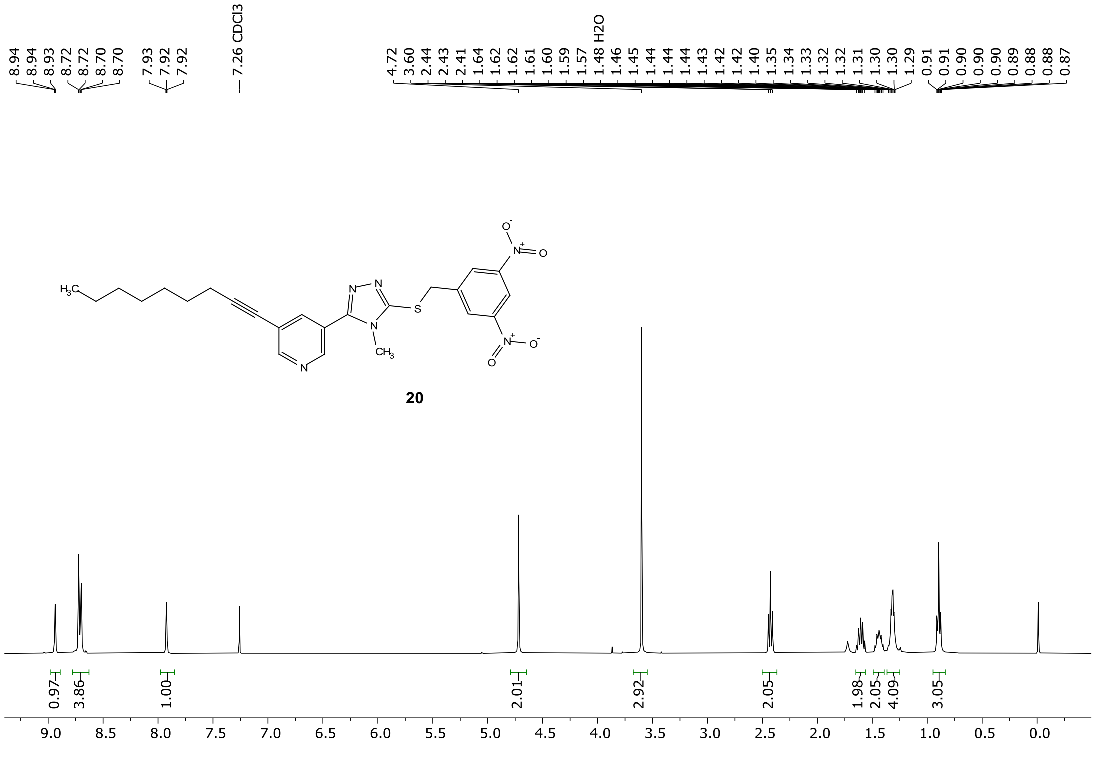

The $^{1}H$ NMR spectrum of compound **20**

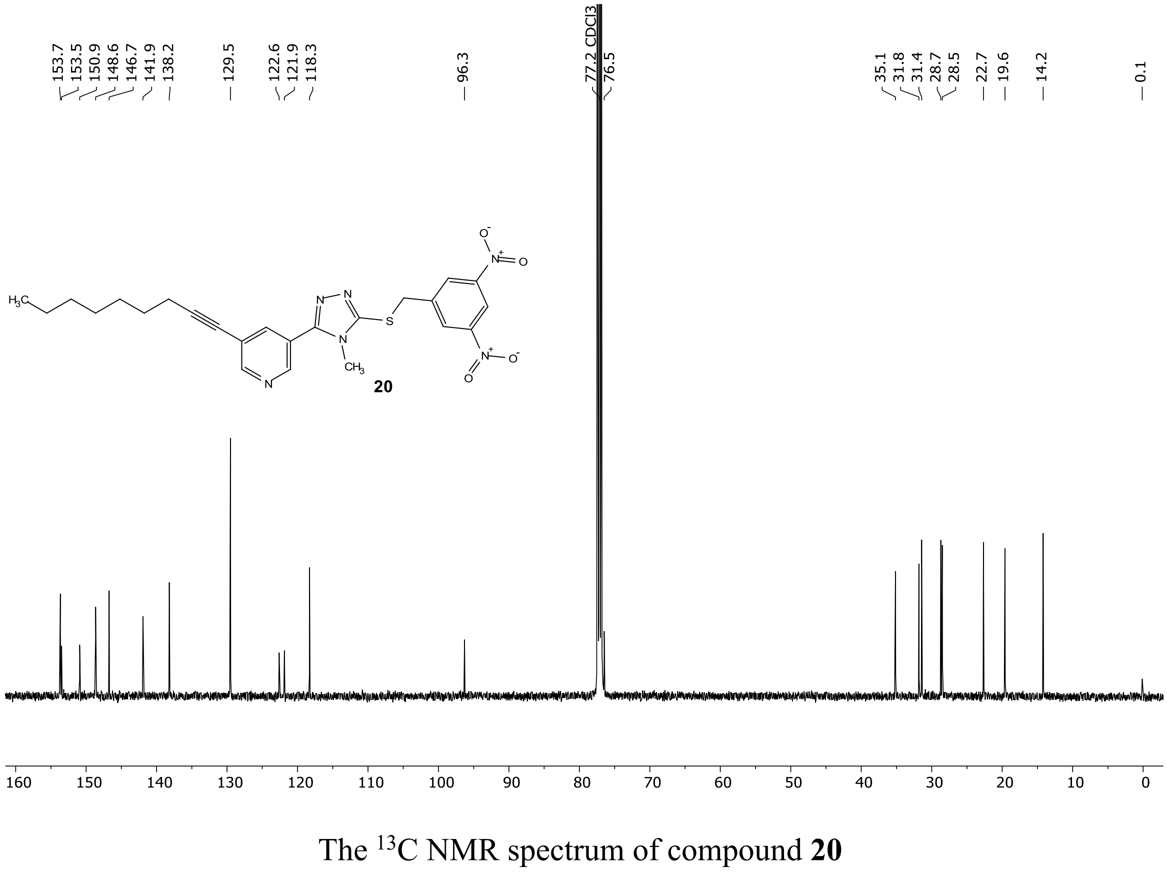

The $^{13}C$ NMR spectrum of compound **20**

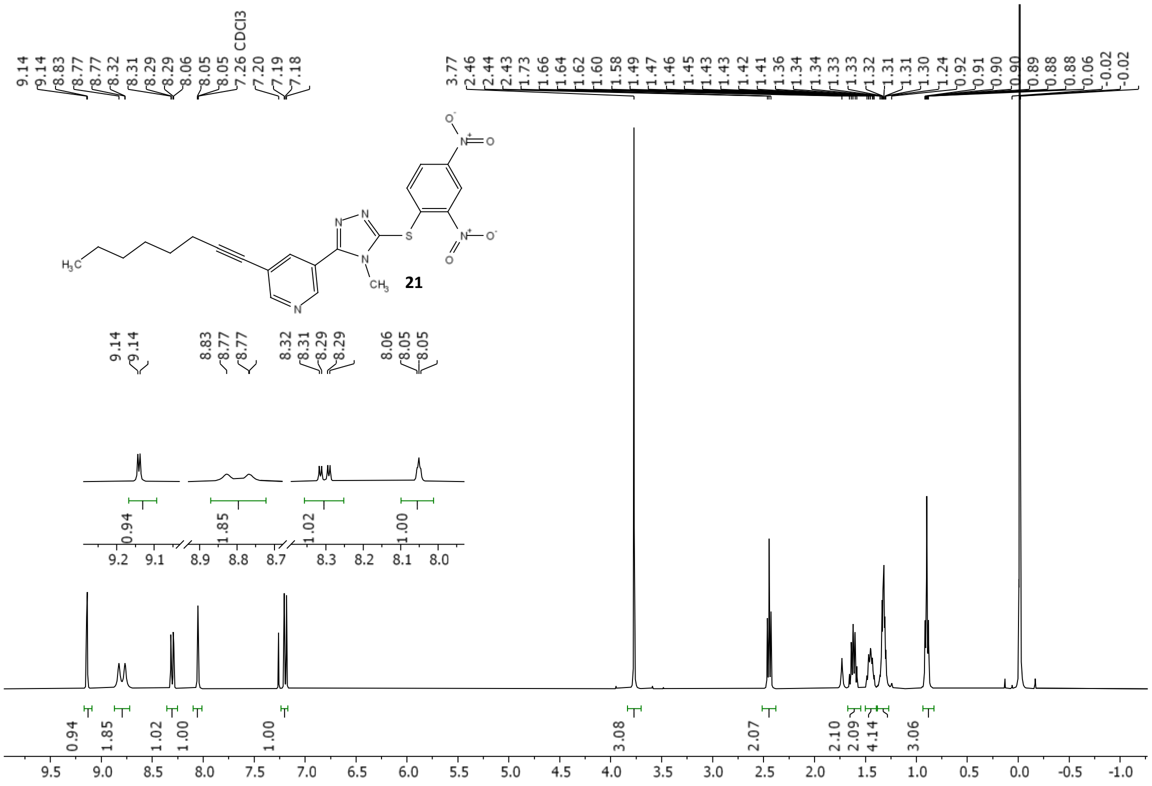

The [1]H NMR spectrum of compound **21**

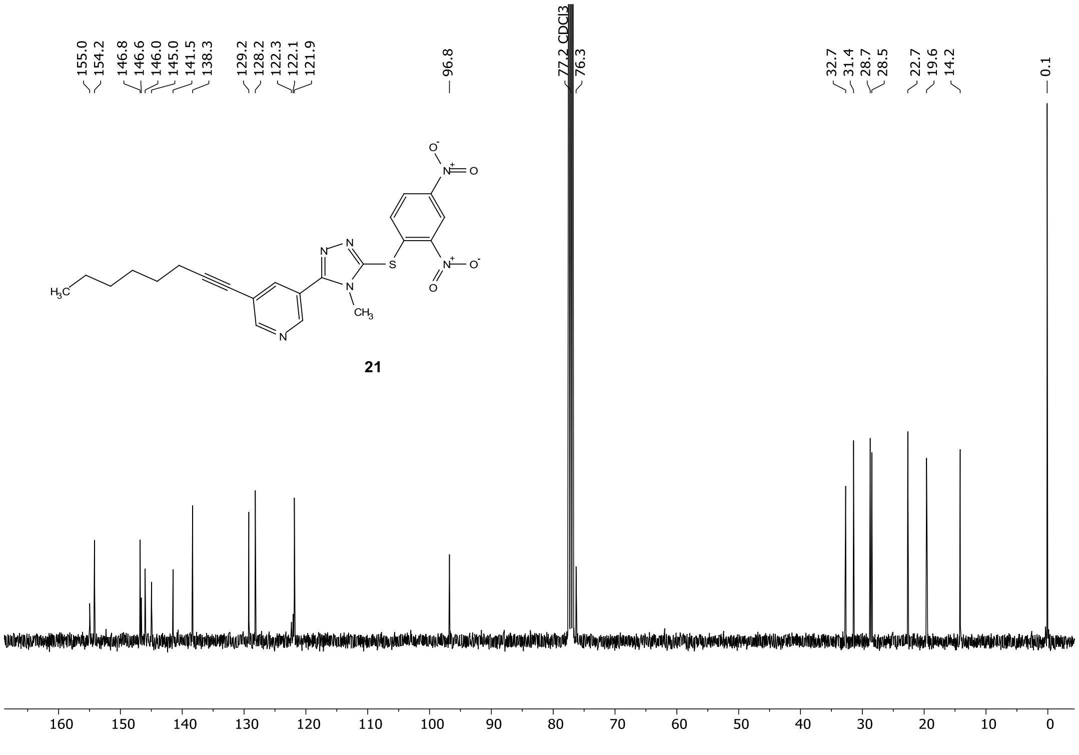

The [13]C NMR spectrum of compound **21**

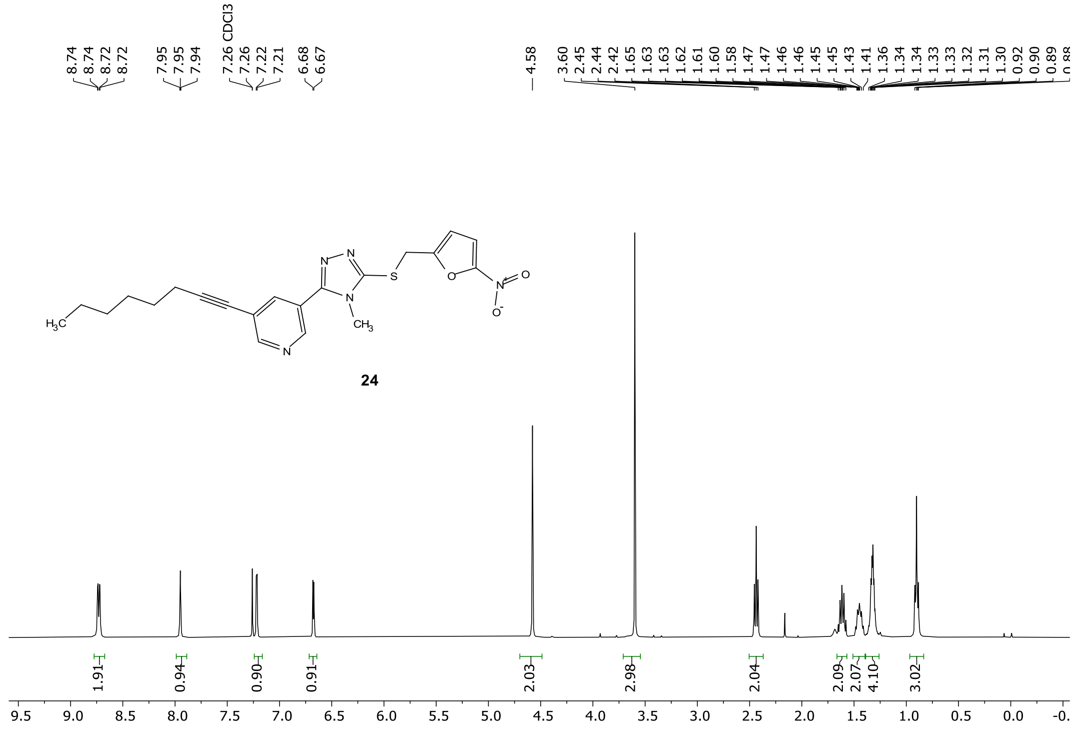

The $^{1}H$ NMR spectrum of compound **24**

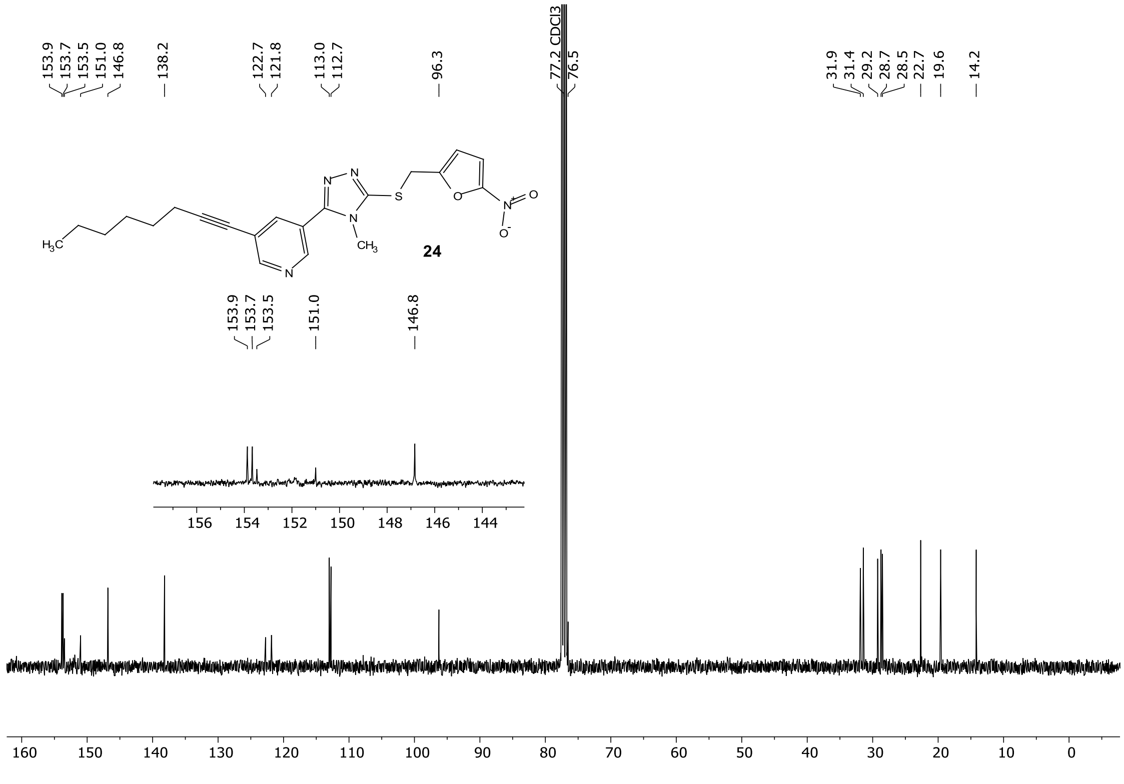

The $^{13}C$ NMR spectrum of compound **24**

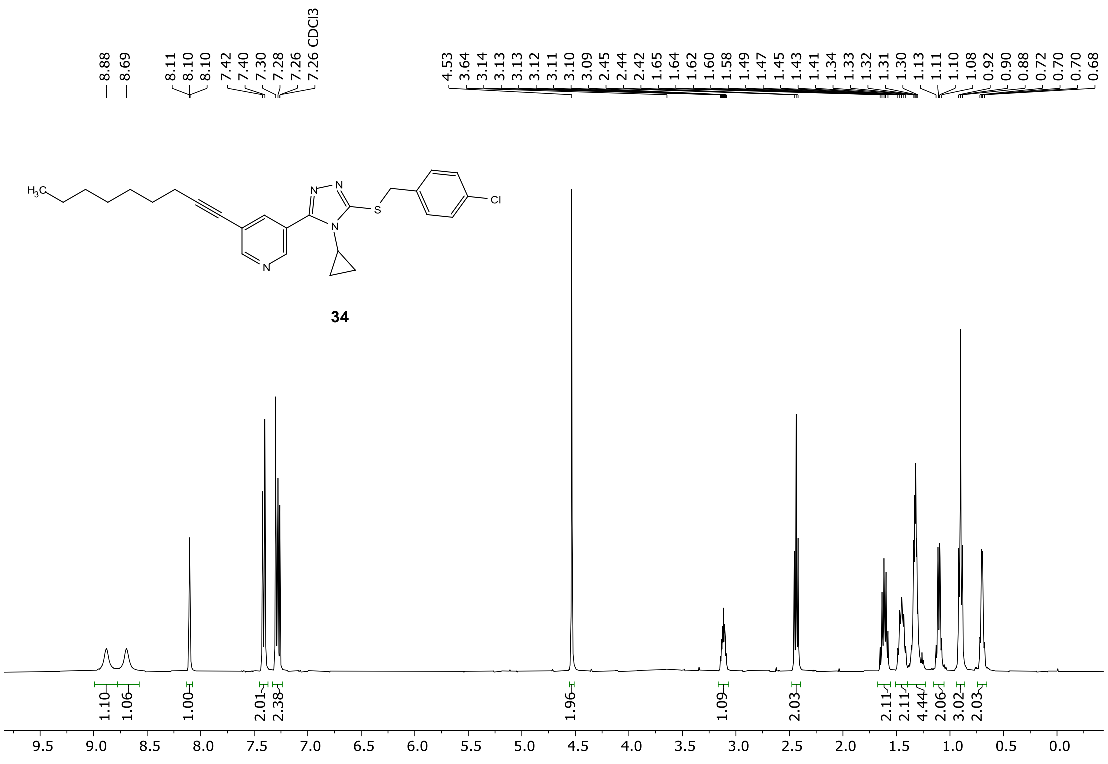

The $^{1}$H NMR spectrum of compound **34**

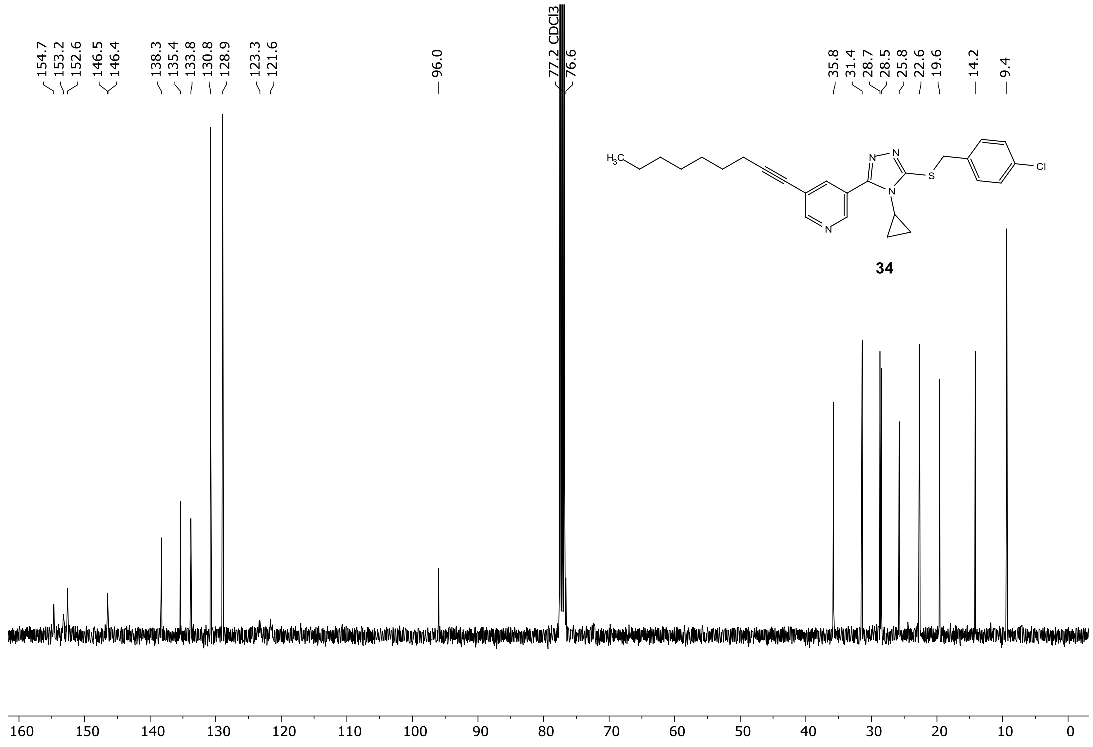

The $^{13}$C NMR spectrum of compound **34**

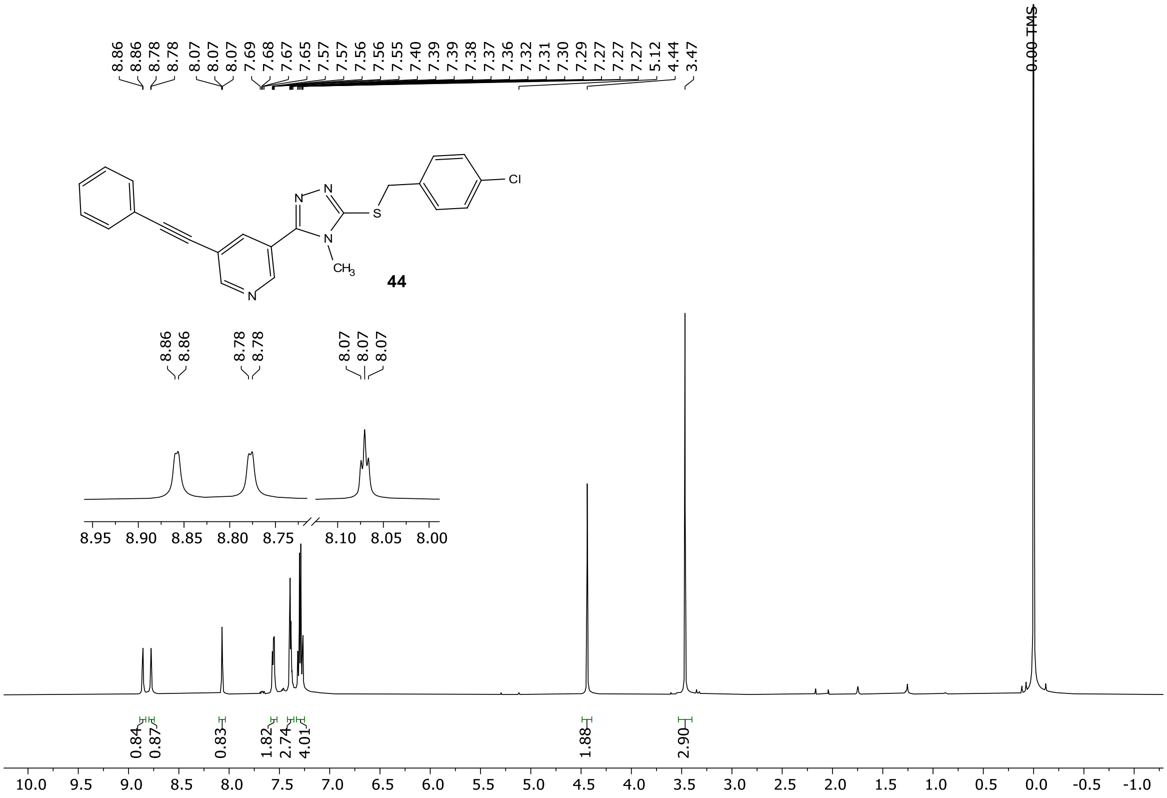

The [1]H NMR spectrum of compound **44**

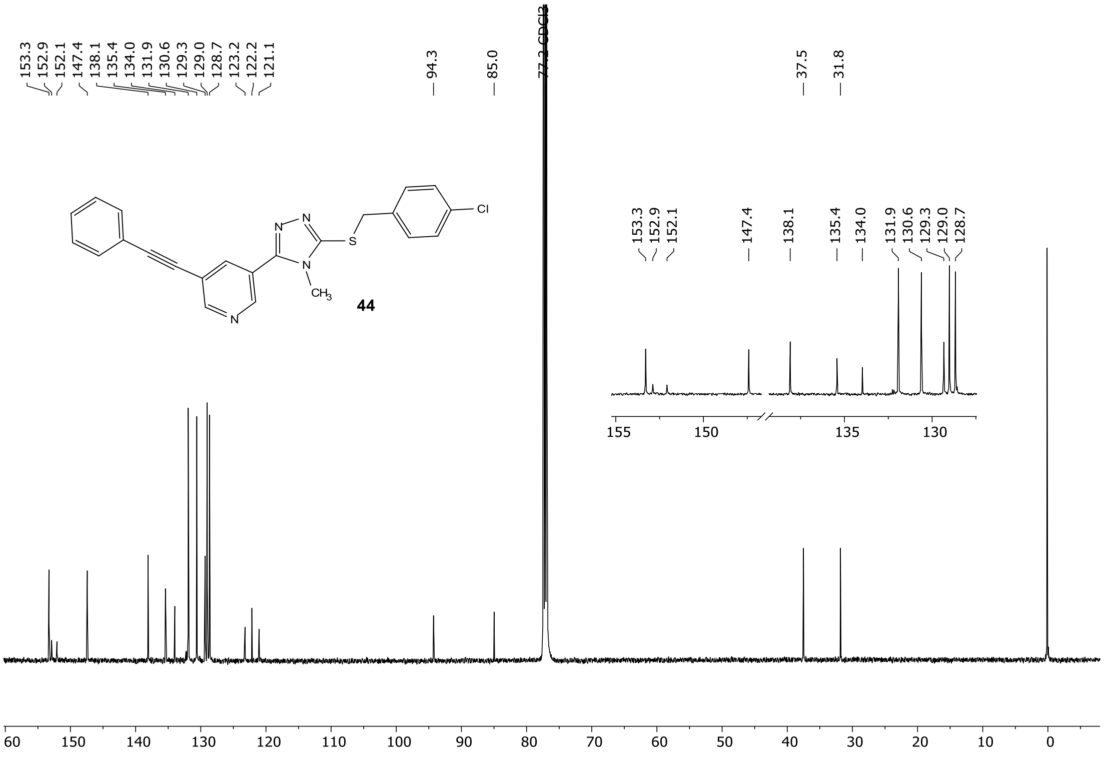

The [13]C NMR spectrum of compound **44**

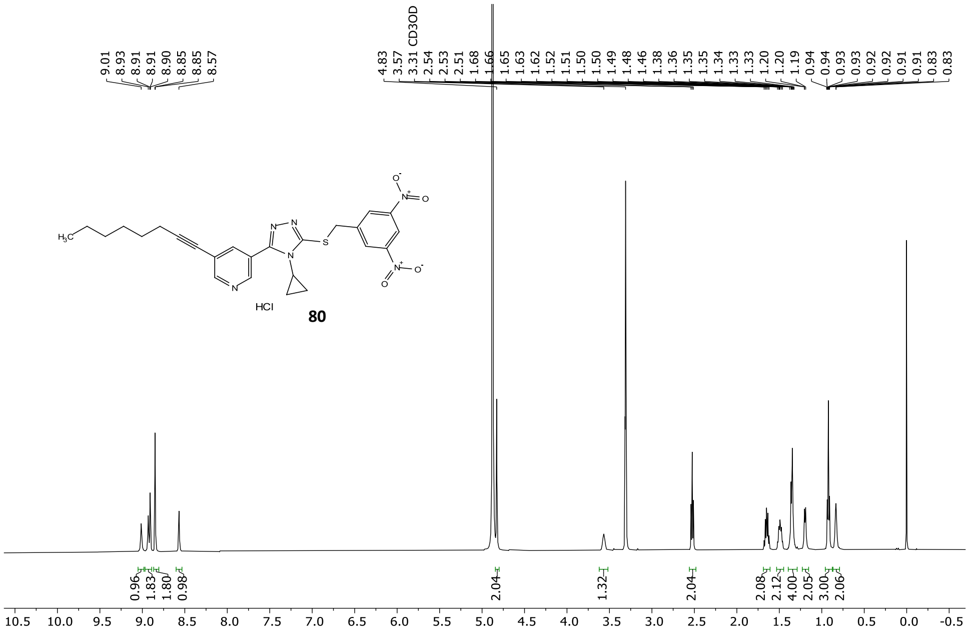

The $^{13}$C NMR spectrum of compound **80**

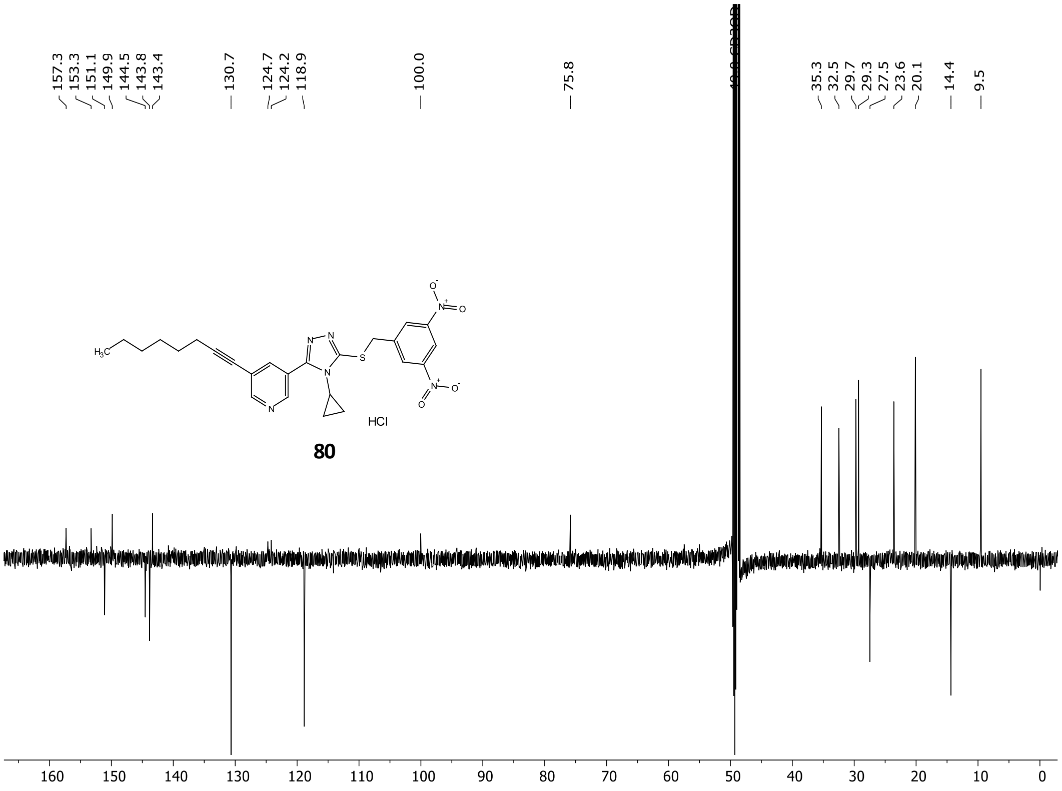

The $^{13}$C NMR spectrum of compound **80**

### 4. HPLC purity analysis

#### 4.1 Procedure for the determination of Purity

The samples were dissolved in 50% acetonitrile and separated on a capillary C18 column C18 column (0.3 mm × 150 mm, 2 μm, Thermo Fisher Scientific) connected with a Dionex Ultimate 3000 UPLC system (Thermo Fisher Scientific). The gradient consisted of HPLC grade water (A) and HPLC grade acetonitrile (ACN). The samples were loaded on to column and the flow rate was 6 μl/min. The gradient consisted of 50-98% ACN over 13 min, hold at 98% ACN for 10 min, return to 50% ACN over 2 min and hold for 20 min. The eluting sample's signal was monitored by UV absorbance at 254 nm. The purity of the sample was calculated by the using Chromeleon 7.3, supplied with the Dionex UPLC system. The peaks were selected manually and its area as well as relative peak area were provided by the Chromeleon. All the compounds that were run had >95% purity.

#### 4.2 HPLC trace of selected compounds ((1, 8, 9, 18, 20, 21, 24, 34, 36, 44, 80)

Compound **1**

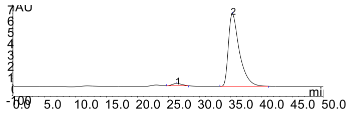

| Peak | Retention time (min)Time | Purity% | Peak Area |
|---|---|---|---|
| 1 | 26.355 | 2.59 | 34.9665 |
| 2 | 35.322 | 97.41 | 1316.9135 |

Compound **8** (TiB-02-037/VS12-12)

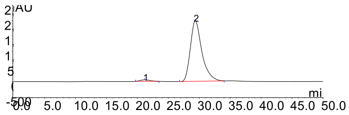

| Peak | Retention time (min)Time | Purity% | Peak Area |
| --- | --- | --- | --- |
| 1 | 21.192 | 2.12 | 92.8951 |
| 2 | 29.363 | 97.88 | 4286.2408 |

Compound **9** (TiB-02-068/VS12-15)

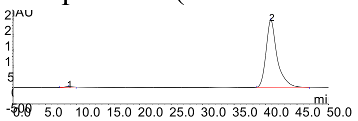

| Peak | Retention time (min)Time | Purity% | Peak Area |
| --- | --- | --- | --- |
| 1 | 8.685 | 0.68 | 30.068 |
| 2 | 40.833 | 99.32 | 4405.953 |

Compound **18** (TiB-02-153/VS12-30)

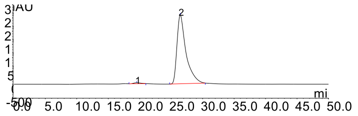

| Peak | Retention time (min)Time | Purity% | Peak Area |
| --- | --- | --- | --- |
| 1 | 19.538 | 1.04 | 45.4767 |
| 2 | 26.445 | 98.96 | 4310.7589 |

Compound **20**

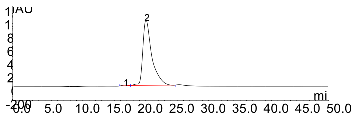

| Peak | Retention time (min)Time | Purity% | Peak Area |
|---|---|---|---|
| 1 | 17.712 | 0.6 | 9.8147 |
| 2 | 21.042 | 99.4 | 1629.8044 |

Compound **21**

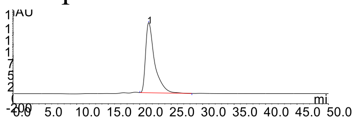

| Peak | Retention time (min)Time | Purity% | Peak Area |
|---|---|---|---|
| 1 | 21.432 | 100 | 2503.1583 |

Compound **24**

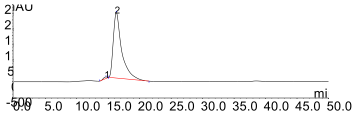

| Peak | Retention time (min)Time | Purity% | Peak Area |
|---|---|---|---|
| 1 | 17.575 | 0.72 | 8.5887 |
| 2 | 19.477 | 1.7 | 20.3325 |
| 3 | 26.29 | 0.62 | 7.3661 |
| 4 | 35.092 | 96.96 | 1156.2514 |

Compound **34**

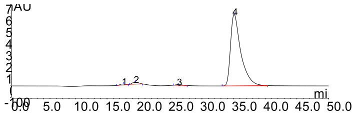

| Peak | Retention time (min)Time | Purity% | Peak Area |
|---|---|---|---|
| 1 | 17.575 | 0.72 | 8.5887 |
| 2 | 19.477 | 1.7 | 20.3325 |
| 3 | 26.29 | 0.62 | 7.3661 |
| 4 | 35.092 | 96.96 | 1156.2514 |

Compound **36**

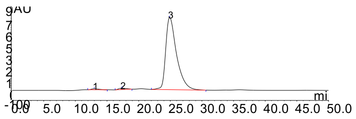

| Peak | Retention time (min)Time | Purity% | Peak Area |
|---|---|---|---|
| 1 | 12.995 | 0.92 | 14.5525 |
| 2 | 17.367 | 1.32 | 20.8362 |
| 3 | 24.905 | 97.76 | 1545.617 |

Compound **44**

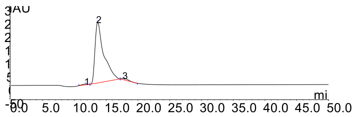

| Peak | Retention time (min)Time | Purity% | Peak Area |
|---|---|---|---|
| 1 | 11.798 | 0.94 | 3.5629 |
| 2 | 13.668 | 97.32 | 369.7484 |
| 3 | 17.807 | 1.74 | 6.6223 |

Compound **80**

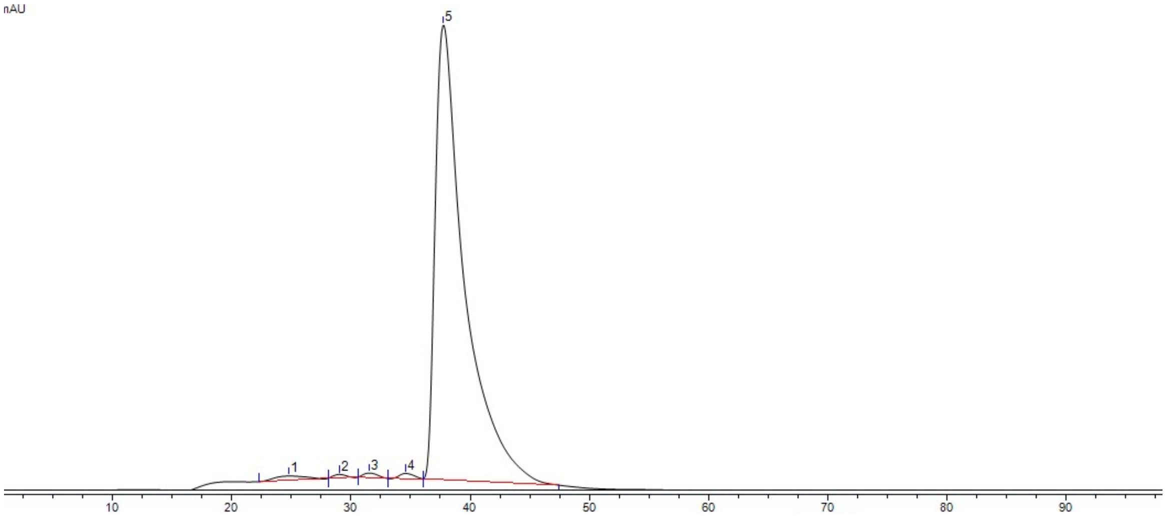

| Peak | Retention time (min) | Purity % | Peak Area |
|---|---|---|---|
| 1 | 24.832 | 1.46 | 4.6509 |
| 2 | 29.183 | 0.9 | 2.8755 |
| 3 | 31.818 | 0.74 | 2.3467 |
| 4 | 34.71 | 0.42 | 1.3362 |
| 5 | 37.538 | 96.49 | 308.0405 |

### 5. References

(1) *Clinical and Laboratory Standards Institute, Methods for Dilution Antimicrobial Susceptibility Tests for Bacteria That Grow Aerobically; Approved Standard 9th Ed., 2012. Document M7-A9.*

(2) Clinical and Laboratory Standards Institute, Reference Method for Broth Dilution Antifungal Susceptibility Testing of Yeasts; Approved Standard 2nd Ed., 2002. Document M27-A2.

(3) Clinical and Laboratory Standards Institute, Reference Method for Broth Dilution Antifungal Susceptibility Testing of Filamentous Fungi; Approved Standard, 2008. Document M38-A2.
